# NAD^+^ activates renal metabolism and protects from chronic kidney disease in a model of Alport syndrome

**DOI:** 10.1101/2024.02.26.580911

**Authors:** Bryce A. Jones, Debora L. Gisch, Komuraiah Myakala, Amber Sadiq, Ying-Hua Cheng, Elizaveta Taranenko, Julia Panov, Kyle Korolowicz, Ricardo Melo Ferreira, Xiaoping Yang, Briana A. Santo, Katherine C. Allen, Teruhiko Yoshida, Xiaoxin X. Wang, Avi Z. Rosenberg, Sanjay Jain, Michael T. Eadon, Moshe Levi

**Author notes:** Corresponding Author: Moshe Levi, Basic Science Building, Room 353 3900 Reservoir Rd NW Washington, DC 20007.

## Abstract

Chronic kidney disease (CKD) is associated with renal metabolic disturbances, including impaired fatty acid oxidation (FAO). Nicotinamide adenine dinucleotide (NAD^+^) is a small molecule that participates in hundreds of metabolism-related reactions. NAD^+^ levels are decreased in CKD, and NAD^+^ supplementation is protective. However, both the mechanism of how NAD^+^ supplementation protects from CKD, as well as the cell types involved, are poorly understood. Using a mouse model of Alport syndrome, we show that nicotinamide riboside (NR), an NAD^+^ precursor, stimulates renal peroxisome proliferator-activated receptor alpha signaling and restores FAO in the proximal tubules, thereby protecting from CKD in both sexes. Bulk RNA-sequencing shows that renal metabolic pathways are impaired in Alport mice and activated by NR in both sexes. These transcriptional changes are confirmed by orthogonal imaging techniques and biochemical assays. Single nuclei RNA-sequencing and spatial transcriptomics, both the first of their kind from Alport mice, show that NAD^+^ supplementation restores FAO in proximal tubule cells. Finally, we also report, for the first time, sex differences at the transcriptional level in this Alport model. In summary, we identify a nephroprotective mechanism of NAD^+^ supplementation in CKD, and we demonstrate that the proximal tubule cells substantially contribute to this benefit.

## INTRODUCTION

Chronic kidney disease (CKD) is a clinical diagnosis characterized by the gradual loss of renal function. The pathophysiology of CKD is complex, and kidney diseases of distinct etiologies can all converge on CKD (1). Nevertheless, several unifying mechanisms emerge when comparing healthy kidneys to their diseased counterparts. These include the progressive worsening of renal fibrosis, inflammation, and metabolic disturbances. This suggests that treatments that prevent these changes might also prevent the associated loss of renal function. Herein, we focus on preventing the progression of kidney disease by activating renal metabolism with the nicotinamide adenine dinucleotide (NAD^+^) precursor, nicotinamide riboside (NR).

Decreased NAD^+^ levels contribute to acute kidney injury (AKI), and NAD^+^ supplementation is protective in models of AKI (2–4). Using both gain- and loss-of-function transgenic mice, Tran et al. (5) comprehensively showed that the peroxisome proliferator-activated receptor-γ coactivator 1-α (PGC-1α) protects from AKI by upregulating genes in the *de novo* NAD^+^ synthesis pathway in the renal tubules, and these effects are replicated by NAD^+^ supplementation. PGC-1α is a coactivator that controls metabolism-related gene regulatory networks, including by direct interaction with the peroxisome proliferator-activated receptor α (PPARα) (6). Their data clearly indicate that improvements in renal tubular mitochondrial function, including FAO, substantially contribute to the protective effects of NAD^+^ supplementation in AKI (5).

Recent studies suggest that reduced NAD^+^ may also play a role in CKD. Endogenous NAD^+^ biosynthesis is impaired in CKD, and there is a corresponding decrease in levels of NAD^+^ and its related metabolites (7, 8). Consistent with this, both promoting NAD^+^ salvage and preventing NAD^+^ breakdown protect from CKD (9, 10). Furthermore, pharmacological NAD^+^-supplementation has generally been shown to protect against CKD (11–16). However, unlike in AKI, the mechanism of how NAD^+^ supplementation protects the kidney in CKD is still poorly understood. A key limitation is that no study thus far has investigated the cell type specific effects of exogenous NAD^+^ supplementation in a model of CKD, which may affect the podocytes and/or the renal tubules to varying extents.

In a recent study, we demonstrated that NR treatment of diabetic mice restored mitochondrial function, including sirtuin 3 activation, thereby preventing mitochondrial damage. This reduced expression of the cyclic GMP–AMP synthase–stimulator of interferon genes (cGAS-STING) pathway and thus protected the kidney. Importantly, we also identified changes in metabolic genes, including NR-mediated increases in mRNA transcripts for *Ppargc1a*, *Nrf1*, *Tfam*, *Cpt1a*, *Acadm*, and *Acadl* (15). These genes are encoded in the nuclear genome, and thus it is unlikely that the changes were a direct result of mitochondrial sirtuin 3. Instead, it implies a mechanism of NAD^+^ supplementation that acts at the transcriptional level to modulate expression of key metabolic genes. We therefore sought to identify the molecular mechanism underlying this change.

To begin to dissect this mechanism, as well as cell types responsible, we returned to the NAD^+^ supplementation data reported from models of AKI. We found the data reported by Tran et al. (5) to be very compelling evidence for the role of PGC-1α in the renal tubules, and it was consistent with the transcriptional activation of metabolic genes that we observed in our recent study (15). Furthermore, mitochondrial dysfunction and associated FAO defects are key metabolic disturbances that drive CKD (17, 18), and restoring renal metabolism has been shown to be protective (19). Given this rationale, we hypothesized that NAD^+^supplementation with NR protects from CKD by activating renal metabolism in the proximal tubules.

We tested this hypothesis with a three-step sequential approach. We first tested NR in a mouse model of Alport syndrome to show that NAD^+^ supplementation protects against CKD at multiple timepoints. We then employed biochemical techniques, including bulk RNA-sequencing (RNA-seq) and immunoblotting, to assess metabolic dysregulation. Finally, we performed single-nuclei RNA-seq (snRNA-seq) and spatial transcriptomics to show that NAD^+^ supplementation enhances renal metabolism in the proximal tubules. The results presented herein provide strong evidence that NR activates the NAD^+^–PGC-1α–PPARα–FAO axis in the proximal tubules, thereby stimulating metabolism and protecting the kidney.

## MATERIALS AND METHODS

### Animal models

Animal studies were approved by the Institutional Animal Care and Use Committee of Georgetown University and adhered to standards set by the Public Health Service Policy on Humane Care and Use of Laboratory Animals. *Col4a3*^tm1Dec^ mice on the C57BL/6J background slowly develop kidney disease, and they were obtained as a gift from Sanofi (Framingham, MA) (20, 21). Col4a3^-/-^ was the disease genotype, and Col4a3^+/-^ was the control genotype. Genotyping was performed by Transnetyx (Cordova, TN) using RT-qPCR.

For all studies, mice were housed at ambient temperature with a 12:12-hr light-dark cycle and free access to food and water. Litter-to-litter variation was controlled by balancing treatment groups across litters. Nicotinamide riboside (CAS No. 1341-23-7) was obtained from ChromaDex (Los Angeles, CA) through participation in their External Research Program. Heparinized plasma and organs were collected upon euthanasia with carbon dioxide.

### Sex as a biological variable

All experiments were conducted in male and female mice until we established a solid rationale for only studying a single sex. This rationale is comprehensively addressed in the Results section.

### NAD^+^ quantification in control and Alport mice

Control and Alport mice were maintained on a grain-based chow (Cat. No. 3005740-220, LabDiet, St. Louis, MO) and euthanized at 25 weeks old.

### NAD supplementation experiment in control and Alport mice

Control and Alport mice were maintained on a grain-based chow (Cat. No. 3005740-220, LabDiet) until 10 weeks old. They were then switched to a purified control diet alone (Cat. No. TD.130352, Envigo, Indianapolis, IN) or admixed with NR (0.5% w/w, Cat. No. TD.190868, Envigo) for the remainder of the study. Administration of this dose of NR to mice via the diet has been reported previously (15). Photoplethysmography, echocardiography, and 24-hr urine collection were performed at 24-25 weeks old. Mice were euthanized at 25 weeks old.

### Replication of NAD supplementation experiment in Alport mice

Control and Alport mice were maintained on a grain-based chow (Cat. No. 3005740-220, LabDiet). NR was administered in the drinking water (5 g/L) to one half of the mice, starting at six-weeks of age. NR water was replaced twice per week. NR is stable in water dispensers for at least six days, and it does not affect water intake (22). Twenty-four–hour urine collection was performed at 24-25 weeks old, and mice were euthanized at 35 weeks old.

### In vivo measurements

Urine was collected from mice by housing them in metabolic cages. Mice were habituated to the cages for one day prior to urine collection. Echocardiography (Vevo 3100, VisualSonics, Toronto, Canada) was performed by an experienced preclinical ultrasound technician as previously described (23) and in accordance with recent guidelines (24). Blood pressure was measured by tail photoplethysmography (Model BP-2000-M-6, Visitech Systems, Apex, NC). Habituation cycles were performed for the two days prior to collection of the blood pressure data.

### Biochemical assays

NAD^+^ was quantified in a mid-transverse kidney piece (Cat. No. E2ND-100, BioAssay Systems, Hayward, CA). Urine albumin (Cat. No. 1011, Ethos Biosciences, Logan Township, NJ) and plasma creatinine (Cat. No. DICT-500, BioAssay Systems) were determined according to the manufacturers’ instructions. Expression of proteins of interest were quantified from a mid-transverse kidney piece by immunoblotting and normalized to total protein using Ponceau S as previously described (25). The primary antibodies used for immunoblotting were CPT1α (Cat. No. ab128568, Abcam, Cambridge, United Kingdom), fibronectin (Cat. No. F3648, MilliporeSigma, Burlington, MA), MCAD (Santa Cruz Biotechnology, Dallas, TX), and PGC-1α (Cat. No. AB3242, MilliporeSigma). The secondary antibodies were anti-rabbit (MilliporeSigma) and light chain-specific anti-mouse (Cat. No. AP200P, MilliporeSigma).

### Histopathology and immunohistochemistry

Tissues were drop-fixed in 10% neutral buffered formalin for 24-hours at 4 °C, dehydrated with an ethanol-xylene gradient, and embedded in paraffin. Formalin-fixed, paraffin-embedded tissues were sectioned (3 µm) onto glass slides using a microtome. Picrosirius red (PSR) staining was performed (Cat. No. SO-674, Rowley Biochemical, Danvers, MA), and polarized images were analyzed by thresholding in ImageJ2 (26). Renal cortical tubulointerstitial fibrosis was quantified using unpolarized PSR images after excluding glomerular, vascular, and medullary contributions in QuPath (27). PSR data are presented as the percent of pixels that stained positive. Immunohistochemistry for p57^kip2^ (Cat. No. ab75975, Abcam) followed by periodic acid–Schiff (Cat. No. 22-110-645, Fisher Scientific, Hampton, NH) post-staining without hematoxylin counterstaining was performed and analyzed as previously described (28). Mesangial index was quantified using QuPath (27). Immunohistochemistry for CD45 (Cat. No. 65087-1-Ig, Proteintech, Rosemont, IL) was performed as previously described (25). Polarized images were acquired with an IX83 Inverted Microscope (Olympus Scientific Solutions, Tokyo, Japan). Unpolarized brightfield images were acquired with an Aperio GT 450 (Leica Microsystems, Wetzlar, Germany) and a Hamamatsu NanoZoomer (Hamamatsu Photonics, Hamamatsu City, Japan).

### Analysis of previously published RNA-sequencing datasets

RNA-seq data from previously published RNA-seq experiments were downloaded from the Sequence Read Archive (29). Accession numbers SRR1611815, SRR1611816, SRR1611817, SRR1611818, SRR1611819, SRR1611820, and SRR1611821 correspond to 3 male control and 4 male Alport kidneys, respectively, on the 129X1/SvJ background, 5.5 weeks of age (30). Accession numbers SRR1611806, SRR1611807, SRR1611808, SRR1611809, SRR1611810, and SRR1611811 correspond to 3 male control and 3 male Alport kidneys, respectively, on the 129X1/SvJ background, 9 weeks of age (30). Accession numbers SRR15102716, SRR15102717, SRR15102718, SRR15102719, SRR15102720, SRR15102721, SRR15102722, SRR15102723, SRR15102724, SRR15102725, and SRR15102726 correspond to 5 male control and 6 male Alport kidneys, respectively, on an F1 mixed 129/SvJ and C57BL/6J background, 15 weeks of age (31). Sequencing files were aligned and processed with BioJupies, a webserver that automatically analyzes RNA-seq datasets and generates Jupyter Notebooks (32). Gene ontology (GO) enrichment analyses and Kyoto Encyclopedia of Genes and Genomes (KEGG) pathway analyses obtained from BioJupies were plotted in GraphPad Prism (San Diego, CA) (33, 34).

### Bulk RNA-sequencing and data analyses

Bulk RNA-seq was performed on isolated kidney cortex from 25-week-old *Col4a3*^tm1Dec^ mice on the C57BL/6J background. All eight combinations of experimental groups were investigated: male versus female, control versus Alport, and vehicle versus NR (*N* = 4 mice per group, 32 mice total). Total RNA was extracted with spin columns (Cat. No. 74104, Qiagen, Germantown, MD), and RNA-seq using the poly(A) selection method was performed by Genewiz (South Plainfield, NJ). Sequencing files were aligned, processes, and analyzed with BioJupies (32). GO enrichment analyses, KEGG pathway analyses, and transcription factor enrichment analyses (33–35) on the top 500 upregulated and downregulated genes were downloaded from BioJupies and further processed to identify statistically significant changes occurring in *a priori* defined comparisons, as described in the Results.

Genes contributing to pathways that were reduced in Alport mice and restored by NR treatment were visualized by plotting KEGG graphs of the top 500 downregulated genes in vehicle-treated Alport mice (compared to vehicle-treated control mice) and the top 500 upregulated genes in NR-treated Alport mice (compared to vehicle-treated Alport mice). Genes contributing to pathways activated by NR treatment were visualized by plotting KEGG graphs of the top 500 upregulated genes in NR-treated control mice (compared to vehicle-treated control mice) and the top 500 upregulated genes in NR-treated Alport mice (compared to vehicle-treated Alport mice). KEGG graphs were rendered by Pathview Web (36, 37). Each gene contributed only once, and genes (primarily pseudogenes) that did not convert to the Ensembl namespace with gConvert were not plotted (38).

A separate analysis was performed to investigate a potential sex-genotype interaction. The subset of 16 vehicle-treated mice, 4 from each sex/genotype combination, were compared in a two-by-two factorial design using iDEP, a web application for analyzing previously aligned RNA-seq data (39). Differential gene expression between the sexes, the genotypes, and the interaction term were calculated. Genes with both at least a two-fold change in expression and an adjusted *P*-value (FDR) less than 0.05 were deemed statistically significant. All principal component analyses were performed using iDEP and plotted with GraphPad Prism (39).

A separate analysis was performed to investigate a potential effect of mouse genetic background. The subset of 8 vehicle-treated male mice, 4 from each genotype, were compared with control and Alport mice on the 129 and mixed B6/129 backgrounds. A meta-analysis was performed using ExpressAnalyst as previously described (40, 41). Briefly, raw read counts were filtered, Log2 normalized, and batch corrected. *P*-values from genes with adjusted *P*-values (FDR) less than 0.05 in at least 1 study were combined using Fisher’s method. KEGG pathway analyses were performed on genes with at least a two-fold change in expression and an adjusted *P*-value (FDR) less than 0.05 (42).

### Single nuclei RNA-sequencing

Single nuclei RNA-seq and the simultaneous assay for transposase-accessible chromatin were performed on kidneys from 25-week-old *Col4a3*^tm1Dec^ mice on the C57BL/6J background. All eight combinations of experimental groups were investigated: male vs. female, control vs. Alport, and vehicle vs. nicotinamide riboside (*N* = 1 mouse per group, 8 mice total). Male and female nuclei of the same genotype/treatment combination were pooled, and only the pooled snRNA-seq data are presented here.

For snRNA-seq, 49,488 nuclei were isolated from tissue cryosections and processed using the Chromium Next GEM Single Cell Multiome ATAC + Gene Expression (v1.0) kit as previously described (43). The RNA and ATAC libraries were sequenced separately on an Illumina NovaSeq 6000 system (Software v.1.7.0 and v.1.7.5). For RNA analysis, cell barcodes passing the following quality control filters were used for downstream analyses: 1) passed 10X Cell Ranger Arc (RNA) filters; 2) nCount_RNA showing greater than 1,000 and less than 25,000 in non-mitochondrial genes detected; and 3) cells with percentage of mitochondrial transcripts greater than 25 percent were removed. RNA counts were normalized using sctransform, and the clustering was done with 0.5 resolution. This integration was performed using Seurat (v.5.0.0). Cluster annotations are based on published gene markers (44) and converted by BioMart (v.2.56.1) to the mouse homologous genes. The differential expressed genes are performed by RunPresto with logFC threshold = 0, Wilcox test, and corrected by Bonferroni from package SeuratWrappers (v.0.3.2). The pathway analyses were performed by pathfindR (v.2.3.0.9001) (45). The mmu_STRING database to identify relevant networks of protein interactions and mmu_KEGG database to contextualize these networks within known biological pathways are enriched.

### Spatial transcriptomics

Spatial transcriptomics was performed on kidneys from 25-week-old *Col4a3*^tm1Dec^ mice on the C57BL/6J background. All four combinations of male experimental groups were investigated: control versus Alport, and vehicle versus NR (*N* = 1 male mouse per group, 4 mice total). The same tissue blocks were used for both snRNA-seq and spatial transcriptomics.

Murine kidney tissues were embedded in O.C.T. compound (Cat. No. 23-730-571, Fisher Scientific) immediately after euthanasia. Samples were processed according to the Visium Spatial Gene Expression protocol (10x Genomics, CG000240 protocol) (46). One sample from each condition underwent cryosectioning to yield a section of 10 µm thickness. The sections were stained with periodic acid–Schiff and imaged using a Keyence BZ-X810 microscope, equipped with a Nikon 10X CFI Plan Fluor objective lens. The brightfield images were compiled and matched with Visium fiducials to create comprehensive mosaics. The mRNA from the tissue sections was extracted following a 12-minute permeabilization period. This mRNA then adhered to oligonucleotides at the fiducial spots and was subsequently reverse transcribed. In the subsequent stages of library creation and sequencing, the mRNA was converted into second-strand cDNA. This was followed by denaturation, amplification of the cDNA, and purification using SPRIselect cDNA cleanup (Visium CG000239 protocol). Finally, the cDNA sequencing was performed using an Illumina NovaSeq 6000 system. For spatial analysis, Space Ranger (v2.0.0) with the reference mouse genome (mm10-2020-A) was used to perform expression analysis, mapping, counting, and clustering. Labels were transferred from snRNA-seq to ST to spatially localize the cell types based on gene expression profiles using Seurat (v.5.0.0) onto 8,748 spots.

In ST samples, spots with positive expression of *Nphs2* and *Wt1* were annotated as glomeruli. Conversely, spots were annotated as cortical PT when positive expression of *Slc34a1* was observed without expression of *Nphs2* and *Wt1*. The outer stripe of the medulla was selected by the presence of *Slc3a1* expression above *Slc34a1* expression and without *Nphs2* or *Wt1* expression. Each spot could only be assigned to a single functional tissue unit. An investigator confirmed annotations of each spot using histology or excluded spots if the histology was inconsistent with the marker gene expression. A pseudobulk comparison was made across all spots in the sample. Differential expression between the Alport NR and Alport Vehicle spots for each annotation was evaluated with a Mann-Whitney test. Pathway enrichment was performed with pathfinder (31608109) using STRING network and KEGG pathways. For *Cpt1a* and *Acadm*, spots were classified as non-zero if expression was higher than zero, or zero if no expression was detected. An odd’s ratio with 95% confidence interval was used to assess the likelihood of non-zero spots localizing to each region.

### Statistical analyses for kits, immunoblots, and microscopy

One-way analysis of variance (ANOVA) was performed with GraphPad Prism. For ANOVA post-hoc tests, the decision to compare only the following groups was made *a priori*: (1) control + vehicle vs. control + NR, (2) control + vehicle vs. Alport + vehicle, (3) Alport + vehicle vs. Alport + NR, and (4) control + vehicle vs. Alport + NR.

## RESULTS

### Alport mice have reduced NAD^+^ levels and impaired renal metabolism

To confirm that pathways related to both NAD^+^ and renal metabolism are dysregulated in Alport mice, we re-analyzed previously published RNA-seq data from two independent experiments (30, 31). GO enrichment and KEGG pathway analyses were performed on the 500 most down-regulated genes in Alport mice, and both NAD^+^ biosynthetic pathways (**Figs. S1A-B** ) and fatty acid metabolic pathways (**Fig. S2** ) were significantly enriched. This is in stark contrast to the pathways identified from analyses of up-regulated genes, most of which were related to inflammation and fibrosis (**Figs. S1C-D**) (25). Finally, we confirmed that kidney NAD^+^ levels were lower in male and female Alport mice compared to control mice (**Fig. 1**).

**Figure 1:**
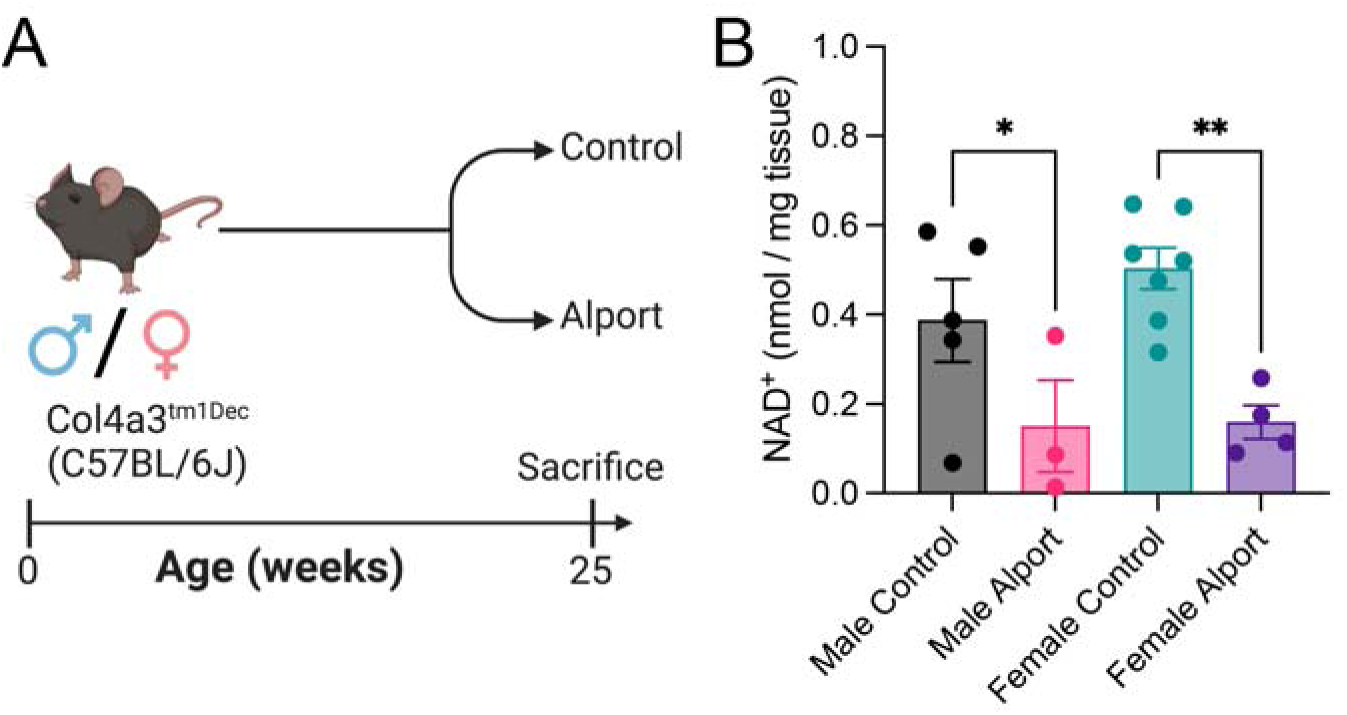
Kidney NAD^+^ is reduced in Alport mice. (**A**) Experimental design: Control and Alport mice of both sexes were sacrificed at 25-weeks of age. (**B**) Alport mice had lower levels of kidney NAD^+^ than control mice. Significance was determined by one-way ANOVA with the Holm–Šídák correction for multiple comparisons. Data are expressed as the means ± SEM, and each datum represents one mouse.

### NAD^+^ supplementation protects Alport mice from kidney disease

Given that kidney NAD^+^ levels were decreased in Alport mice compared to control mice, we hypothesized that NAD^+^ supplementation with NR would reduce the severity of kidney disease. Our colony of Alport mice on the C57BL/6J background slowly develop kidney disease until death at 35-40 weeks of age, and we investigated two timepoints.

In our initial experiment, mice were treated with or without NR between 10 and 25 weeks of age (**Fig. 2A**). We did not observe any changes in echocardiography or blood pressure measurements between the groups, excluding these as potential confounding variables (**Tables S1-2** ). Twenty-four–hour urinary albumin excretion, a marker of kidney damage, was up to 1000-fold increased in Alport mice, and NR treatment prevented this increase in both sexes (**Fig. 2B**). Plasma creatinine was unchanged between control and Alport mice at the 25-week timepoint, consistent with the slowly progressing phenotype of Alport mice on the B6 background (**Table S3**). As shown by polarized microscopy of PSR-stained kidney sections, a technique that is highly specific for collagen (47), NR treatment prevented the progression of overall renal fibrosis in both sexes (**Figs. 2C-D** ). In addition, renal cortical tubulointerstial fibrosis—quantified by excluding medullary, vascular bundle, and glomerular contributions—was increased in Alport mice and reduced by NR treatment in both sexes (**Figs. S3A-B** ). Renal inflammatory infiltrate was also increased in Alport mice and reduced by NR treatment (**Figs. S3C-D**). Finally, the prevention of renal fibrosis was secondarily confirmed by immunoblotting for fibronectin which was reduced in NR-treated male (significant, *P* < .001) and female (trend, *P* < .10) Alport mice (**Fig. S4**).

**Figure 2:**
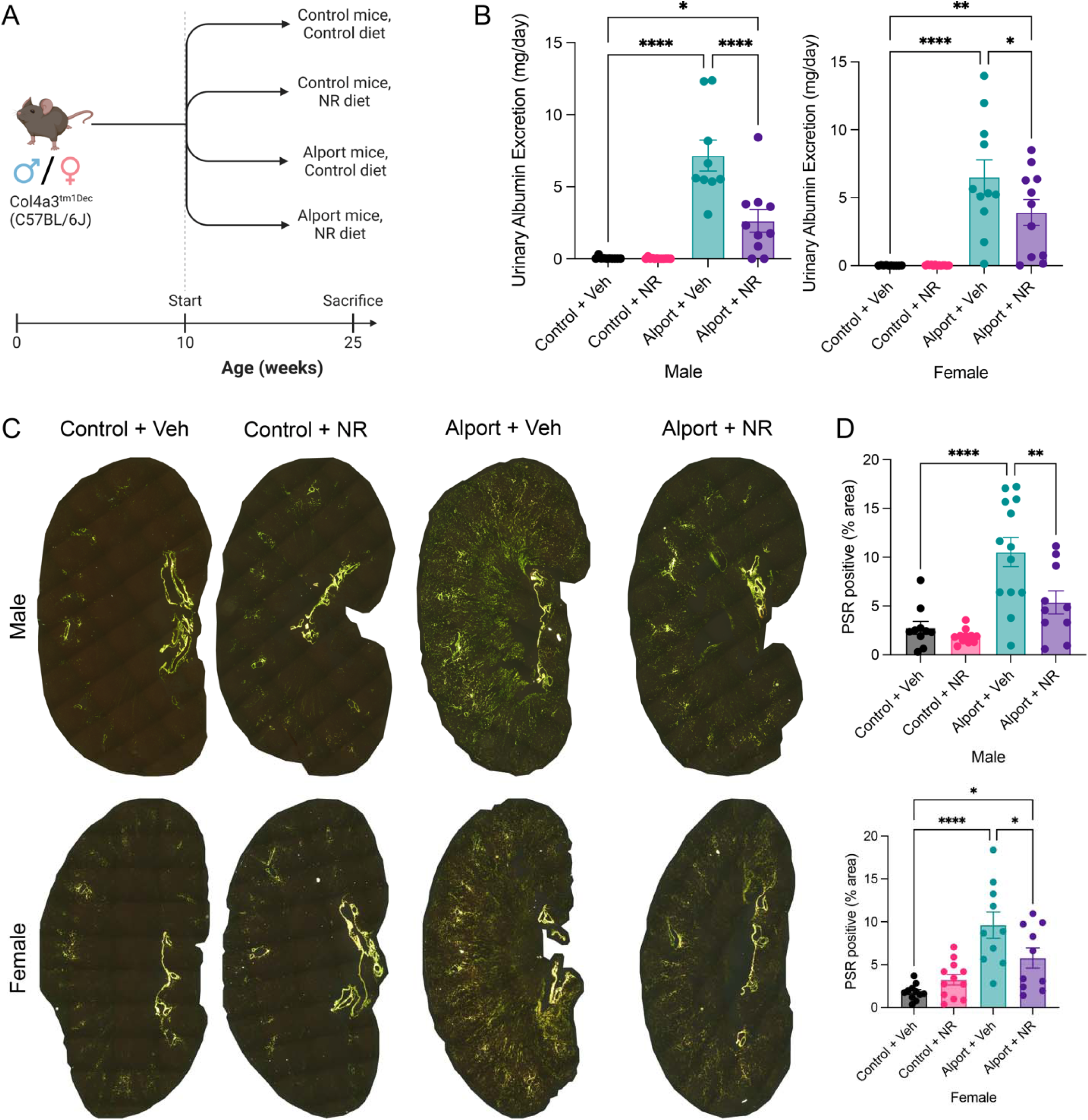
NAD^+^ supplementation protects the kidney in Alport mice. (**A**) Experimental design: Control and Alport mice of both sexes were treated with or without NR between 10-weeks and 25-weeks of age. (**B**) NR treatment reduced 24-hour urinary albumin excretion in Alport mice of both sexes. (**C**) Representative images of PSR-stained kidneys acquired with polarized light. Yellow-green-orange birefringence is highly specific for fibrosis. (**D**) Quantification of PSR-stained kidneys shows that NR treatment reduced renal fibrosis in both sexes. Significance was determined by one-way ANOVA with the Holm–Šídák correction for multiple comparisons. Data are expressed as the means ± SEM, and each datum represents one mouse. \**P* < 0.05, \*\**P* < 0.01, \*\*\*\**P* < 0.0001. NR, nicotinamide riboside; PSR, picrosirius red; Veh, vehicle.

We then repeated the experiment in both male and female mice, although we aged the mice longer to an average of 35 weeks (**Fig. S5A**). Twenty-four–hour urinary albumin excretion was increased in Alport mice, and NR treatment ameliorated this increase in both sexes (**Fig. S5B**). At this later timepoint, plasma creatinine was greatly increased in Alport mice compared to control mice, and NR treatment prevented this increase in both sexes (**Fig. S5C** ). Both our initial and replication experiments had substantial numbers of littermate-matched mice in each group, and they were temporally separated by greater than one year.

### NAD^+^ supplementation protects from podocyte and tubular injury in Alport mice

Glomerular damage was further assessed by immunostaining for the podocyte marker p57^kip2^. Volumetric podocyte density is a podometric that controls for the thickness of the histological section, the size of the podocyte nucleus, and the size of the glomerulus (28, 48, 49). Compared to control mice, Alport mice had reduced glomerular volumetric podocyte density, both podocyte and glomerular hypertrophy, and an increased mesangial index. NR treatment prevented these pathologic changes in both sexes (**Figs. 3A-D** and **S6A**). In females, but not males, the corrected podocyte number per glomerulus was reduced in Alport mice and restored with NR treatment (**Fig. S6B** ). However, unlike the volumetric podocyte density, the corrected podocyte number per glomerulus does not control for glomerular hypertrophy and should be interpreted with caution. These results, in combination with the reduction in urinary albumin excretion, demonstrate that NR treatment protects from glomerular damage in the Alport model of kidney disease.

**Figure 3:**
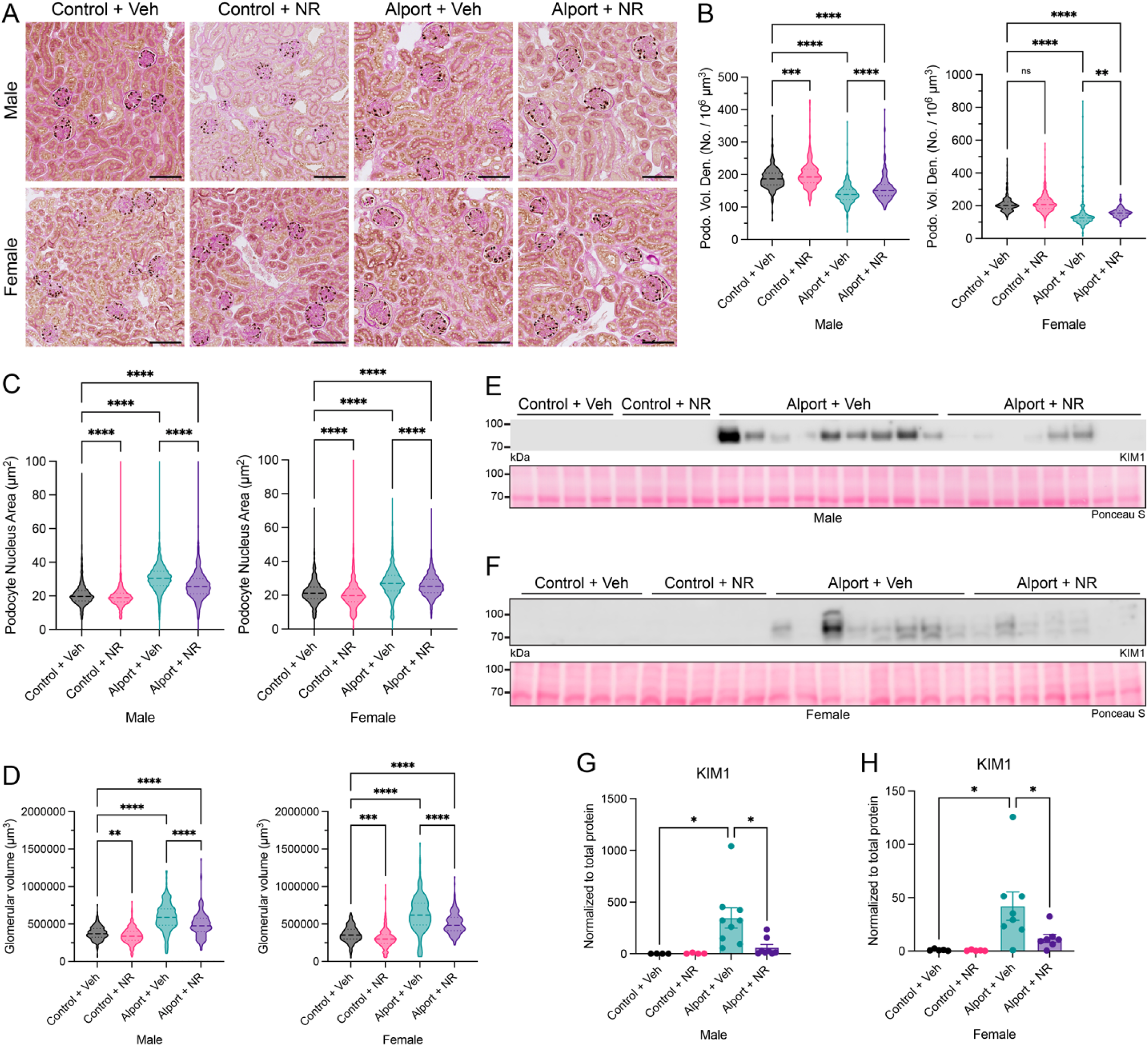
NAD^+^ supplementation prevents both glomerular and tubular injury in Alport mice. (**A**) Representative images of immunohistochemistry for p57^kip2^, followed by periodic acid–Schiff post-staining without hematoxylin counterstaining. Podocyte nuclei are stained brown. (**B-D**) Quantification of p57^kip2^ immunostaining with PodoCount, a validated algorithm to analyze p57^kip2^-stained whole slide images. Podocyte volumetric density (*B*) was reduced in Alport mice and restored by NR treatment in both sexes. Alport mice had podocyte nuclear hypertrophy (*C*) and glomerular hypertrophy (*D*) that was reduced by NR treatment in both sexes. (**E,F**) Immunoblots for KIM-1, a tubular injury marker, in male (*E*) and female (*F*) kidney homogenate. Ponceau S, a nonspecific protein stain, was used as a loading control. (**G,H**) Quantification of immunoblots (*E,F*) shows that KIM-1 is increased in Alport mice and reduced by NR treatment in male (*G*) and female (*H*) mice. Scale bars represent 100 µm. Significance was determined by one-way ANOVA with the Holm–Šídák correction for multiple comparisons. Data are expressed as the means ± SEM. Each datum represents one glomerulus (*B-D*) or one mouse (*G-H*). \**P* < 0.05, \*\**P* < 0.01, \*\*\**P* < 0.001, \*\*\*\**P* < 0.0001. Kidney injury molecule-1, KIM-1; NR, nicotinamide riboside; PSR, picrosirius red; Veh, vehicle.

Although Alport syndrome is classically associated with a specific glomerular defect, glomerular injury can also cause tubular injury (50, 51). In addition, the proximal tubules are highly metabolically active, and they may have limited reserve to cope with increased protein leakage. This might be further exacerbated by an NAD^+^ deficiency. We therefore hypothesized that Alport mice would exhibit substantial tubular pathology that was reversed by NR treatment. Consistent with our hypothesis, renal kidney injury molecule 1 (KIM-1) expression, a marker of tubular damage (52, 53), was increased in Alport mice, and NR treatment prevented this increase in both sexes (**Fig. 3E-H**). We then investigated the mechanism underlying this protective effect.

### Bulk RNA-sequencing identifies renal cortical metabolic defects in Alport mice that are prevented by NAD supplementation

We performed bulk RNA-seq on isolated kidney cortex to both confirm our hypothesis that NAD^+^ supplementation normalizes renal metabolism and to identify if PGC-1α–PPARα signaling is a molecular mechanism driving this change. Principal component analysis demonstrated separation by genotype (PC1, 42.7% of variance, *P* = 2.75×10^-11^), sex (PC2, 11.3% of variance, *P* = 1.16×10^-07^), and treatment (PC3, 7.3% of variance, *P* = 2.07×10^-03^). In both sexes, NR treatment shifted Alport samples towards the control genotype, suggesting the prevention of the disease process (**Fig. S7**).

For each sex, we then performed GO biological process, KEGG pathway, and transcription factor enrichment analyses (33–35) comparing: 1) NR-treated control mice vs. vehicle-treated control mice; 2) vehicle-treated Alport mice vs. vehicle-treated control mice; and 3) NR-treated Alport mice vs. vehicle-treated Alport mice. The decision to compare these groups was made *a priori* as they are the most relevant biological comparisons.

Consistent with our hypothesis, GO enrichment analyses revealed that numerous metabolism-related biological processes were changed between the groups. In both sexes, GO biological processes involving fatty acids, including fatty acid β-oxidation, were among the most highly enriched set of processes that were simultaneously downregulated in vehicle-treated Alport mice (versus vehicle-treated control mice) (**Figs. 4A** and **S8A**) and upregulated in NR-treated Alport mice (versus vehicle-treated Alport mice) (**Figs. 4B** and **S8B**). Many GO biological processes that are integral to energy metabolism were also simultaneously enriched in both comparisons, including acetyl-CoA and acyl-CoA metabolic process (**Figs. 4A-B** and **S8A-B**). KEGG pathway analyses yielded similar results in both sexes (**Figs. 4C-D** and **S8C-D**). The severe impairment of renal metabolism in Alport mice is evident by the downregulation of genes involved in metabolic pathways (KEGG

**Figure 4:**
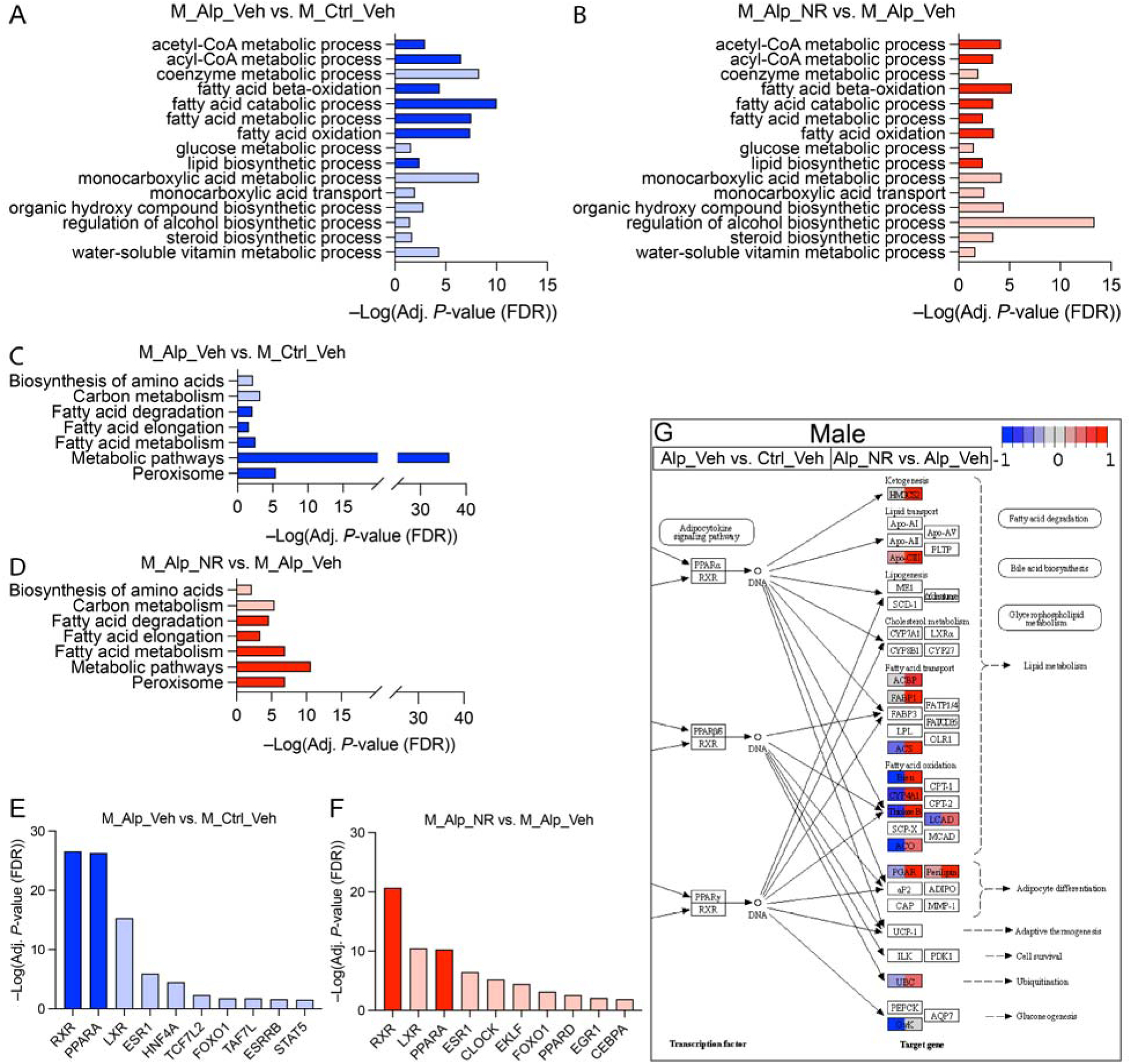
NAD^+^ supplementation activates renal metabolism in male Alport mice. Bulk kidney cortex RNA-seq data from control and Alport mice, treated with or without NR, were analyzed (*N* = 4 mice per group). (**A,B**) Gene ontology (GO) biological processes that are both (*A*) reduced in vehicle-treated male Alport mice (vs. vehicle-treated male control mice) and (*B*) increased in NR-treated male Alport mice (vs. vehicle-treated male Alport mice) are shown. (**C,D**) KEGG pathways that are both (*C*) reduced in vehicle-treated male Alport mice (vs. vehicle-treated male control mice) and (*D*) increased in NR-treated male Alport mice (vs. vehicle-treated male Alport mice) are shown. (**E,F**) Transcription factor analyses suggest that the RXR/PPARα gene regulatory network is (*E*) inhibited in vehicle-treated male Alport mice (vs. vehicle-treated male control mice) and (*F*) activated in NR-treated male Alport mice (vs. vehicle-treated male Alport mice). Processes (*A*,*B*), pathways (*C*,*D*), and transcription factors (*E*,*F*) that are directly involved in fatty acid metabolism are highly enriched in all comparisons, emphasized by either dark blue (decreased) or dark red (increased). (**G**) Partial KEGG graph for the PPAR signaling pathway (KEGG Entry No. 03320). The left and right sides of each gene box represent data from vehicle-treated male Alport mice (vs. vehicle-treated male control mice) and NR-treated male Alport mice (vs. vehicle-treated male Alport mice), respectively. The sub-pathway that was the most restored by NR treatment was fatty acid oxidation. Alp, Alport; Ctrl, control; M, male; NR, nicotinamide riboside; Veh, vehicle.

Entry No. 01100) with an adjusted *P*-value (FDR) approaching forty orders of magnitude in both sexes (**Figs. 4C** and **S8C**). Transcription factor analyses revealed differential activity of the RXR/PPARα heterodimer as a likely candidate underlying the observed transcriptional differences (**Figs. 4E-F** and **S8E-F**).

Based on the data from the GO enrichment, KEGG pathway, and transcription factor analyses, we plotted the gene changes on KEGG graphs for the PPAR signaling pathway (KEGG Entry No. 03320) (**Figs. 4G**, **S8G**, and **S9-10**), the peroxisome (KEGG Entry No. 04146) (**Figs. S11-12** ), and fatty acid degradation (KEGG Entry No. 00071) (**Figs. S13-14**). A multitude of genes involved in these pathways, especially those related to FAO, were reduced in vehicle-treated Alport mice (compared to vehicle-treated control mice) and restored in NR-treated Alport mice (compared to vehicle-treated Alport mice). Data from both comparisons are visualized simultaneously on the KEGG graphs, the former on the left half of each rectangle, and the latter on the right half.

All together, these data represent clear evidence for impaired renal metabolism in Alport mice, including fatty acid metabolism, that is restored by NR treatment. Next, we investigated NR treatment in control mice to confirm that NR is directly responsible for normalizing renal metabolism.

### NAD supplementation activates renal metabolism in control mice, confirming this as a causal mechanism of NR-mediated kidney protection

Although the dramatic change in metabolic state between vehicle- and NR-treated Alport mice suggests that NR protects the kidney by activating renal metabolism, it is not enough alone to show a causal relationship. In other words, because renal metabolism in Alport mice becomes gradually more impaired as kidney disease progresses, any intervention that reduces kidney disease will also cause a coincidental improvement of renal metabolism. This is not because the intervention activates renal metabolism, *per se*. It is instead an artifact that arises because the intervention-treated group has less severe kidney disease than the vehicle-treated group at the timepoint of study. To truly determine the mechanism of a drug, it is necessary to investigate its effects in healthy control mice. We therefore compared NR-treatment to vehicle-treatment in control mice, and we juxtapose this with the comparison of NR-treatment to vehicle-treatment in Alport mice.

In both sexes, metabolism-related GO biological processes were among the most highly enriched set that were simultaneously upregulated in both NR-treated control mice (versus vehicle-treated control mice) (**Figs. 5A-B** ) and in NR-treated Alport mice (versus vehicle-treated Alport mice) (**Fig. S15** ). KEGG pathway analyses mirrored these results (**Figs. 5C-D** and **S16**), and transcription factor analyses also predicted the activation of RXR/PPARα in the NR-treated samples (**Figs. 5E-F**). The striking similarity of NR treatment in both control and Alport mice is demonstrated by visualization on KEGG graphs for the PPAR signaling pathway (KEGG Entry No. 03320) (**Figs. S17-18**), the peroxisome (KEGG Entry No. 04146) (**Figs. S19-20**), and fatty acid degradation (KEGG Entry No. 00071) (**Figs. S21-22**).

**Figure 5:**
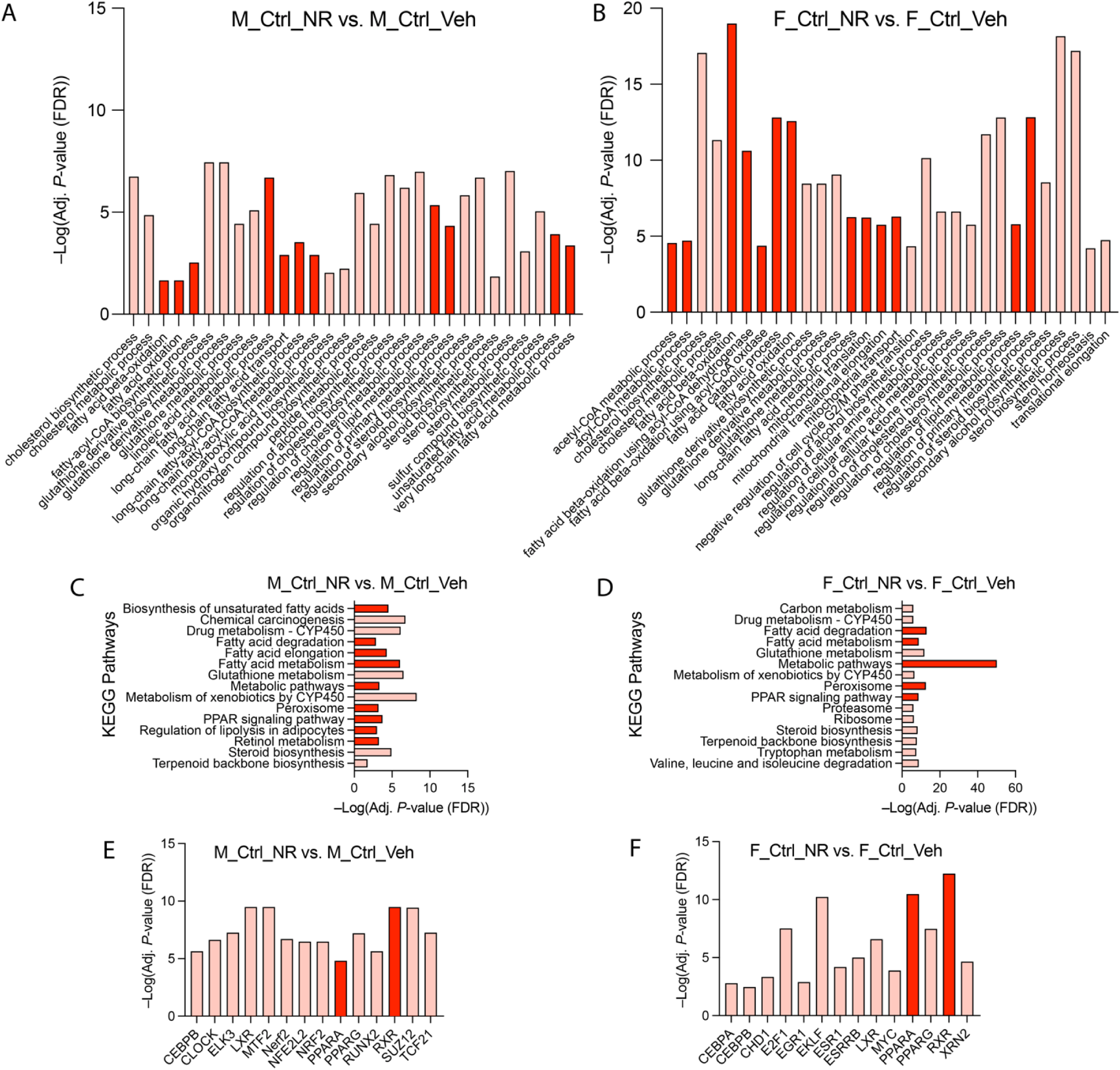
NAD^+^ supplementation activates renal metabolism in control mice, not just in Alport mice. Bulk kidney cortex RNA-seq data from control and Alport mice, treated with or without NR, were analyzed (*N* = 4 mice per group). Gene ontology (GO) biological processes and KEGG pathways that were simultaneously upregulated in both NR-treated control mice (vs. vehicle-treated control mice) and in NR-treated Alport mice (vs. vehicle-treated Alport mice) were identified. (**A,B**) GO biological processes that were increased in NR-treated control mice (vs. vehicle-treated control mice) in males (*A*) and females (*B*). (**C,D**) KEGG pathways that were increased in NR-treated control mice (vs. vehicle-treated control mice) in males (*C*) and females (*D*). These data (*A-D*) are presented adjacent to the corresponding sex-match comparisons from NR-treated Alport mice (vs. vehicle-treated Alport mice) in the *Supplemental Material* . (**E,F**) Transcription factor analyses suggests that the RXR/PPARα gene regulatory network is activated in NR-treated control mice (vs. vehicle-treated control mice) in both males (*E*) and females (*F*).

In summary, the effect of NR treatment on metabolism-related pathways in control mice was essentially identical to that in Alport mice. These results strongly suggest that NR protects the kidney *via* activating renal metabolism, more specifically, the RXR/PPARα signaling pathway that stimulates FAO. We then sought to confirm these results with orthogonal biochemical assays, but prior to doing so, we investigated the effects of genetic background and sex.

### The renal transcriptome of Alport mice is similar across genetic backgrounds

Experimental investigations on Alport syndrome typically use *Col4a3*-null mice on the 129 or B6 genetic backgrounds, and occasionally F1 hybrids thereof. Alport mice on the 129 background develop kidney disease much more rapidly than their counterparts on the B6 background, and 129.B6F1 hybrids have an intermediate phenotype (54). We performed a meta-analysis to confirm that the changes in renal transcriptome we observed in our B6 Alport colony are representative of *Col4a3*-null mice on the other backgrounds.

Previously published RNA-seq data from male control and Alport mice on the 129 and 129.B6F1 backgrounds were compared with vehicle-treated male control and Alport mice from the current study (B6 background). Principal component analysis demonstrated separation by genotype (PC1, 73.2% of variance). The second-largest principal component accounted for only 3.2% of variance, and it was not associated with genetic background (**Fig. S23A** ). KEGG pathway analyses showed similar impairments in metabolic processes across all genetic backgrounds and the combined meta-analysis (**Figs. S23B-E**).

These data show that the changes in renal transcriptome in Alport mice are independent of genetic background when compared to their respective controls. These results also suggest, but do not prove, that NR treatment would have a similar mechanism of action in all Alport mice regardless of genetic background.

### Male and female Alport mice have distinct inflammatory and fibrotic responses

Alport syndrome affects both males and females, but it has not yet been reported if the disease progression differs between the sexes at the molecular level. Of the six RNA-seq datasets from Alport mouse kidneys that are deposited in the Gene Expression Omnibus, only one is from both sexes (30, 31, 55–59). However, it used a unique outbred model that is the first of its kind (56). To address this gap, we analyzed the subset of 16 vehicle-treated mouse kidneys, four from each sex/genotype combination.

Principal component analysis showed that the data was tightly clustered with respect to both genotype (PC1, 57.1% of variance, *P* = 4.66×10^-08^) and sex (PC2, 14.3% of variance, *P* = 9.27×10^-08^) (**Fig. S24A**). A total of 114 genes were differentially regulated with both sex and disease, as identified by a sex:genotype interaction (**Fig. S24B**). GO enrichment analysis revealed that extracellular matrix and neovascularization processes were comparatively less upregulated in female Alport mice than in male Alport mice when compared to their sex-matched controls (**Figs. S24C**). In many cases, genes that were increased in female Alport mice (versus female control mice), such as the pro-fibrotic and pro-inflammatory genes *C3*, *Col1a2*, *Fgfbp1*, and *Ticam2* (**Fig. S24D**), were increased to a greater degree in male Alport mice (versus male control mice). We refer to this gene set as having a negative sex:genotype interaction because the increase in Alport mice compared to control mice is smaller (or even decreased, not increased) in female mice compared to male mice. An additional 70 genes had a negative sex:genotype interaction (**Fig. S25A** ), while 40 genes had a positive sex:genotype interaction (**Fig. S25B**). To further parse out a potential role of sex as a biological variable, pathway analysis using a modified GSEA algorithm was performed on the entire dataset as previously described (39, 60), not just the subset of genes with a significant sex:genotype interaction. This also identified inflammation-related pathways as differentially regulated with both sex and disease (**Fig. S26** ). Despite these transcriptional differences, there were no differences between the sexes in histological markers of fibrosis or inflammation (**Fig. S3**).

In summary, these data support moderate differential regulation of fibrosis- and inflammation-related pathways between the sexes at the transcriptional level in this mouse model of Alport syndrome. Importantly, neither analysis identified a sex:genotype interaction in metabolism-related pathways, our primary area of interest in this study. Therefore, we chose to only study male mice for all remaining biochemical assays because they seemed to have a more severe phenotype at the transcriptional level. However, the sexes were pooled for the snRNA-seq experiment due to the outsized benefit these data could have for the scientific community.

### NAD^+^ supplementation restores kidney mitochondrial fatty acid oxidation

Once we confirmed the absence of a sex:genotype interaction in metabolism-related pathways, we then returned to investigating the effects of NR treatment. Mitochondrial dysfunction is a well-accepted mechanism seen in models of CKD, including in a mouse model of Alport syndrome (30). In addition, enhancing *de novo* NAD^+^ synthesis has been shown to improve mitochondrial FAO (61). We therefore hypothesized that mitochondrial FAO would be impaired in Alport mice and restored by NR treatment. To test this, we immunoblotted for key proteins in the mitochondrial FAO pathway: PGC-1α, carnitine palmitoyltransferase 1-α (CPT1α), and medium-chain acyl-coenzyme A dehydrogenase (MCAD).

PGC-1α is a coactivator that regulates PPARα function, and both CPT1α and MCAD are PPARα target genes (6, 62, 63). Consistent with the observed effects of NR in both control and Alport mice on RNA-seq, PCG- 1α levels were increased with NR treatment in both genotypes (**Fig. 6A-B**), and this serves as a good positive control for the effects of NR.

**Figure 6:**
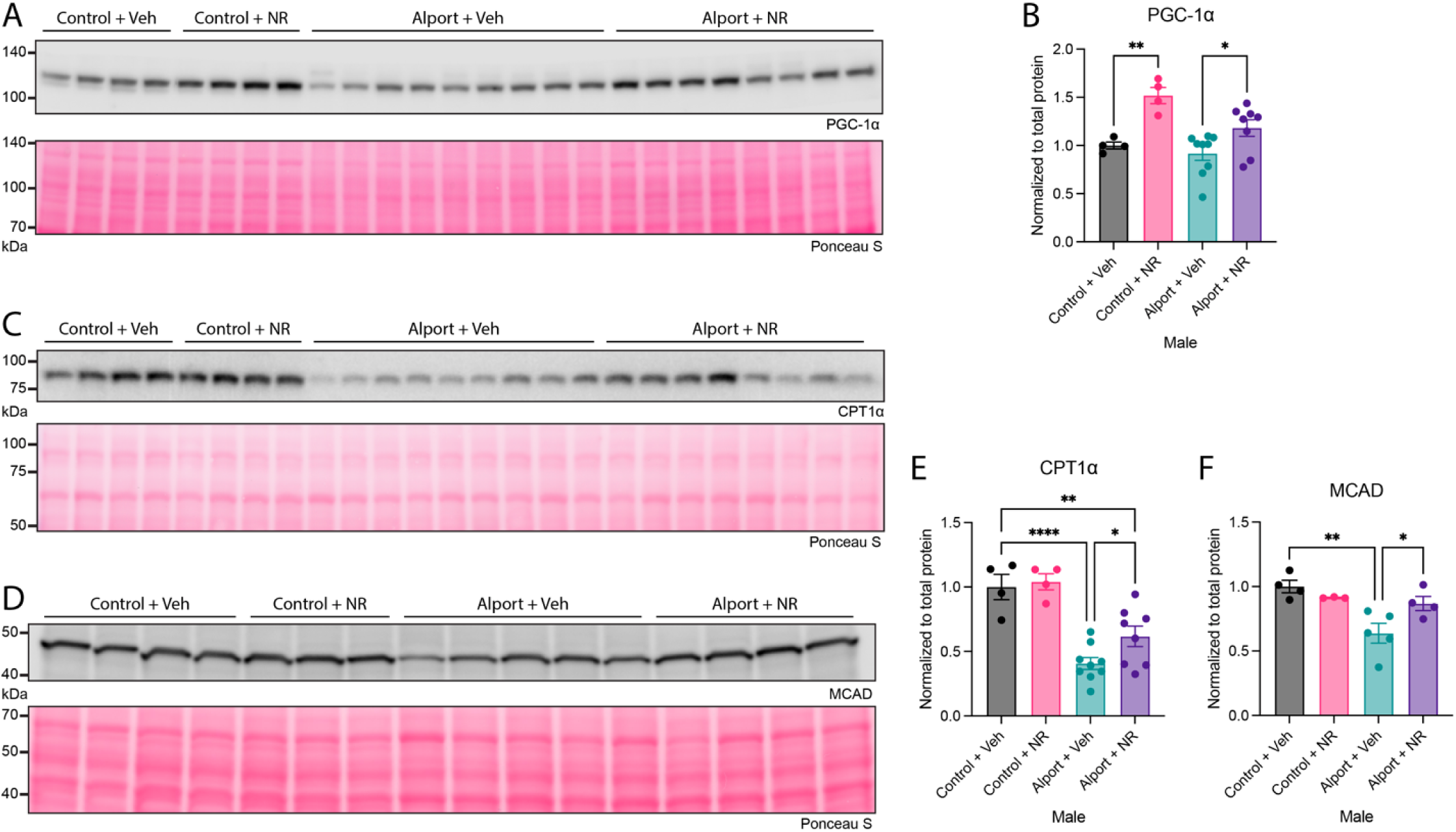
NAD^+^ supplementation activates renal fatty acid metabolism. (**A,B**) NR treatment increased kidney PGC-1α in both control and Alport mice. (**C-F**) Kidney CPT1α (*C,E*) and MCAD (*D,F*), key players in mitochondrial fatty acid oxidation, were reduced in Alport mice and restored by NR treatment. Ponceau S, a nonspecific protein stain, was used as a loading control. Significance was determined by one-way ANOVA with the Holm– Šídák correction for multiple comparisons. Data are expressed as the means ± SEM. Each datum represents one mouse. \**P* < 0.05, \*\**P* < 0.01, \*\*\*\**P* < 0.0001. CPT1α, carnitine palmitoyltransferase 1-α; MCAD, medium-chain acyl-coenzyme A dehydrogenase; NR, nicotinamide riboside; PGC-1α, peroxisome proliferator-activated receptor-γ coactivator 1-α; Veh, vehicle.

CPT1α and MCAD are both integral to mitochondrial FAO, and defects in either protein can cause severe clinical diseases (64). CPT1α controls fatty acyl-CoA transport into the mitochondria, and it is the rate-limiting step of long-chain FAO and medium-chain FAO of nine or more carbons in length (65). In our samples, CPT1α was reduced in Alport mice, and it was restored with NR treatment (**Figs. 6C** and **6E**). Medium-chain fatty acids, at least those up to eight carbons in length, can enter the mitochondria independent of CPT1α and then are converted to fatty acyl-CoAs once inside (65). However, regardless of how they enter the mitochondria, MCAD catalyzes the first step in mitochondrial FAO of all lengths of medium chain acyl-CoAs (64). In our samples, MCAD was also reduced in Alport mice (versus control mice) and restored by NR treatment (**Figs. 6D** and **6F**).

These changes in the key enzymes regulating both medium- and long-chain FAO are robust orthogonal data that validate the transcriptomic changes seen on RNA-seq. They further demonstrate that Alport mice have impaired fatty acid utilization and that NR protects the kidney by activating renal metabolism.

### NAD supplementation restores the canonical functions of the proximal tubule, and reduces immune cell and myofibroblast appearance

The data presented thus far strongly establish that NR protects the kidney via activating renal cortical transcription of metabolic genes, and this is associated with corresponding improvements in renal expression of mitochondrial FAO proteins. However, they provide little insight into the molecular changes occurring within individual cells. To investigate the cell type specific effects of NR treatment in Alport syndrome, as well as to show unambiguously that NR stimulates metabolism in the proximal tubules, we performed snRNA-seq on our samples.

Nuclei from control and Alport mice, both with and without NR treatment, were extracted, sequenced with snRNA-seq, and merged to create an atlas of thirty-one cell type clusters (**Fig. 7A**). Clusters were defined by known marker gene expression and most cell types were represented across all four conditions (**Fig. S27**). However, the proportion of cell type distribution varied across conditions (**Fig. 7B**). Podocytes and reference proximal tubule (PT) cells were more represented in control mice, whereas adaptive PT (aPT) and injured PT (iPT) had greater representation in Alport mice. Vehicle-treated Alport mice had enrichment of characteristic injury pathways in both the PT (**Fig. S28** ) and podocytes (**Fig. S29** ) as compared to vehicle-treated control mice. These pathways included fatty acid degradation, PPAR signaling, cell adhesion, and immune signaling. Osteopontin (*Spp1*) and Neutrophil gelatinase-associated lipocalin (*Lcn2*) were upregulated in both the PT and podocyte.

**Figure 7:**
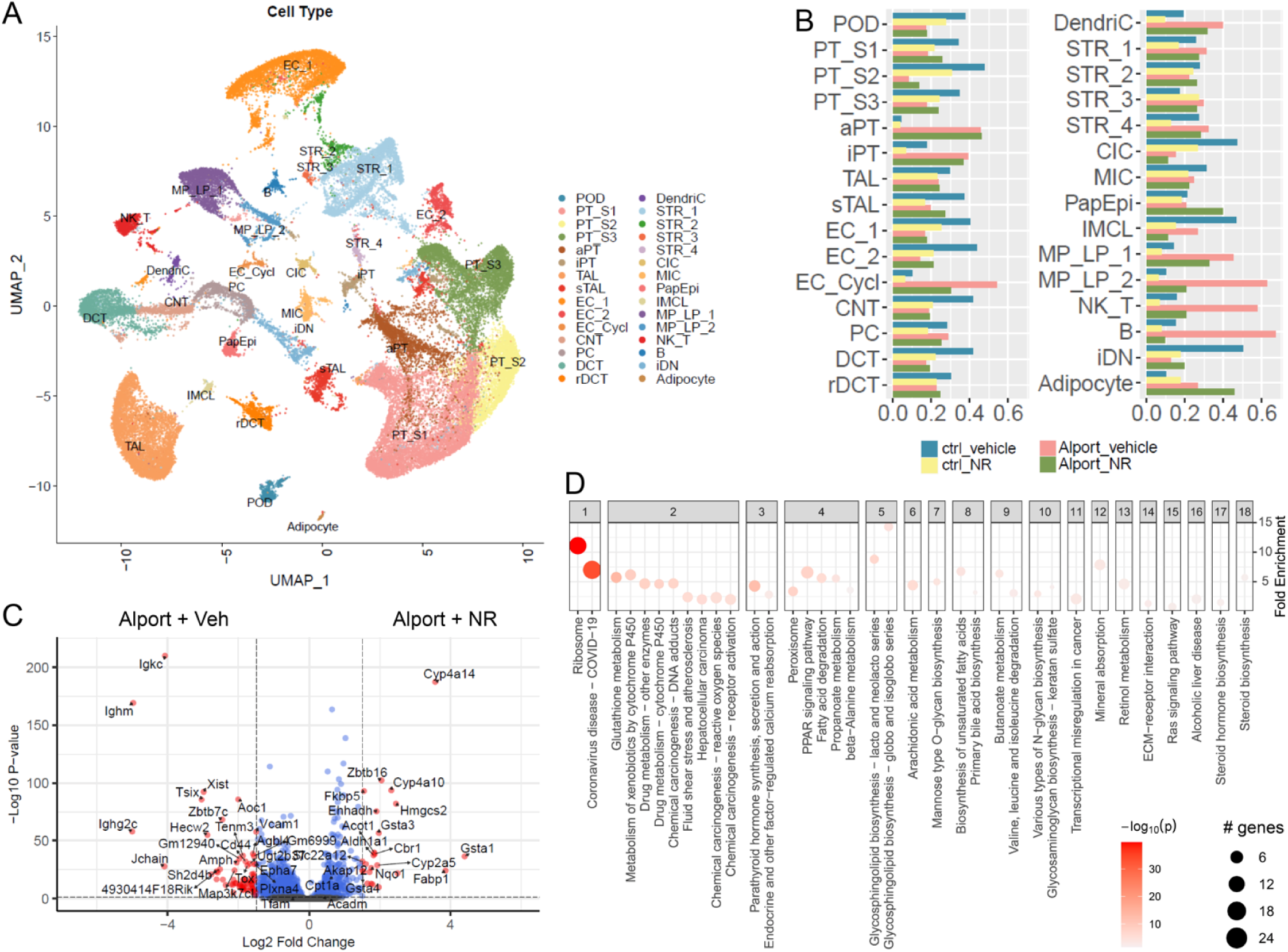
Single nucleus RNA-sequencing of control and Alport mice, with and without NAD^+^ supplementation. (**A**) The snRNA-seq reduction by uniform manifold approximation and projection (UMAP) of 49,488 nuclei in 31 clusters. (**B**) Proportion of nuclei per cluster by condition. Myofibroblasts and immune cells were upregulated in Alport mice. The effect was mitigated by NR treatment. (**C**) Differentially expressed genes in NR-treated Alport mice (vs. vehicle-treated Alport mice). Genes upregulated by NR-treatment are displayed on the right. Mice treated with NR had reduced expression of immune signaling transcripts such as *Ncam2* and *Jchain*. (**D**) Enriched pathways between Alport mice treated with vehicle and NR. NR treatment significantly impacted translation (ribosome), metabolism, endocrine function, and PPAR signaling in the proximal tubule cell. Ctrl, control; NR, nicotinamide riboside; Veh, vehicle.

The injury phenotypes of the proximal tubule cell and podocyte in Alport mice shared properties. In the NR-treated proximal tubule, we found transcriptomic evidence supporting restoration of several vital cellular functions: translation (ribosome), metabolism, endocrine function, fatty acid degradation, and PPAR signaling (**Figs. 7C-D** ). In the podocyte, the enriched pathways of NR treatment also included PPAR signaling and fatty acid degradation, although fewer genes were differentially expressed due to the smaller sample size (**Fig. S30**).

We next sought to understand the cell-type specific and spatially anchored gene expression changes of *Cpt1a* and *Acadm* (MCAD) (**Fig. 8**). *Cpt1a* and *Acadm* expression by snRNA-seq was significantly increased in the proximal tubule S1 and S2 cells of Alport mice after NR treatment (**Fig. 8A** ). Spatial transcriptomic (ST) profiling supported the observed effects seen in the snRNA-seq atlas. ST revealed that expression of *Cpt1a*, the rate limiting step for mitochondrial long-chain FAO, is reduced in Alport mice and restored with NR treatment (**Figs. 8B** and **S31A**). We assessed the spatial distribution of *Cpt1a* expression across all spots (pseudobulk), in functional tissue units of glomeruli, the cortical tubulointerstitium (sans glomeruli), and the medullary outer stripe. These functional tissue units were selected by a combination of histologic assessment and marker gene expression with *Wt1* for glomeruli, *Slc34a1* for cortical tubulointerstitium, and *Slc3a1* for outer stripe (**Fig. S32** ). *Cpt1a* expression was significantly upregulated in the pseudobulk, cortex, and outer stripe in Alport mice after NR treatment (**Fig. 8C**). Similarly, *Acadm* (the gene for MCAD) expression is reduced in the Alport condition, and this reduction is mitigated with NR treatment (**Figs. 8D-E** and **S31B**). The expression of *Cpt1a* and *Acadm* in glomeruli of Alport mice treated with NR trended toward an increase, although the number of glomerular spots was small. *Tfam* (mitochondrial transcription factor A) expression was neither differentially expressed in the PT S1 or S2 cells of the snRNA-seq dataset (**Figs. 8A** and **S31C**). We assessed differential gene expression in the cortex of the ST samples (**Fig. 8F** ) and found similar pathways were enriched to those observed in the snRNA-seq dataset (**Fig. 8G**).

**Figure 8:**
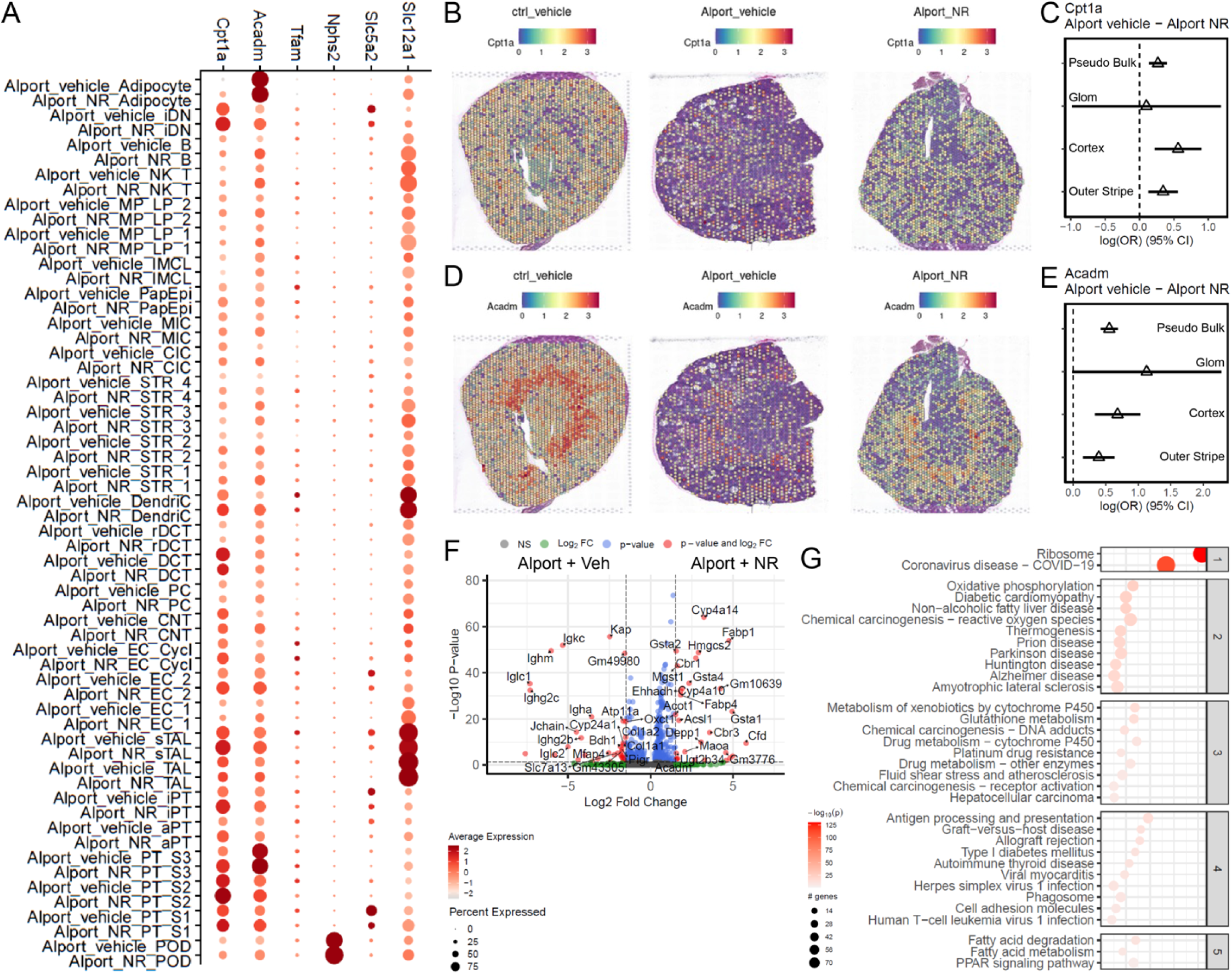
Spatial localization of *Cpt1a* and *Acadm*. (**A**) Expression of *Acadm*, *Cpt1a*, and *Tfam* in cell types and conditions by single nucleus RNA sequencing. (**B**) Visium spatial transcriptomics was performed on vehicle-treated male control mice (*left*), vehicle-treated male Alport mice (*center*), and NR-treated male Alport mice (*right*). *Cpt1a* expression is depicted. (**C**) *Cpt1a* expression was restored with NR treatment. (**D,E**) Expression of Acadm was reduced in Alport mice and restored after NR treatment. (**F**) Differentially expressed genes within the cortical tubulointerstitium in NR-treated Alport mice (vs. vehicle-treated Alport mice). Genes upregulated by NR-treatment are displayed on the right. (**G**) Pathway enrichment in the cortical tubulointerstitium in NR-treated Alport mice (vs. vehicle-treated Alport mice). CI, confidence interval; Cortex, cortical tubulointerstitium; Ctrl, control; Glom, glomerulus; NR, nicotinamide riboside; OR, odds ratio with 95% CI; Veh, vehicle.

The proportions of myofibroblasts, macrophages, T lymphocytes, and B lymphocytes were all increased in Alport mice within the snRNA-seq atlas. NR treatment was successful in reducing immune cell infiltration and fibrosis associated with the Alport condition. Using spatial transcriptomics, these cell types were localized to the tissue with a label transfer method (**Fig. S33**). NR treatment led to reduced immune cell infiltration and stromal cell appearance in Alport mice. Taken together, these data suggest that NR treatment restores the canonical functions of the PT, restores expression of *Cpt1a* and *Acadm*, reduces immune cell infiltration, and reduces myofibroblast appearance and fibrosis.

## DISCUSSION

Herein, we have comprehensively shown that NR protects the kidney in a mouse model of Alport syndrome. In addition, we report several other noteworthy contributions. We are the first to report single cell (nuclei) and spatial transcriptomic data from the Alport model. We are also the first to report sex-differences in the bulk renal transcriptome of Alport mice. Finally, at both the bulk and single cell (nuclei) level, we report that NR-mediated renal activation of metabolism occurs in healthy control mice and diseased Alport mice. The similarity in response between both control and Alport mice strongly suggests that normalization of renal metabolism is responsible for the nephroprotective effect of NAD^+^ supplementation in CKD, a key mechanistic insight that was previously unknown.

In our experiments, we identified that activation of renal metabolism in the proximal tubule via the NAD^+^–PGC-1α– PPARα–FAO axis is an important mechanism contributing to the nephroprotective effects of NAD^+^ supplementation in CKD. Our study builds upon prior research on the protective PPARα–FAO axis in Alport mice by identifying the importance of the NAD^+^–PGC-1α axis upstream of it (30). However, perhaps more importantly, we showed that activation of renal metabolism is, at least in part, the direct result of NAD^+^ supplementation—it is not just secondary to the prevention of kidney disease. The highly similar transcriptional responses of control and Alport mice to NAD^+^ supplementation allows us to make this key mechanistic insight, and it underscores the importance of also investigating the effects of drugs in the absence of disease. Nevertheless, although our results show that NAD^+^ supplementation activates renal metabolism and protects against CKD, they do not definitively demonstrate the role of NAD^+^ depletion in the pathogenesis of CKD. For that, further study using genetic knockout models would be needed.

Sex as a biological variable has been, and continues to be, a neglected area of research. It has been reported that male and female *Col4a3*^tm1Dec^ mice on the 129/SvJ background have similar severities of kidney disease (66). We note that this is unusual because male mice are generally more susceptible to kidney disease than female mice (67). Even though the current study was not designed as a comprehensive comparison between the sexes, we still identified moderate sex-specific differential regulation of fibrosis- and inflammation-related pathways at the transcriptional level, albeit not at the histological level. Our gene set represents a starting point that may assist in developing hypotheses to investigate a potential phenotypic sex-difference in this mouse model of Alport syndrome.

Modern transcriptomic approaches, such as snRNA-seq and spatial transcriptomics, provide unparalleled insights into the molecular mechanisms of disease, and there is a focused effort to develop atlases with single-cell resolution (68). Because snRNA-seq is still relatively new, datasets are not yet available for many disease models, especially rare diseases such as Alport syndrome. Our snRNA-seq and spatial transcriptomics datasets are the first of their kind from a model of Alport syndrome. In addition, we also report the first characterization of how NAD^+^ supplementation affects the kidney at the single cell level in both states of health and disease. Our snRNA-seq dataset is further differentiated by the inclusion of both sexes and the simultaneous assay for transposase-accessible chromatin, known as snATAC-seq. However, a full comparison between the sexes, as well as integration of the multiome data, is beyond the scope of the current publication.

Although it has been known that exogenous NAD^+^ supplementation protects from CKD (11–16), the cell types which contribute to this benefit were not rigorously investigated. Models of AKI displayed the strongest and most consistent therapeutic benefit of NAD^+^ supplementation, implying the proximal tubules as a likely location. However, the data from models of CKD were less clear, and several studies suggested a potential role for podocytes (13, 69). Our data demonstrates the prominent role of the proximal tubule in NAD^+^-mediated protection from CKD. However, one limitation of our data was the relatively lower sample size of podocytes compared to proximal tubular epithelial cells in both the snRNA-seq and ST datasets—a challenge common to many single cell studies. While the effects of NAD^+^ supplementation were clear in the proximal tubule, whether NAD^+^ supplementation holds a direct effect on podocytes in Alport disease remains an outstanding question. The snRNA-seq and ST data both suggest a trend toward restored *Cpt1a* and *Acadm* expression in most cell types and functional tissue units.

In summary, NAD^+^ supplementation protects the kidney in a mouse model of Alport syndrome. Mechanistically, NR activates renal metabolism by restoring the NAD^+^–PGC-1α–PPARα–FAO axis within the proximal tubules. Future directions include determining if NAD^+^ supplementation exerts similar effects on other cell types within the kidney and using genetic knockout models to definitively demonstrate the role of tubular NAD^+^ depletion in this model of CKD.

## Acknowledgments

We acknowledge all Levi Lab members, especially Isabel Schaffer and Sharmila Adapa for assistance with polarized microscopy. We thank Bo Zhang and Amanda Knoten for assistance with the snRNA-seq experiments and the Washington University Kidney Translational Research Center for support. We thank members of the Computational Microscopy Imaging Laboratory at the University of Florida for maintaining a cloud-based instance of HistoCloud. Research reported in this publication was supported by the National Institutes of Health Grants F30DK129003 (to B.A.J.), TL1TR001431 (to B.A.J.), R01DK116567 (to M.L.), R01DK127830 (to M.L.), the Georgetown University Lombardi Comprehensive Cancer Center Support Grant P30CA051008, the Washington University Pediatric Center of Excellence in Nephrology grant P50DK133943 (S.J.), and the Indiana University O’Brien Center for Advanced Microscopic Analysis grant U01DK114923 (M.T.E.).

## Author contributions

Conceptualization: BAJ and ML. Methodology: BAJ and ML. Software: BAJ, DLG, RMF, BAS, and MTE. Validation: BAJ, DLG, BAS, and ML. Formal analysis: BAJ, DLG, ET, JP, RMF, and MTE. Investigation: BAJ, AS, YC, KK, and XY. Resources: BAJ, SJ, MTE, and ML. Data Curation: BAJ, DLG, RMF, and MTE. Writing - Original Draft: BAJ. Writing - Review & Editing: BAJ, DLG, KM, AS, YC, ET, JP, KK, RMF, XY, BAS, KCA, TY, XXW, AZR, SJ, MTE, and ML. Visualization: BAJ and DLG. Supervision: ML. Project administration: ML. Funding acquisition: BAJ, SJ, MTE, and ML.

