## Supplemental Material for "NAD^+^ activates renal metabolism and protects from chronic kidney disease in a model of Alport syndrome"

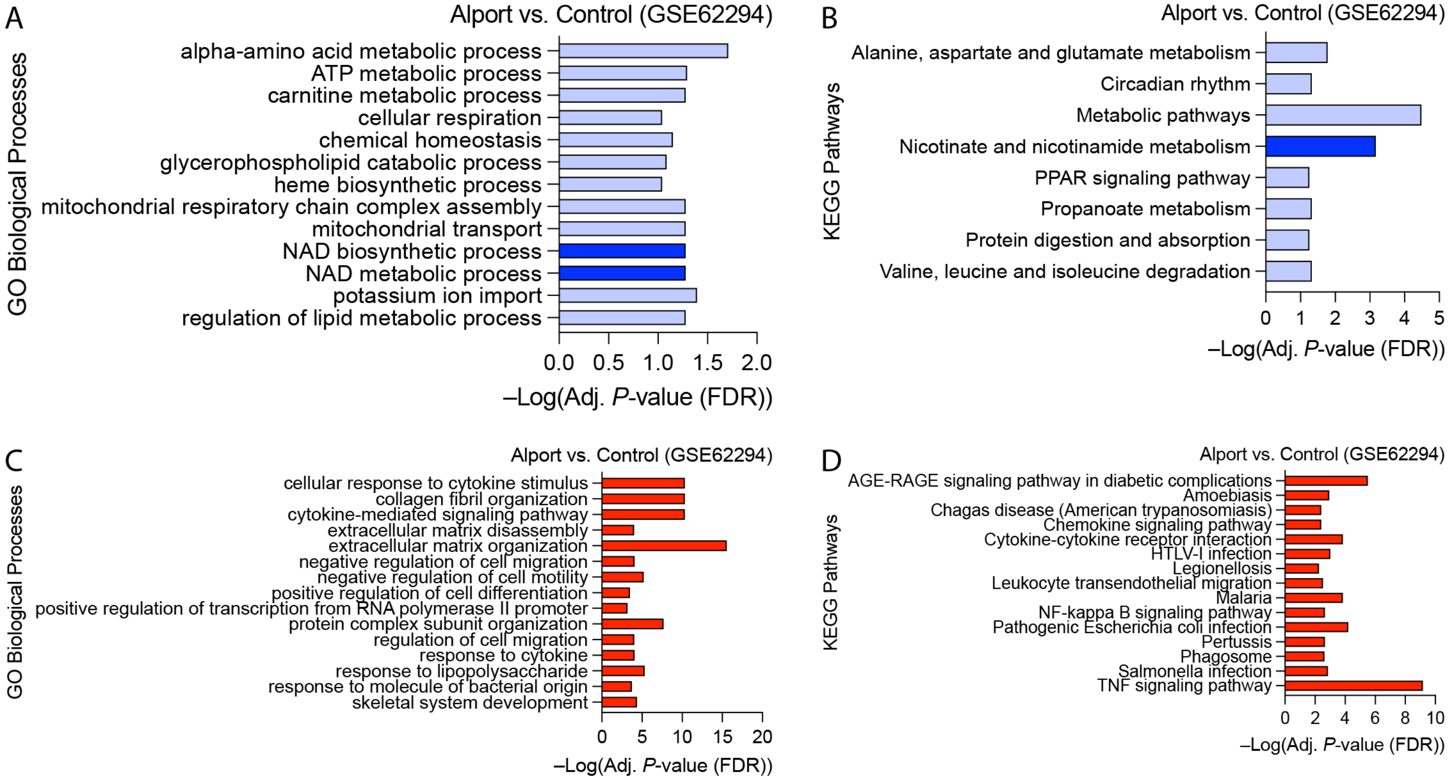

**Figure S1: NAD+ biosynthetic pathways are reduced in Alport mice.** RNA-seq data from a previously published study was downloaded from the Sequence Read Archive and re-analyzed (1-3). Kidneys from Alport mice ($N$ = 4) were compared to control mice ($N$ = 3) on the 129X1/SvJ background, 5.5 weeks in age. (**A-B**) GO enrichment (*A*) and KEGG pathway (*B*) analyses show, respectively, that biological process and pathways related to NAD+ biosynthesis and metabolism are reduced in Alport mice compared to control mice. These pathways are emphasized by dark blue shading. (**C-D**) In contrast to this, GO enrichment (*C*) and KEGG pathway (*D*) analyses show, respectively, that biological process and pathways related to fibrosis and inflammation are increased in Alport mice compared to control mice. Red indicates an increase in Alport mice vs. control mice, and blue indicates a decrease.

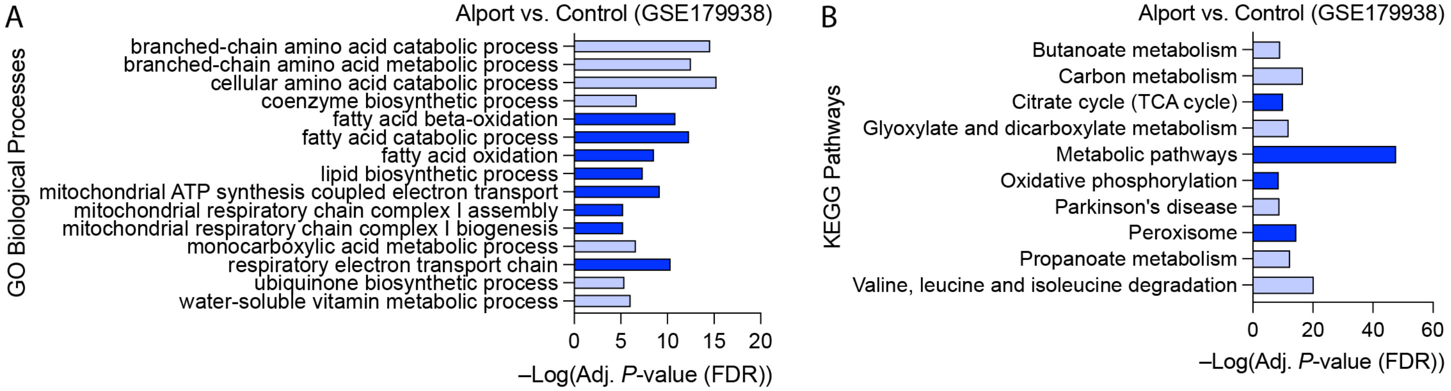

**Figure S2: Alport mice have impaired renal metabolism.** RNA-seq data from a previously published study was downloaded from the Sequence Read Archive and re-analyzed (2-4). Kidneys from Alport mice ($N$ = 6) were compared to control mice ($N$ = 5) on a mixed 129/SvJ and C57BL/6J background, 15 weeks of age. (**A-B**) GO enrichment (*A*) and KEGG pathway (*B*) analyses show, respectively, that biological process and pathways related to renal metabolism, including fatty acid oxidation, are reduced in Alport mice compared to control mice. These pathways are emphasized by dark blue shading. Blue indicates a decrease in Alport mice vs. control mice.

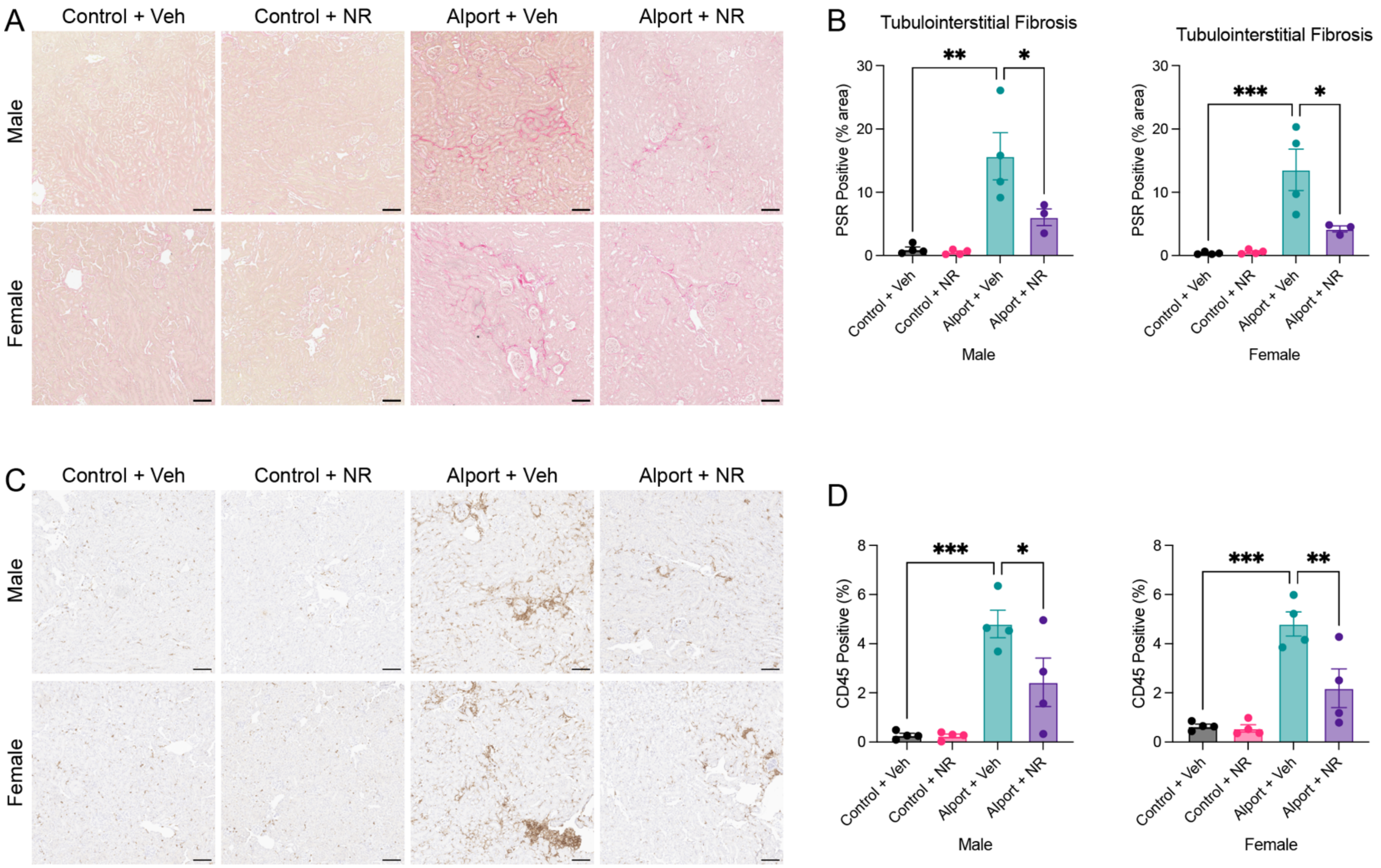

**Figure S3: NAD$^+$ supplementation prevents tubulointerstitial fibrosis and inflammation.** Fibrosis and inflammation were quantified in the subset of samples sent for bulk RNA-sequencing. (**A**) Representative images of PSR-stained kidneys acquired with unpolarized light. Deep red is specific for fibrosis. (**B**) Quantification of PSR-stained kidneys shows that NR treatment reduced renal cortical tubulointerstitial fibrosis in male and female mice. (**C**) Representative images of CD45-stained kidneys. Brown cell membrane staining is specific for inflammatory cells. (**D**) Quantification of CD45-stained kidneys shows that NR treatment reduced renal tubulointerstitial inflammation in both sexes. Scale bars represent 100 µm. Significance was determined by one-way ANOVA with the Holm–Šídák correction for multiple comparisons. Data are expressed as the means ± SEM, and each datum represents one mouse. *P < 0.05, **P < 0.01, ***P < 0.001. NR, nicotinamide riboside; PSR, picrosirius red; Veh, vehicle.

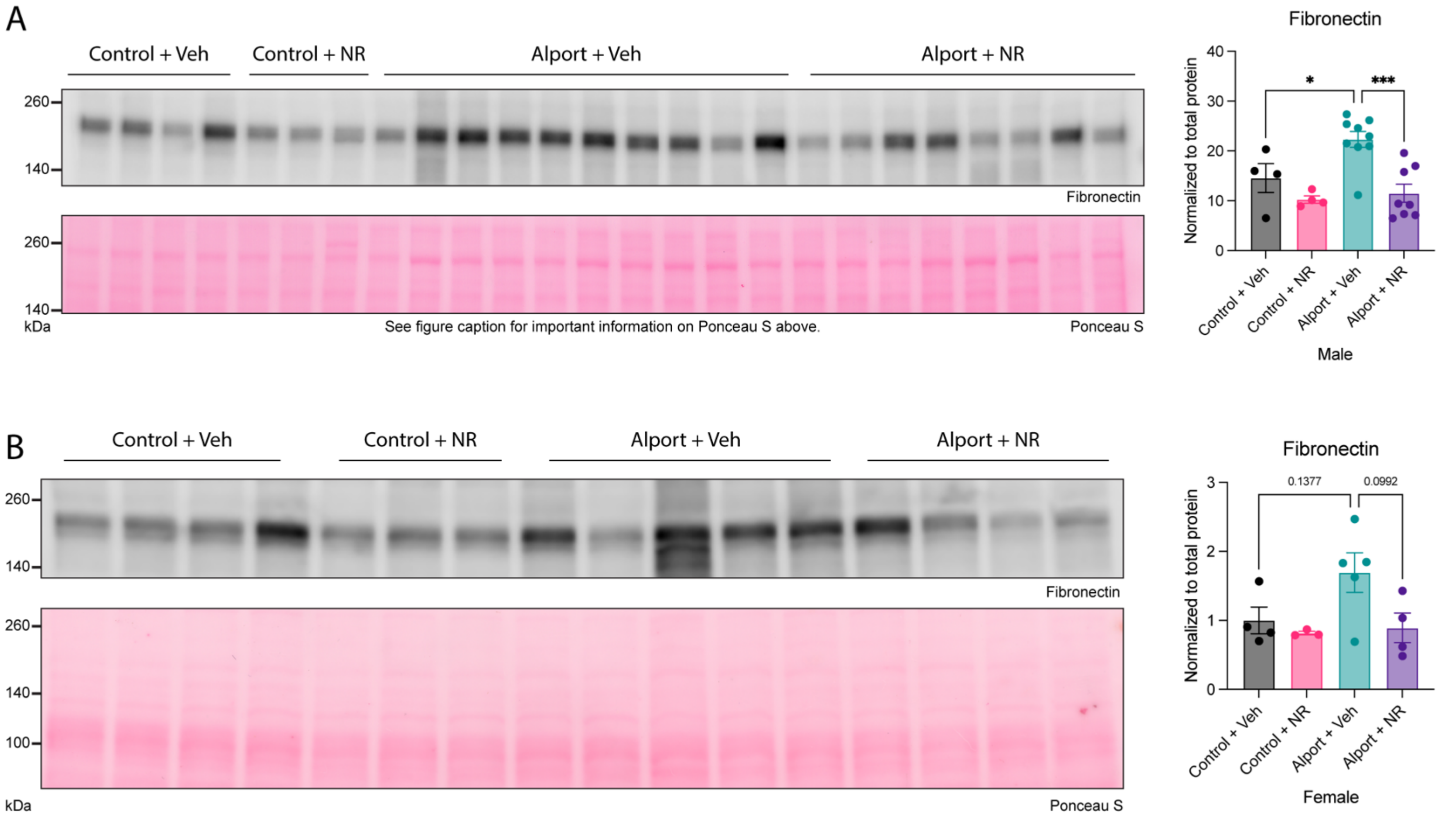

**Figure S4: NAD$^+$ supplementation lowers kidney expression of fibronectin.** (**A**) On immunoblot, fibronectin is increased in vehicle-treated male Alport mice compared to vehicle-treated male control mice. NR treatment in male Alport mice reduces fibronectin expression compared to vehicle-treated male Alport mice. This immunoblot was horizontally cut prior to incubation with the primary antibodies for fibronectin and PGC-1α, presented in the main text. Therefore, data from those blots were normalized to the same Ponceau S image, of which the upper portion is presented here. (**B**) On immunoblot, fibronectin is increased (trend, $P < 0.15$) in vehicle-treated female Alport mice compared to vehicle-treated female control mice. NR treatment in female Alport mice reduces fibronectin expression (trend, $P < 0.10$) compared to vehicle-treated female Alport mice. Ponceau S, a nonspecific protein stain, was used as a loading control. Significance was determined by one-way ANOVA with the Holm–Šídák correction for multiple comparisons. Data are expressed as the means ± SEM. Each datum represents one mouse. *$P < 0.05$, ***$P < 0.001$. NR, nicotinamide riboside; PGC-1α, peroxisome proliferator-activated receptor gamma coactivator 1 alpha; Veh, vehicle.

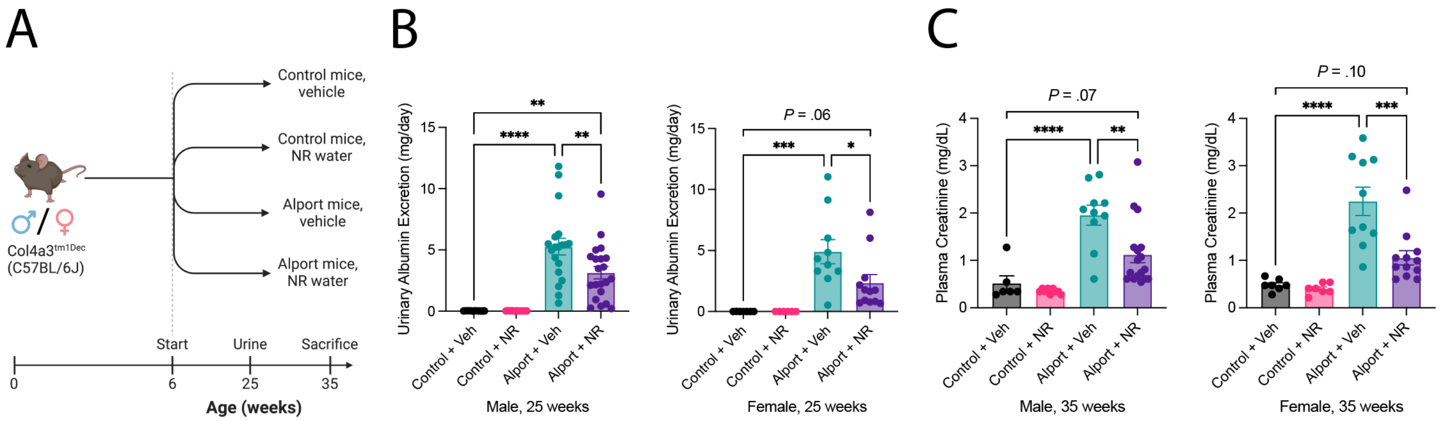

**Figure S5: Replication study showing that NAD+ supplementation prevents kidney disease in Alport mice.** (**A**) Experimental design: Control and Alport mice of both sexes were treated with or without NR between 6-weeks and 35-weeks of age. Urine was collected at 25 weeks of age. (**B**) NR treatment reduced 24-hour urinary albumin excretion in Alport mice of both sexes at 25 weeks of age. (**C**) NR treatment reduced plasma creatinine in Alport mice of both sexes at 35 weeks of age. Significance was determined by one-way ANOVA with the Holm–Šídák correction for multiple comparisons. Data are expressed as the means ± SEM. Each datum represents one mouse. **P* < 0.05, ***P* < 0.01, ****P* < 0.001, *****P* < 0.0001. NR, nicotinamide riboside; Veh, vehicle.

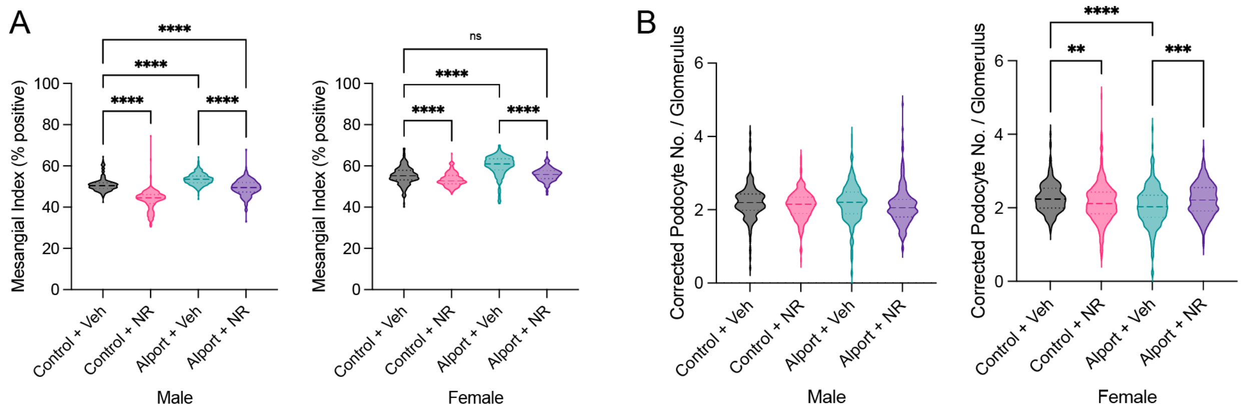

**Figure S6: NAD+ supplementation prevents glomerular injury.** (**A**) Alport mice had an increased mesangial index that was reduced with NR treatment. (**B**) The corrected podocyte number per glomerulus was decreased in female Alport mice and restored by NR treatment. There were no differences in the corrected podocyte number per glomerulus in male mice. However, these results should be interpreted with caution because, unlike the volumetric podocyte density, the corrected podocyte number per glomerulus does not control for differences in glomerular volume between groups. Significance was determined by one-way ANOVA with the Holm–Šídák correction for multiple comparisons. Each datum represents one glomerulus. ***P* < 0.01, ****P* < 0.001, *****P* < 0.0001. No., number; NR, nicotinamide riboside; Veh, vehicle.

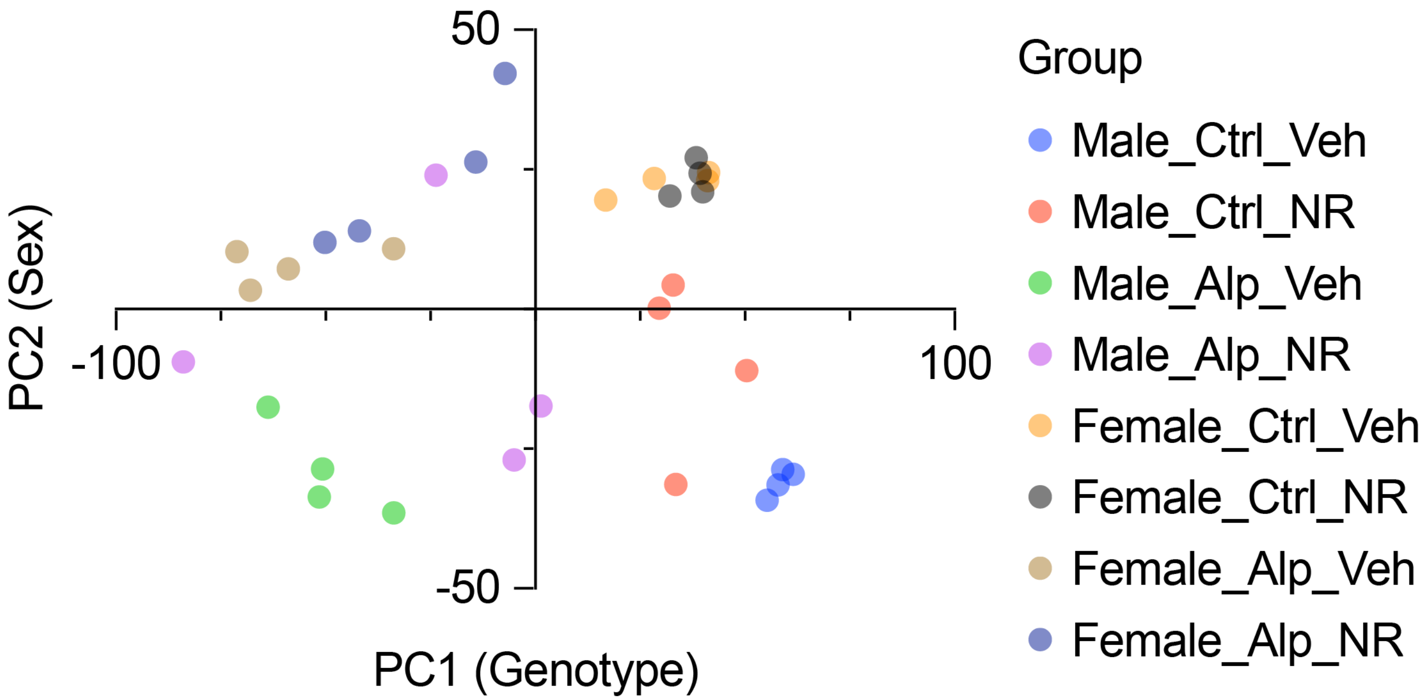

**Figure S7: Principal component analysis shows that NAD$^+$ supplementation protects the kidney.** Principal component analysis was performed on bulk kidney cortex RNA-seq data from control and Alport mice treated with or without nicotinamide riboside. PC1 is correlated with the genotype, and it is responsible for 42.7% of variance ($P = 2.75\times10^{-11}$). PC2 is correlated with the sex, and it is responsible for 11.3% of variance ($P = 1.16\times10^{-07}$). PC3 is correlated with treatment (data not shown), and it is responsible for 7.3% of the variance ($P = 2.07\times10^{-3}$). Each datum represents one mouse. Alp, Alport; Ctrl, control; NR, nicotinamide riboside; PC, principal component; Veh, vehicle.

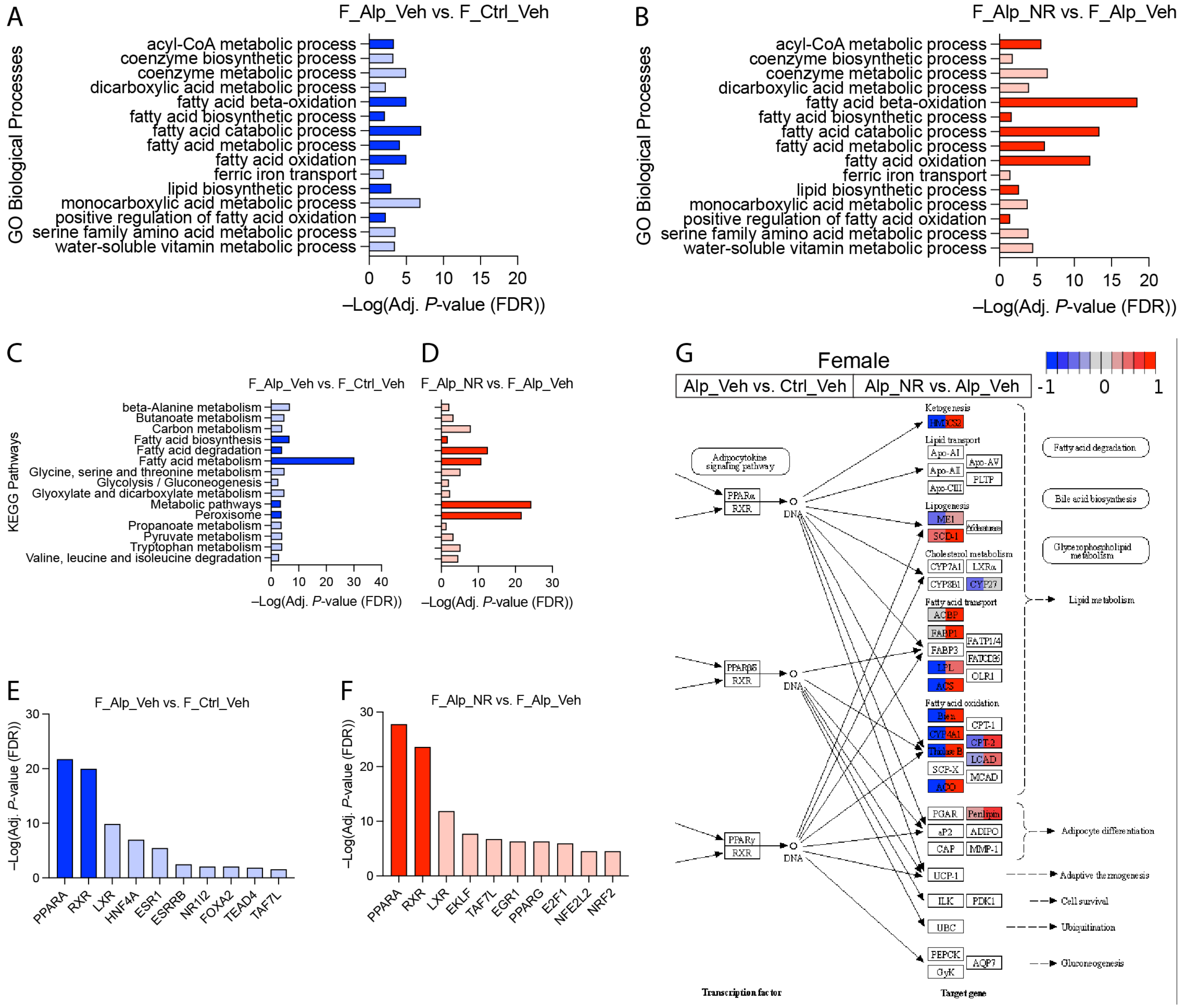

**Figure S8: NAD+ supplementation restores renal metabolism in female Alport mice.** Bulk kidney cortex RNA-seq data from control and Alport mice, treated with or without NR, were analyzed (*N* = 4 mice per group). (**A,B**) Gene ontology (GO) biological processes that are both (*A*) reduced in vehicle-treated female Alport mice (vs. vehicle-treated female control mice) and (*B*) increased in NR-treated female Alport mice (vs. vehicle-treated female Alport mice) are shown. (**C,D**) KEGG pathways that are both (*C*) reduced in vehicle-treated female Alport mice (vs. vehicle-treated female control mice) and (*D*) increased in NR-treated female Alport mice (vs. vehicle-treated female Alport mice) are shown. (**E,F**) Transcription factor analyses suggest that the RXR/PPARα gene regulatory network is (*E*) inhibited in vehicle-treated female Alport mice (vs. vehicle-treated female control mice) and (*F*) activated in NR-treated female Alport mice (vs. vehicle-treated female Alport mice). Processes (*A*,*B*), pathways (*C*,*D*), and transcription factors (*E*,*F*) that are directly involved in fatty acid metabolism are highly enriched in all comparisons, emphasized by either dark blue (decreased) or dark red (increased). (**G**) KEGG graph for the PPAR signaling pathway (KEGG Entry No. 03320). The left and right sides of each gene box represent data from vehicle-treated female Alport mice (vs. vehicle-treated female control mice) and NR-treated female Alport mice (vs. vehicle-treated female Alport mice), respectively. The sub-pathway that was the most restored by NR treatment was fatty acid oxidation. Alp, Alport; Ctrl, control; F, female; NR, nicotinamide riboside; Veh, vehicle.

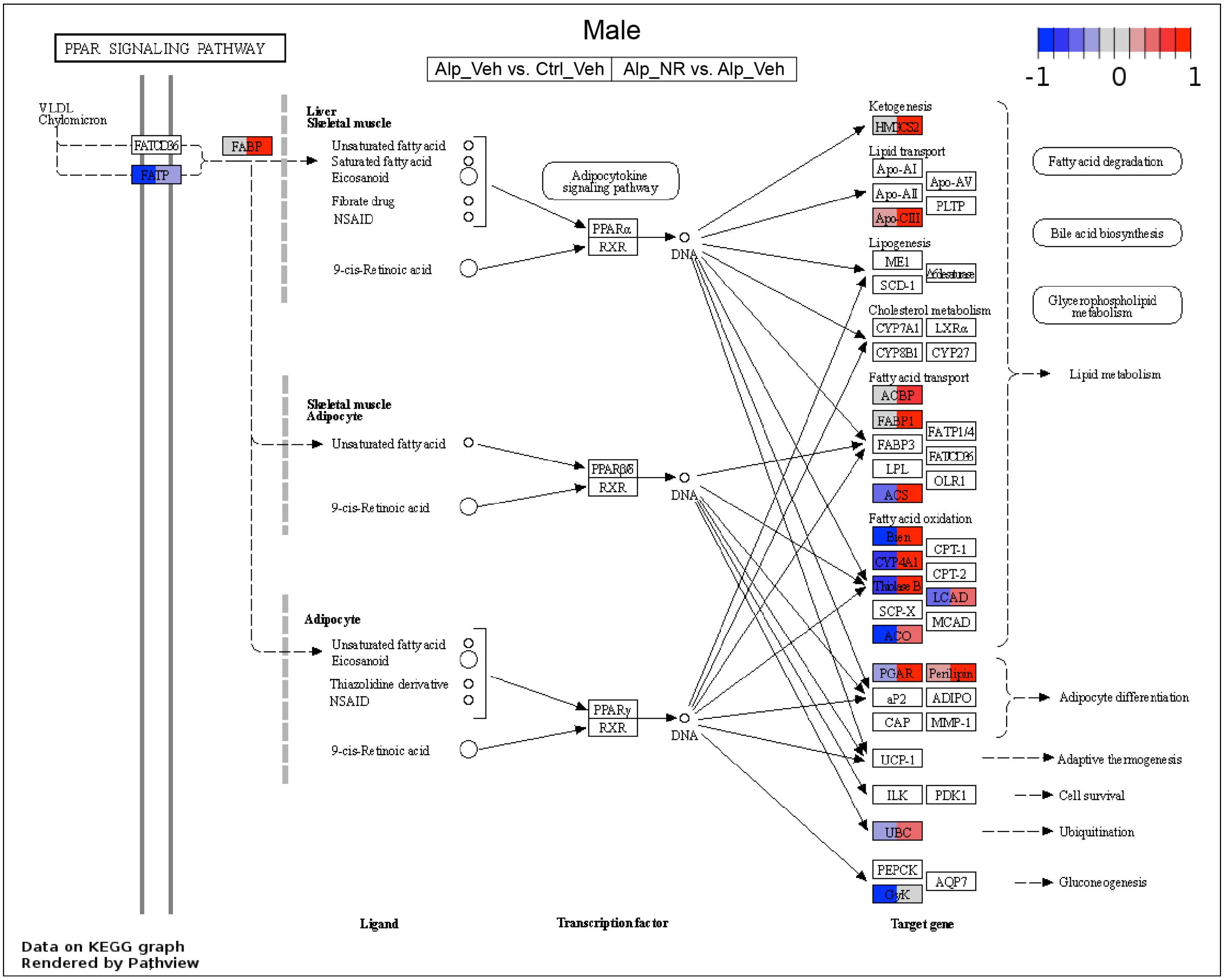

**Figure S9: NAD+ supplementation activates RXR/PPAR signaling in male Alport mice.** Bulk kidney cortex RNA-seq data from control and Alport mice, treated with or without NR, were analyzed (*N* = 4 mice per group). A KEGG graph was plotted for the PPAR signaling pathway (KEGG Entry No. 03320). The left and right sides of each gene box represent data from vehicle-treated male Alport mice (vs. vehicle-treated male control mice) and NR-treated male Alport mice (vs. vehicle-treated male Alport mice), respectively. The PPAR signaling pathway was impaired in vehicle-treated male Alport mice (*left side of boxes*), and this was reversed by NR treatment (*right side of boxes*). The PPAR-regulated downstream pathways that were most affected were those of fatty acid oxidation and fatty acid transport. The right portion of this KEGG graph is presented alongside the male RNA-seq data in the *Results*. The full KEGG graph is presented here. Alp, Alport; Ctrl, control; M, male; NR, nicotinamide riboside; Veh, vehicle.

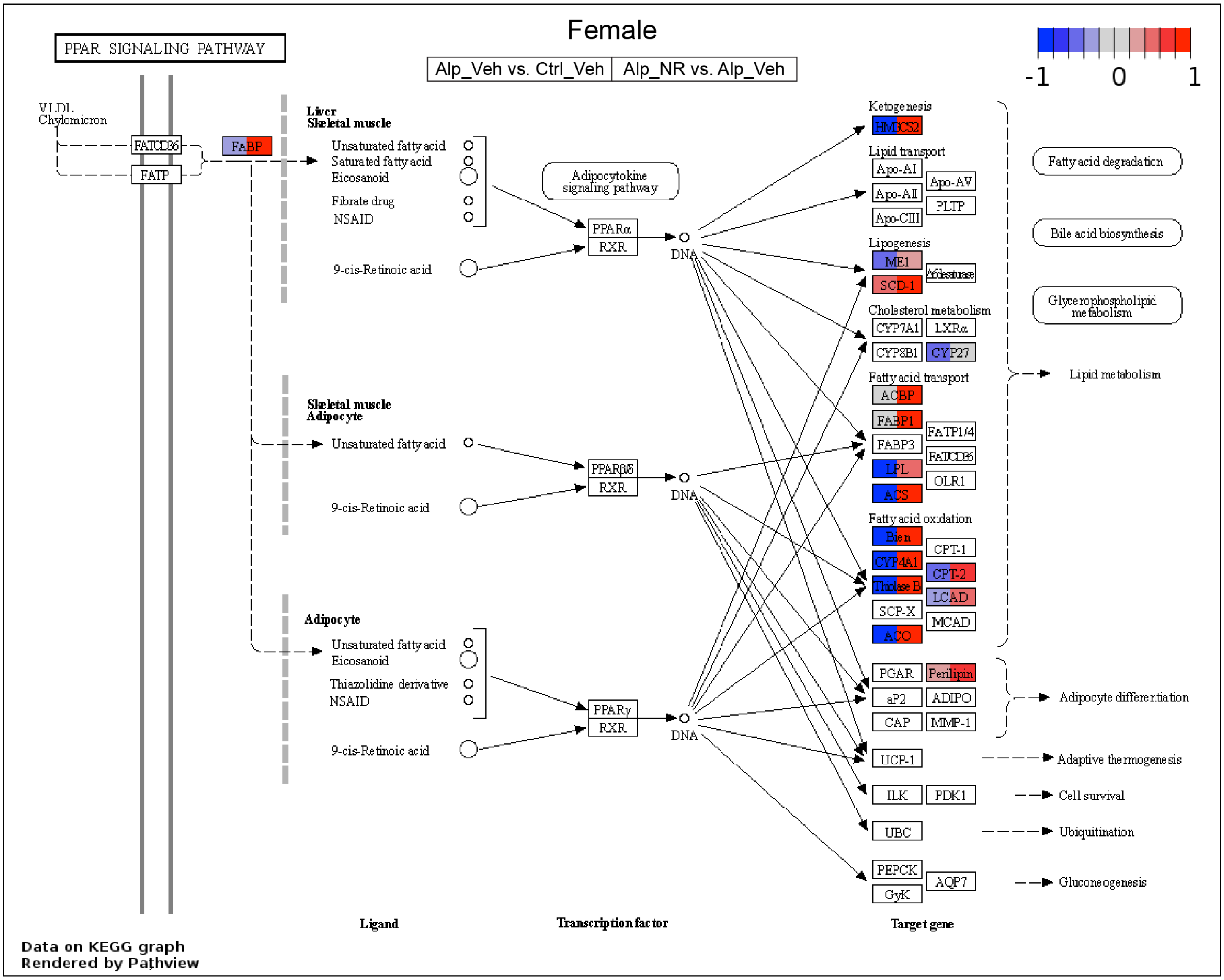

**Figure S10: NAD$^+$ supplementation activates RXR/PPAR signaling in female Alport mice.** Bulk kidney cortex RNA-seq data from control and Alport mice, treated with or without NR, were analyzed (*N* = 4 mice per group). A KEGG graph was plotted for the PPAR signaling pathway (KEGG Entry No. 03320). The left and right sides of each gene box represent data from vehicle-treated female Alport mice (vs. vehicle-treated female control mice) and NR-treated female Alport mice (vs. vehicle-treated female Alport mice), respectively. The PPAR signaling pathway was impaired in vehicle-treated female Alport mice (*left side of boxes*), and this was reversed by NR treatment (*right side of boxes*). The PPAR-regulated downstream pathways that were most affected were those of fatty acid oxidation and fatty acid transport. The right portion of this KEGG graph is presented alongside the female RNA-seq data in the *Supplemental Material*. The full KEGG graph is presented here. Alp, Alport; Ctrl, control; F, female; NR, nicotinamide riboside; Veh, vehicle.

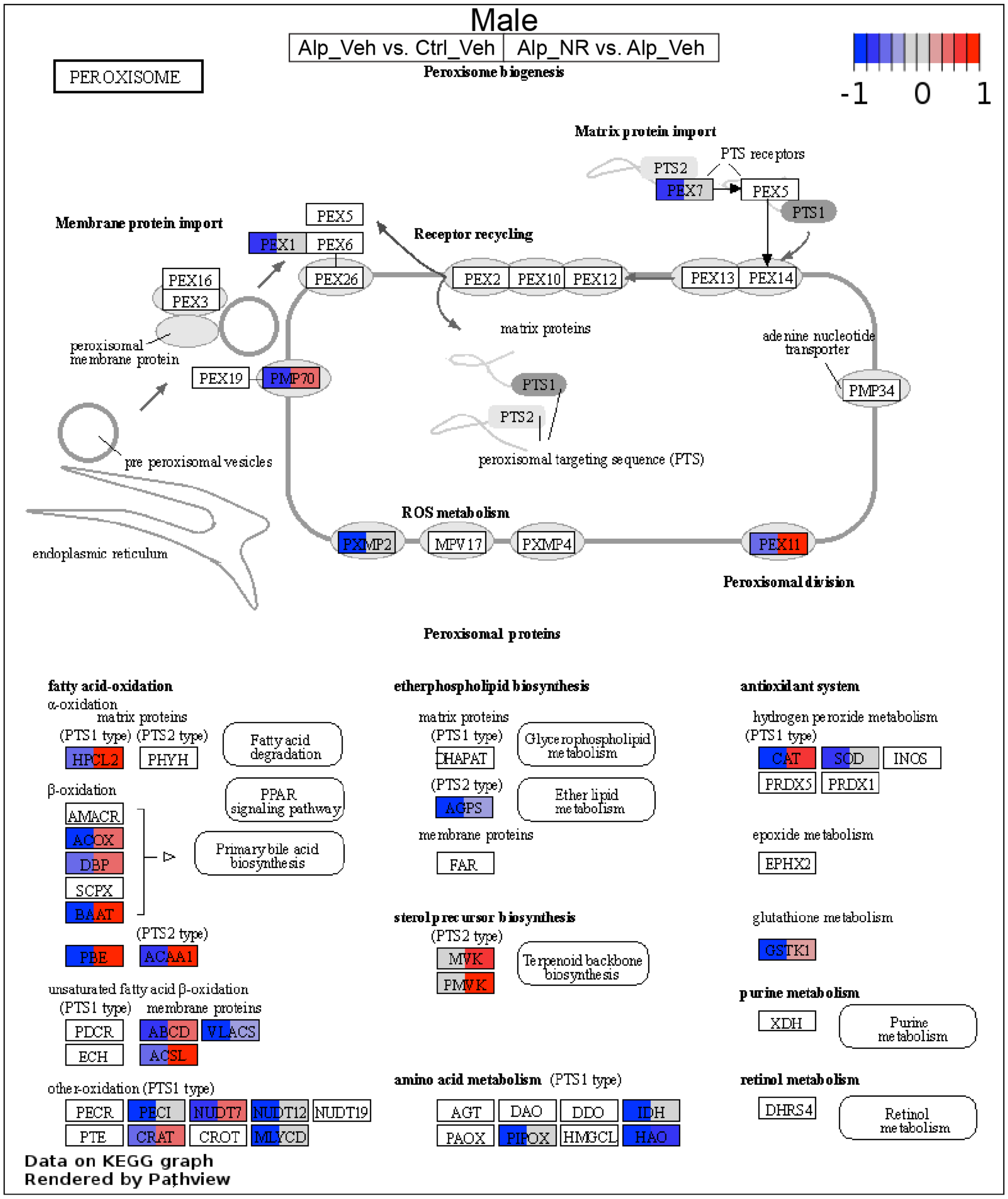

**Figure S11: NAD⁺ supplementation activates peroxisome pathways in male Alport mice.** Bulk kidney cortex RNA-seq data from control and Alport mice, treated with or without NR, were analyzed ($N$ = 4 mice per group). A KEGG graph was plotted for the peroxisome (KEGG Entry No. 04146). The left and right sides of each gene box represent data from vehicle-treated male Alport mice (vs. vehicle-treated male control mice) and NR-treated male Alport mice (vs. vehicle-treated male Alport mice), respectively. The fatty acid oxidation sub-pathway was greatly impaired in vehicle-treated male Alport mice (*left side of boxes*), and this was reversed by NR treatment (*right side of boxes*). Alp, Alport; Ctrl, control; M, male; NR, nicotinamide riboside; Veh, vehicle.

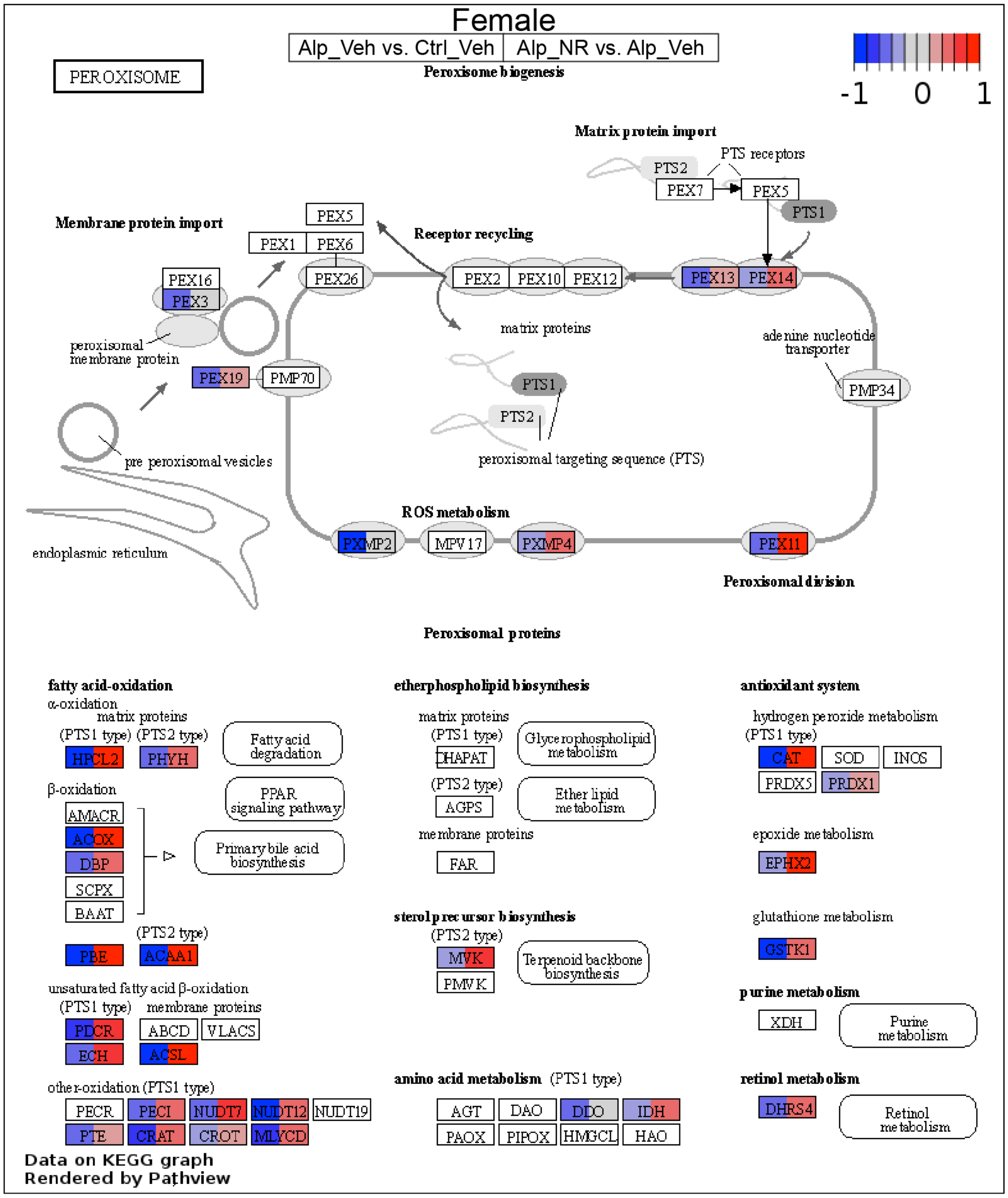

**Figure S12: NAD+ supplementation activates peroxisome pathways in female Alport mice.** Bulk kidney cortex RNA-seq data from control and Alport mice, treated with or without NR, were analyzed (*N* = 4 mice per group). A KEGG graph was plotted for the peroxisome (KEGG Entry No. 04146). The left and right sides of each gene box represent data from vehicle-treated female Alport mice (vs. vehicle-treated female control mice) and NR-treated female Alport mice (vs. vehicle-treated female Alport mice), respectively. The fatty acid oxidation sub-pathway was greatly impaired in vehicle-treated female Alport mice (*left side of boxes*), and this was reversed by NR treatment (*right side of boxes*). Alp, Alport; Ctrl, control; F, female; NR, nicotinamide riboside; Veh, vehicle.

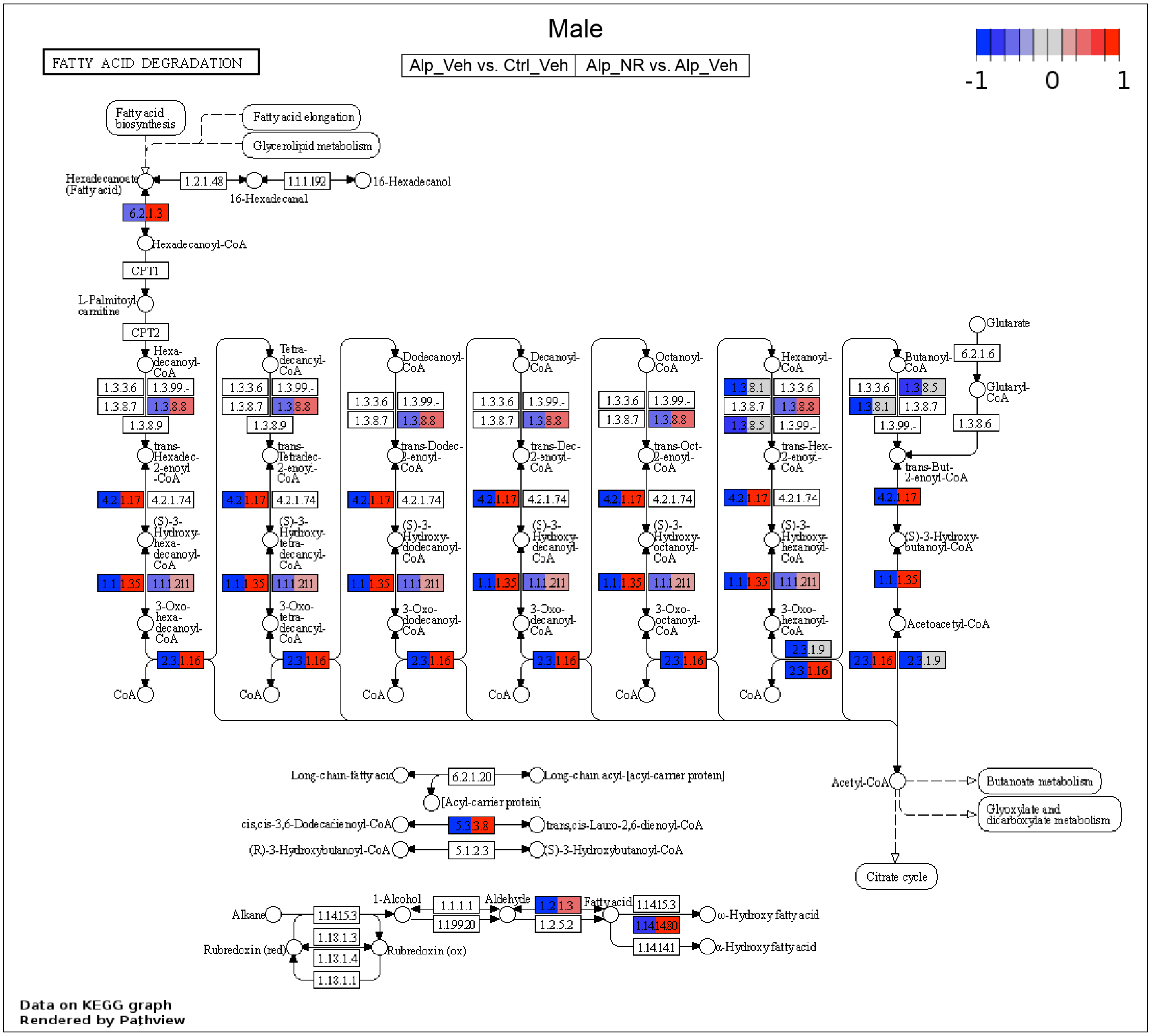

**Figure S13: NAD+ supplementation activates fatty acid degradation in male Alport mice.** Bulk kidney cortex RNA-seq data from control and Alport mice, treated with or without NR, were analyzed ($N$ = 4 mice per group). A KEGG graph was plotted for the fatty acid degradation pathway (KEGG Entry No. 00071). The left and right sides of each gene box represent data from vehicle-treated male Alport mice (vs. vehicle-treated male control mice) and NR-treated male Alport mice (vs. vehicle-treated male Alport mice), respectively. Fatty acid degradation was greatly impaired in male Alport mice (*left side of boxes*), and this was reversed by NR treatment (*right side of boxes*). Alp, Alport; Ctrl, control; M, male; NR, nicotinamide riboside; Veh, vehicle.

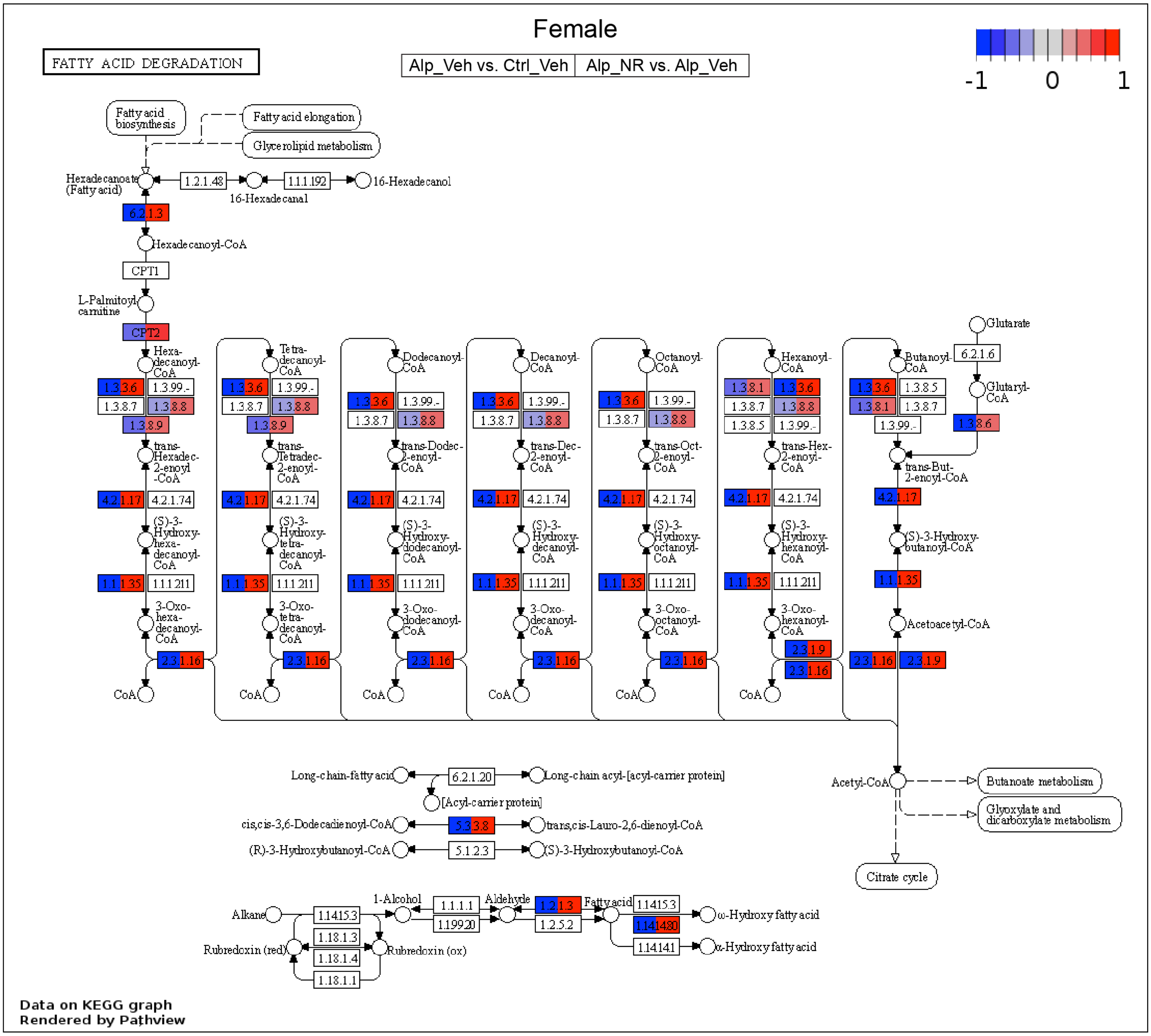

**Figure S14: NAD⁺ supplementation activates fatty acid degradation in female Alport mice.** Bulk kidney cortex RNA-seq data from control and Alport mice, treated with or without NR, were analyzed (*N* = 4 mice per group). A KEGG graph was plotted for the fatty acid degradation pathway (KEGG Entry No. 00071). The left and right sides of each gene box represent data from vehicle-treated female Alport mice (vs. vehicle-treated female control mice) and NR-treated female Alport mice (vs. vehicle-treated female Alport mice), respectively. Fatty acid degradation was greatly impaired in female Alport mice (*left side of boxes*), and this was reversed by NR treatment (*right side of boxes*). Alp, Alport; Ctrl, control; F, female; NR, nicotinamide riboside; Veh, vehicle.

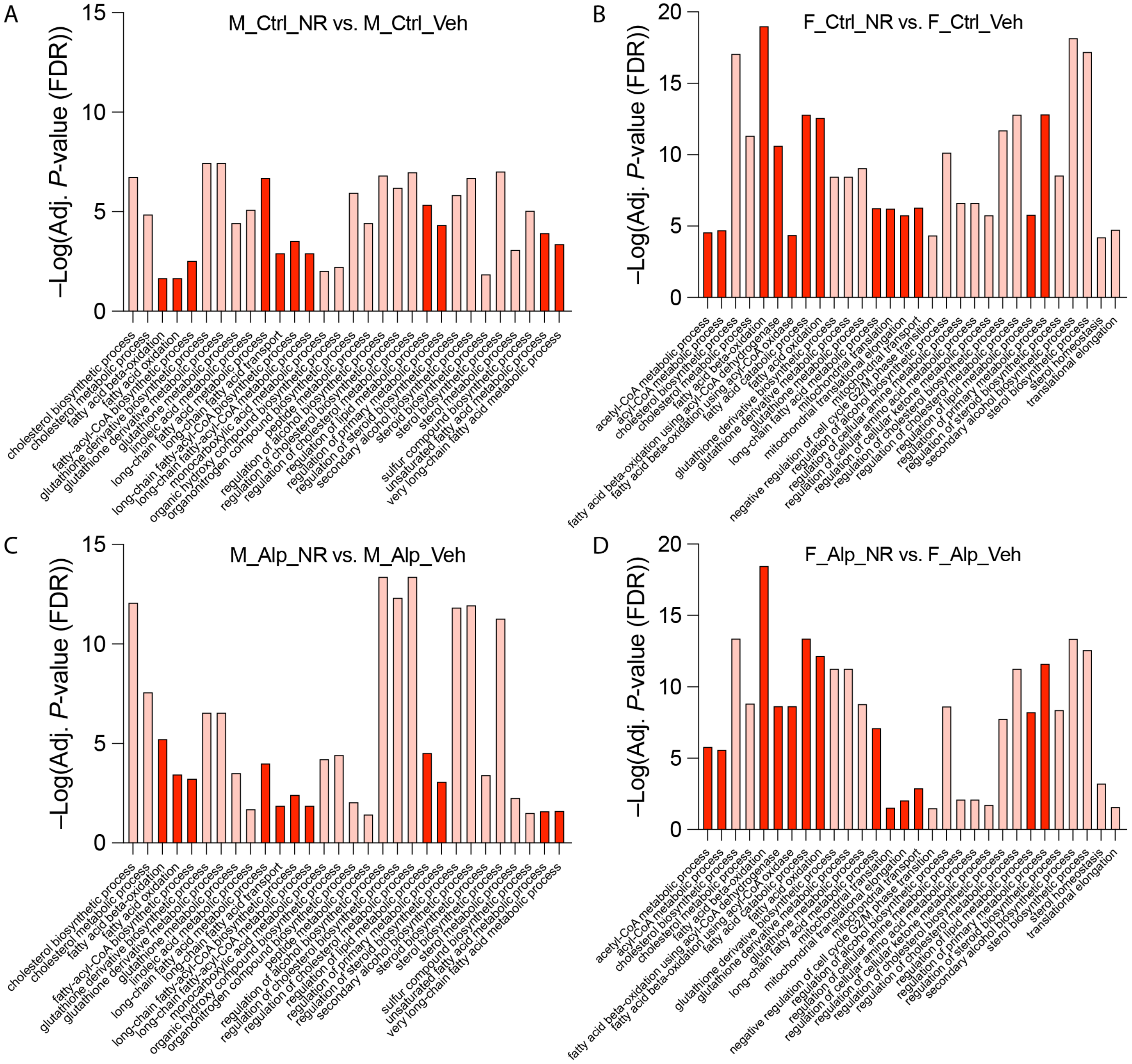

**Figure S15: NAD+ supplementation activates metabolic biological processes in control mice, not just in Alport mice.** Bulk kidney cortex RNA-seq data from control and Alport mice, treated with or without NR, were analyzed ($N$ = 4 mice per group). Gene ontology (GO) biological processes that were simultaneously upregulated in both NR-treated control mice (vs. vehicle-treated control mice) and in NR-treated Alport mice (vs. vehicle-treated Alport mice) were identified. (**A,B**) GO biological processes that were increased in NR-treated control mice (vs. vehicle-treated control mice) in males (*A*) and females (*B*). (**C,D**) GO biological processes that were increased in NR-treated Alport mice (vs. vehicle-treated Alport mice) in males (*C*) and females (*D*). Data in (*A*) and (*B*) were presented in the *Results*, and, for clarity, they are presented adjacent to their corresponding sex-match comparisons from NR-treated Alport mice (vs. vehicle-treated Alport mice). Processes that are directly involved in fatty acid metabolism are highly enriched in all comparisons, emphasized by dark red (increased). Alp, Alport; Ctrl, control; F, female; M, male; NR, nicotinamide riboside; Veh, vehicle.

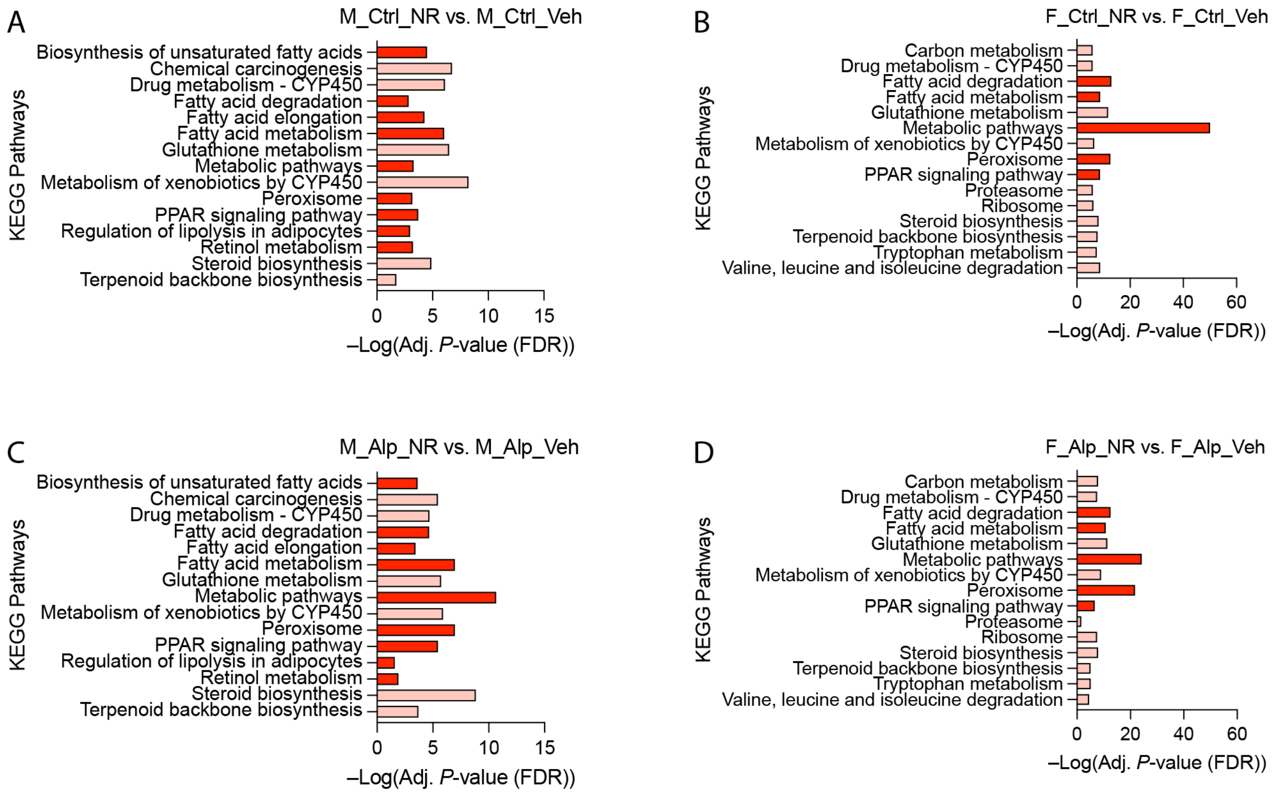

**Figure S16: NAD⁺ supplementation activates metabolic pathways in control mice, not just in Alport mice.** Bulk kidney cortex RNA-seq data from control and Alport mice, treated with or without NR, were analyzed ($N$ = 4 mice per group). (**A,B**) KEGG pathways that were simultaneously upregulated in both NR-treated control mice (vs. vehicle-treated control mice) and in NR-treated Alport mice (vs. vehicle-treated Alport mice) were identified. (**A,B**) KEGG pathways that were increased in NR-treated control mice (vs. vehicle-treated control mice) in males (*A*) and females (*B*). (**C,D**) KEGG pathways that were increased in NR-treated Alport mice (vs. vehicle-treated Alport mice) in males (*C*) and females (*D*). Data in (*A*) and (*B*) were presented in the *Results*, and, for clarity, they are presented adjacent to their corresponding sex-match comparisons from NR-treated Alport mice (vs. vehicle-treated Alport mice). Pathways that are directly involved in fatty acid metabolism are highly enriched in all comparisons, emphasized by dark red (increased). Alp, Alport; Ctrl, control; F, female; M, male; NR, nicotinamide riboside; Veh, vehicle.

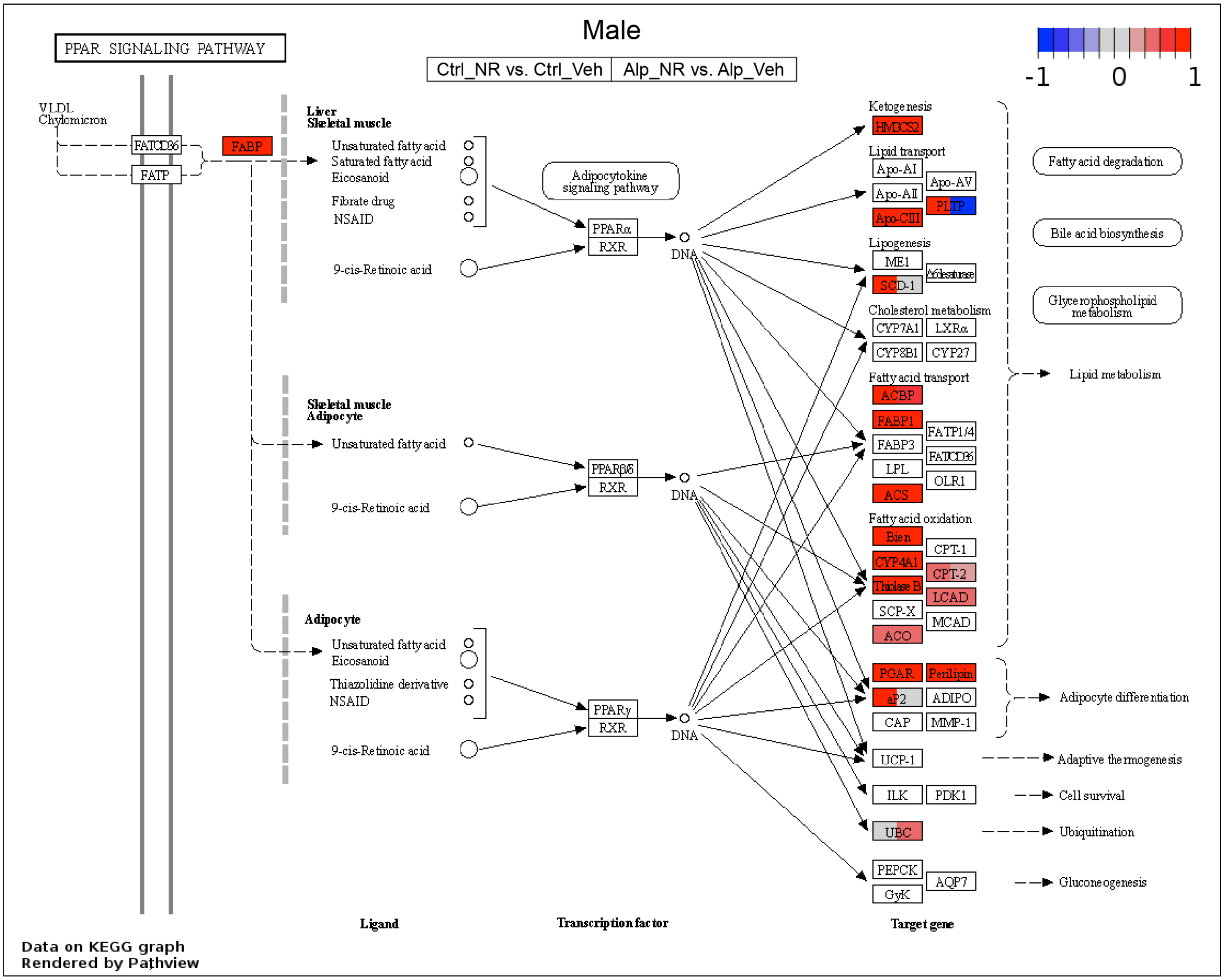

**Figure S17: NAD+ supplementation activates RXR/PPAR signaling in male control mice, not just Alport mice.** Bulk kidney cortex RNA-seq data from control and Alport mice, treated with or without NR, were analyzed (*N* = 4 mice per group). A KEGG graph was plotted for the PPAR signaling pathway (KEGG Entry No. 03320). The left and right sides of each gene box represent data from NR-treated male control mice (vs. vehicle-treated male control mice) and NR-treated male Alport mice (vs. vehicle-treated male Alport mice), respectively. NR treatment activated the PPAR signaling pathway, especially fatty acid oxidation, in both healthy male control mice (*left side of boxes*) and diseased male Alport mice (*right side of boxes*). Alp, Alport; Ctrl, control; M, male; NR, nicotinamide riboside; Veh, vehicle.

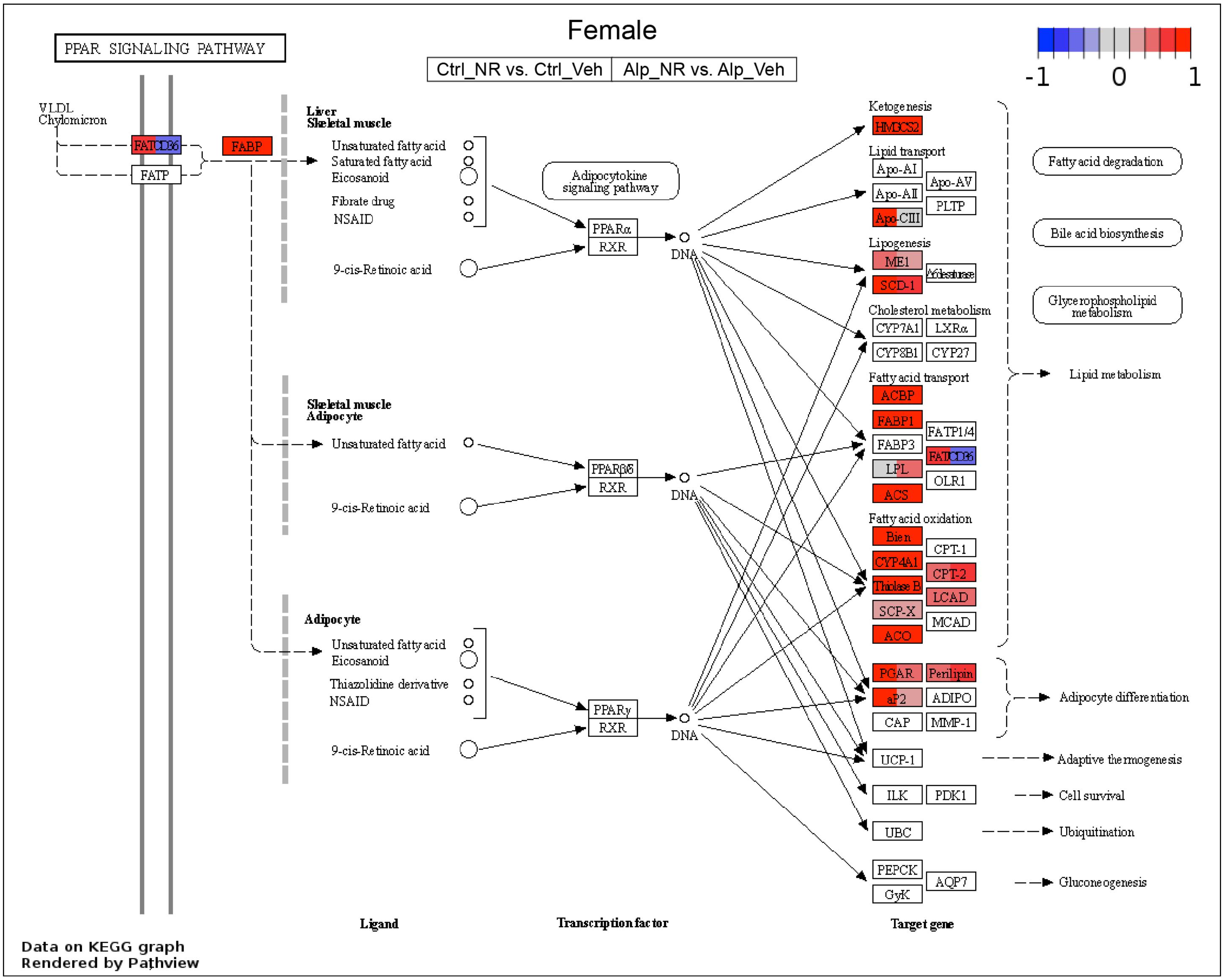

**Figure S18: NAD+ supplementation activates RXR/PPAR signaling in female control mice, not just Alport mice.** Bulk kidney cortex RNA-seq data from control and Alport mice, treated with or without NR, were analyzed (*N* = 4 mice per group). A KEGG graph was plotted for the PPAR signaling pathway (KEGG Entry No. 03320). The left and right sides of each gene box represent data from NR-treated female control mice (vs. vehicle-treated female control mice) and NR-treated female Alport mice (vs. vehicle-treated female Alport mice), respectively. NR treatment activated the PPAR signaling pathway, especially fatty acid oxidation, in both healthy female control mice (*left side of boxes*) and diseased female Alport mice (*right side of boxes*). Alp, Alport; Ctrl, control; F, female; NR, nicotinamide riboside; Veh, vehicle.

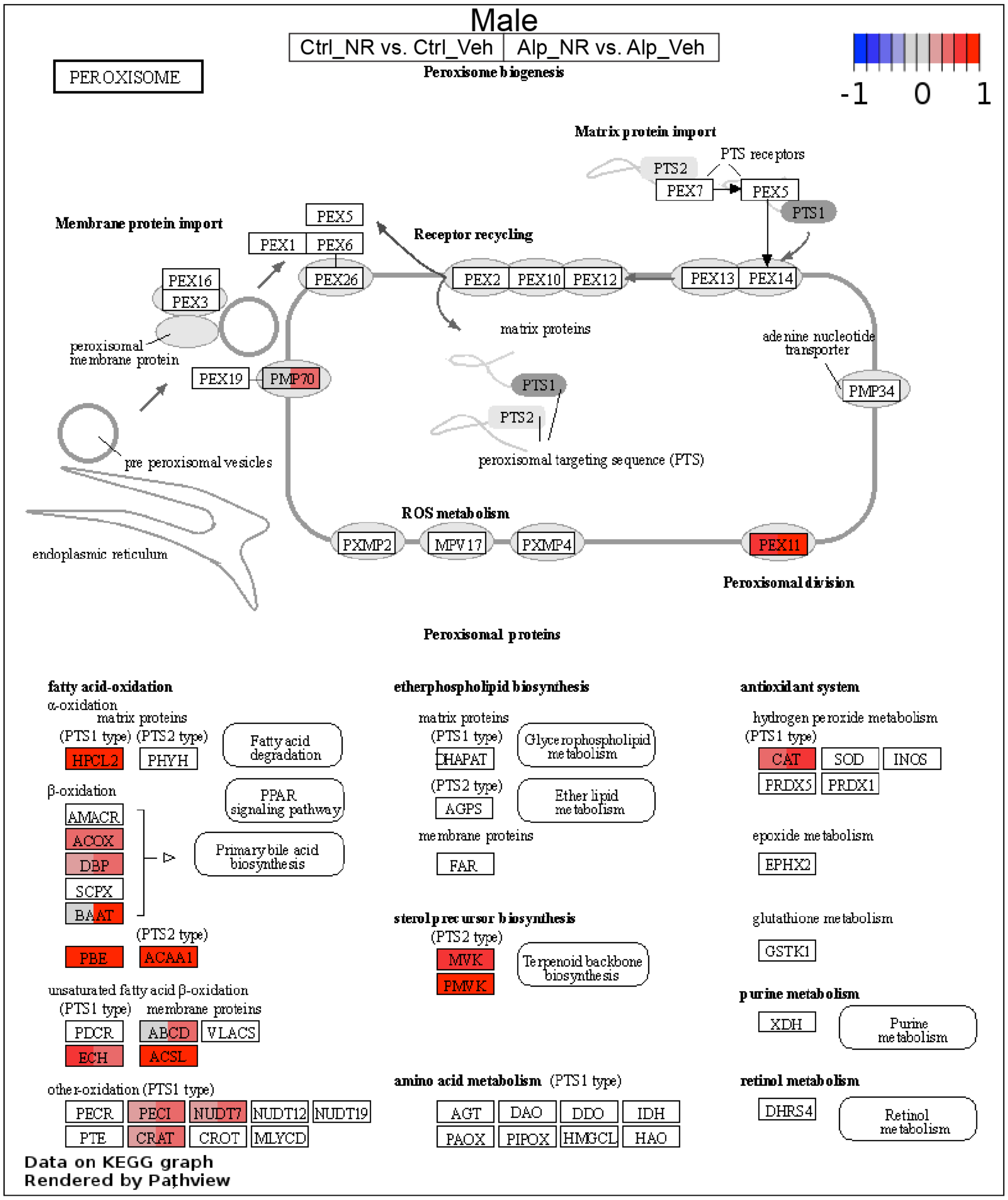

**Figure S19: NAD+ supplementation activates the peroxisome in male control mice, not just Alport mice.** Bulk kidney cortex RNA-seq data from control and Alport mice, treated with or without NR, were analyzed ($N$ = 4 mice per group). A KEGG graph was plotted for the peroxisome (KEGG Entry No. 04146). The left and right sides of each gene box represent data from NR-treated male control mice (vs. vehicle-treated male control mice) and NR-treated male Alport mice (vs. vehicle-treated male Alport mice), respectively. NR treatment activated the peroxisome, including fatty acid oxidation, in both healthy male control mice (*left side of boxes*) and diseased male Alport mice (*right side of boxes*). Alp, Alport; Ctrl, control; M, male; NR, nicotinamide riboside; Veh, vehicle.

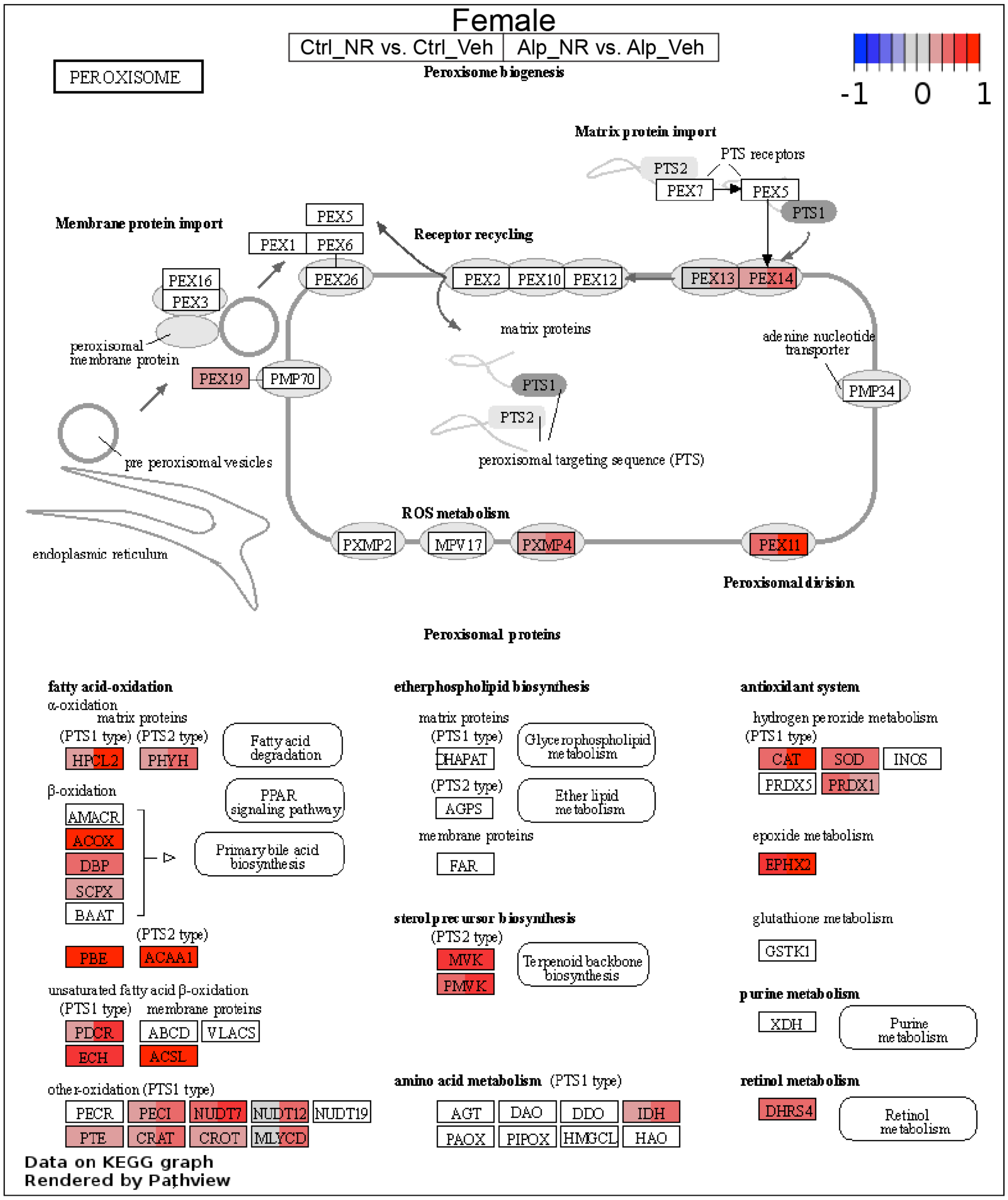

**Figure S20: NAD$^+$ supplementation activates the peroxisome in female control mice, not just Alport mice.** Bulk kidney cortex RNA-seq data from control and Alport mice, treated with or without NR, were analyzed (*N* = 4 mice per group). A KEGG graph was plotted for the peroxisome (KEGG Entry No. 04146). The left and right sides of each gene box represent data from NR-treated female control mice (vs. vehicle-treated female control mice) and NR-treated female Alport mice (vs. vehicle-treated female Alport mice), respectively. NR treatment activated the peroxisome, including fatty acid oxidation, in both healthy female control mice (*left side of boxes*) and diseased female Alport mice (*right side of boxes*). Alp, Alport; Ctrl, control; F, female; NR, nicotinamide riboside; Veh, vehicle.

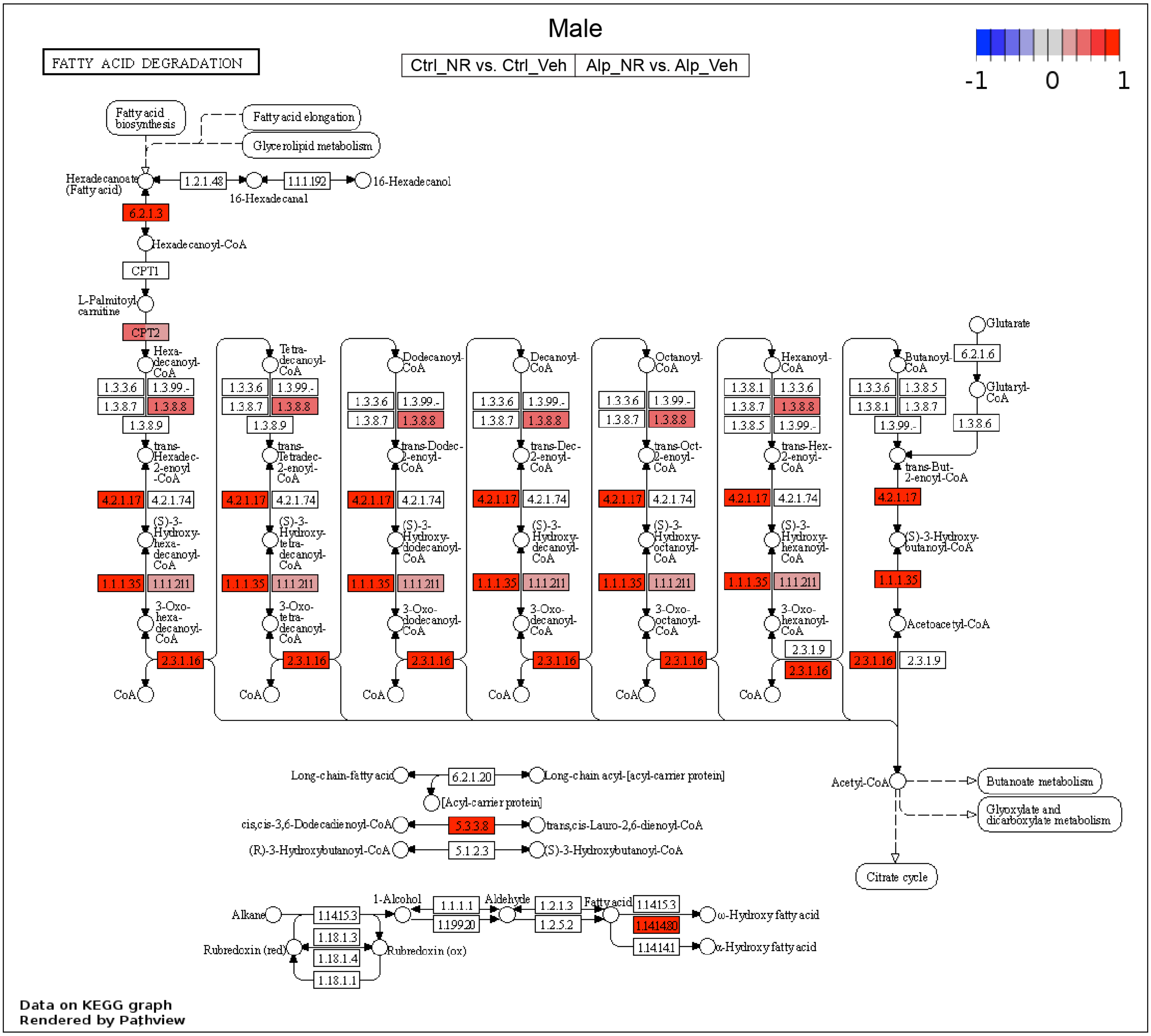

**Figure S21: NAD+ supplementation activates fatty acid degradation in male control mice, not just Alport mice.** Bulk kidney cortex RNA-seq data from control and Alport mice, treated with or without NR, were analyzed (*N* = 4 mice per group). A KEGG graph was plotted for the fatty acid degradation pathway (KEGG Entry No. 00071). The left and right sides of each gene box represent data from NR-treated male control mice (vs. vehicle-treated male control mice) and NR-treated male Alport mice (vs. vehicle-treated male Alport mice), respectively. NR treatment activated fatty acid degradation in both healthy male control mice (*left side of boxes*) and diseased male Alport mice (*right side of boxes*). Alp, Alport; Ctrl, control; M, male; NR, nicotinamide riboside; Veh, vehicle.

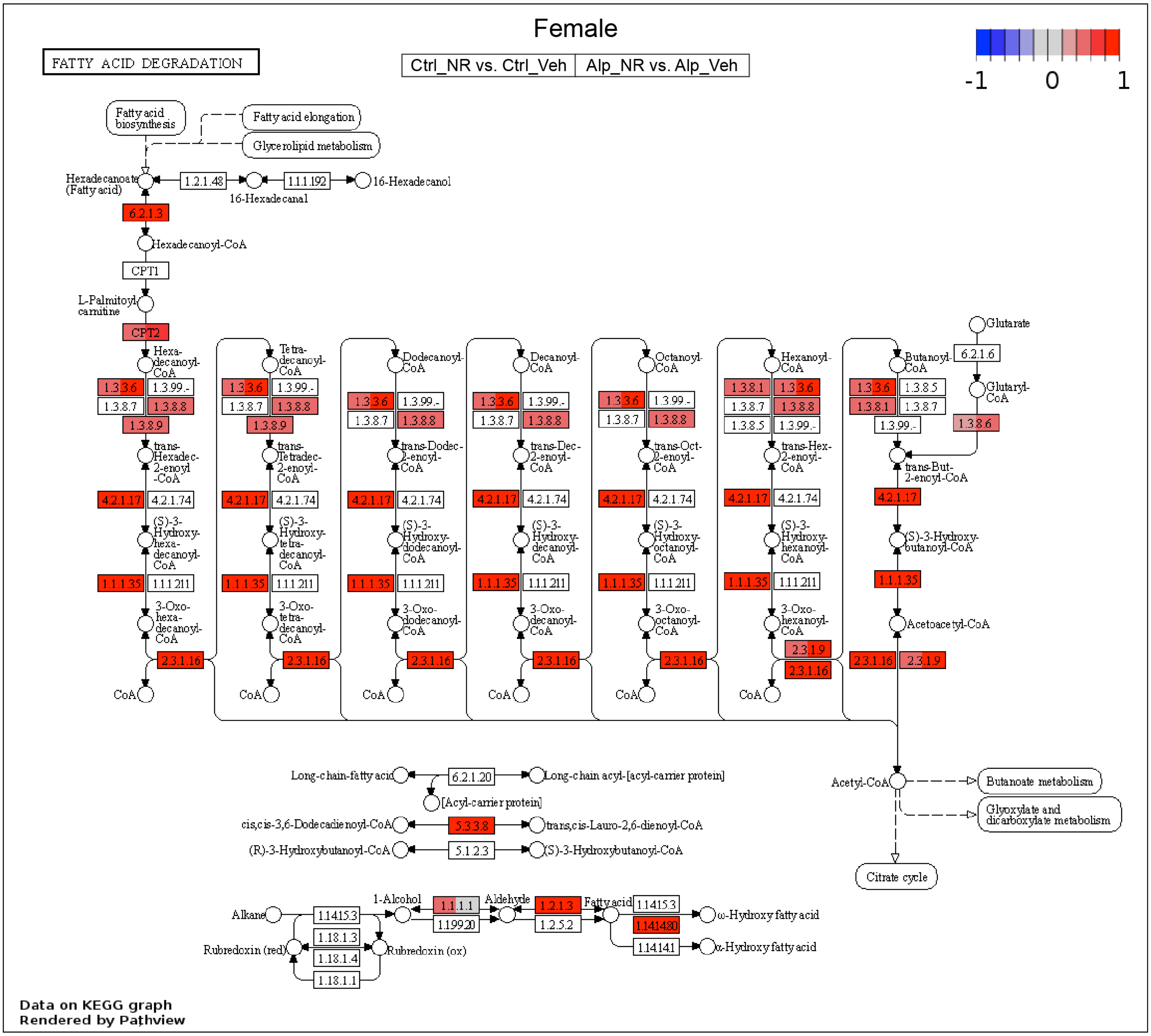

**Figure S22: NAD+ supplementation activates fatty acid degradation in female control mice, not just Alport mice.** Bulk kidney cortex RNA-seq data from control and Alport mice, treated with or without NR, were analyzed (*N* = 4 mice per group). A KEGG graph was plotted for the fatty acid degradation pathway (KEGG Entry No. 00071). The left and right sides of each gene box represent data from NR-treated female control mice (vs. vehicle-treated female control mice) and NR-treated female Alport mice (vs. vehicle-treated female Alport mice), respectively. NR treatment activated fatty acid degradation in both healthy female control mice (*left side of boxes*) and diseased female Alport mice (*right side of boxes*). Alp, Alport; Ctrl, control; F, female; NR, nicotinamide riboside; Veh, vehicle.

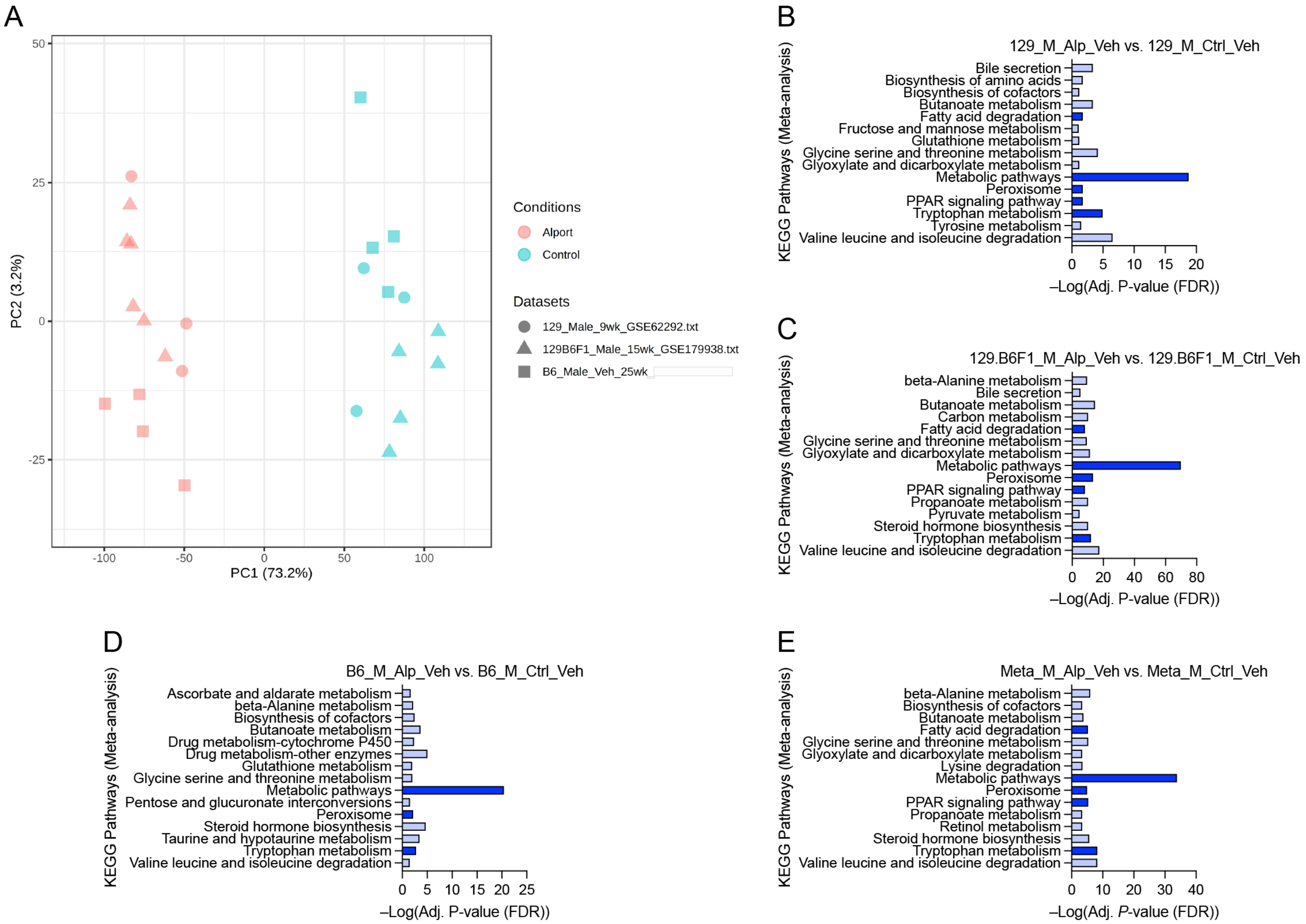

**Figure S23: Alport mice have impaired renal metabolism, irrespective of genetic background.** Bulk kidney cortex RNA-seq data from vehicle-treated male control and Alport mice on the B6 background was compared with previously published kidney RNA-seq data from: 1) male control and Alport mice on the 129 background and 2) a first-generation mixed 129.B6F1 hybrid background. (**A**) Principal component analysis shows good separation between samples by genotype (PC1), but no principal components correlated with genetic background. (**B-D**) KEGG pathway analyses shows that renal metabolic pathways are greatly impaired in Alport mice on the 129X1/SvJ (*B*), 129.B6F1 hybrid (*C*), and C57BL/6J (*D*) backgrounds. (**E**) KEGG pathway analysis of the combined meta-analysis data shows that renal metabolic pathways are greatly impaired in Alport mice, irrespective of 129 or B6 genetic background. Blue indicates a decrease in Alport mice vs. control mice, and important pathways are emphasized by dark blue shading. Alp, Alport; Ctrl, control; M, male; Meta, meta-analysis; PC, principal component; Veh, vehicle; wk, week.

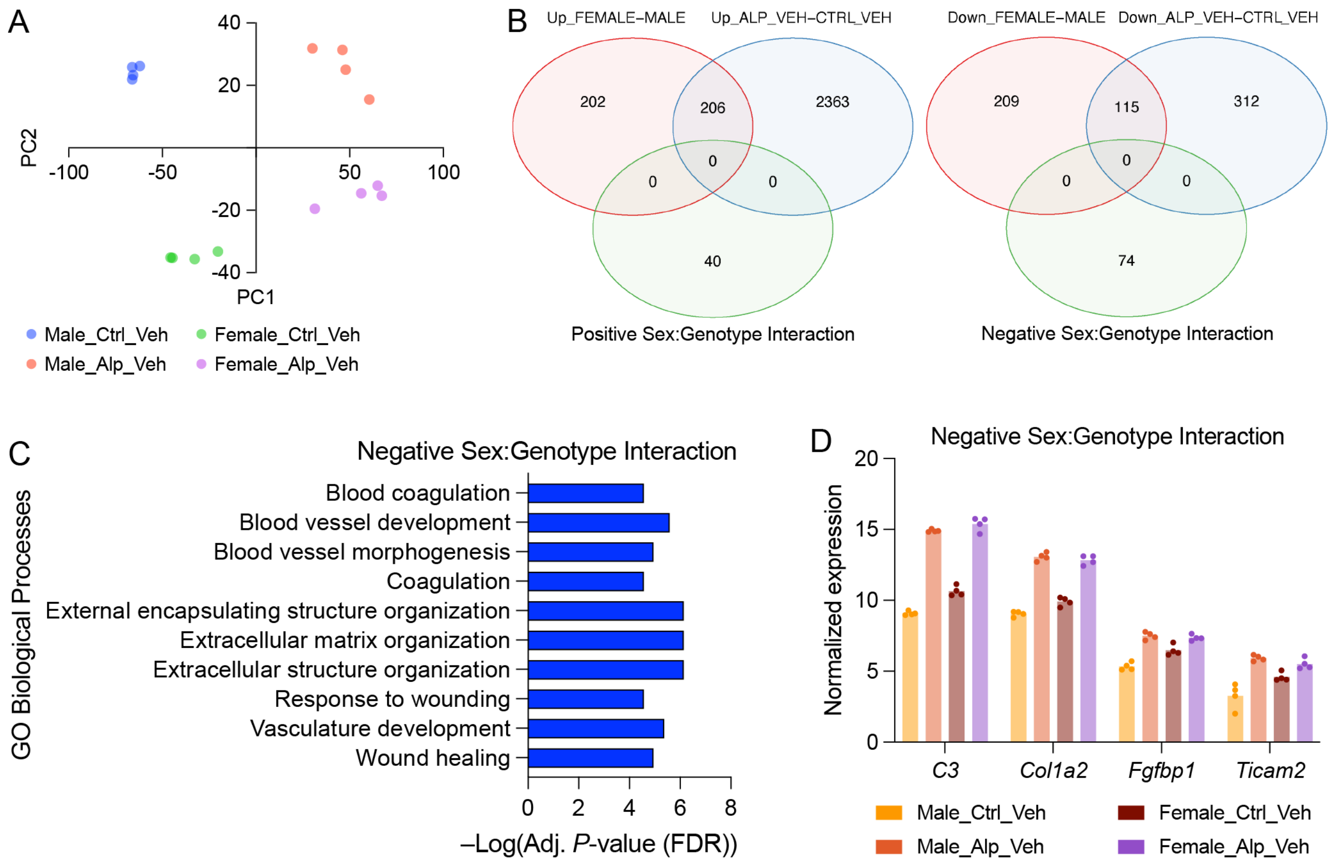

**Figure S24: The *Col4a3*$^{tm1Dec}$ mouse model of Alport syndrome has sex differences at the transcriptional level.** Bulk kidney cortex RNA-seq data from vehicle-treated control and Alport mice were analyzed ($N$ = 4 mice per group). (**A**) Principal component analysis shows good separation between samples by both genotype (PC1) and sex (PC2). (**B**) Venn diagrams showing the number of genes that are differentially regulated between the sexes (*red*) or genotypes (*blue*). Genes with a sex:genotype interaction are shown in green. A positive interaction (*left diagram, green*) indicates that, when comparing genotypes of the same sex, these genes are upregulated (or less downregulated) in the female genotype comparison versus the male genotype comparison. A negative interaction (*right diagram, green*) indicates the reverse of this. (**C**) Gene ontology (GO) biological processes related to fibrosis and neovascularization displayed a significant negative sex:genotype interaction. All statistically significant changes are plotted, and no GO biological processes had a positive sex:genotype interaction. (**D**) Examples of genes related to inflammation (*C3* and *Ticam2*) and fibrosis (*Col1a2* and *Fgfbp1*) that have a negative sex:genotype interaction. This means that, when comparing genotypes of the same sex, these genes are downregulated (or, as in these four cases, less upregulated) in the female genotype comparison versus the male genotype comparison. Each datum represents one mouse. Alp, Alport; Ctrl, control; PC, principal component; Veh, vehicle.

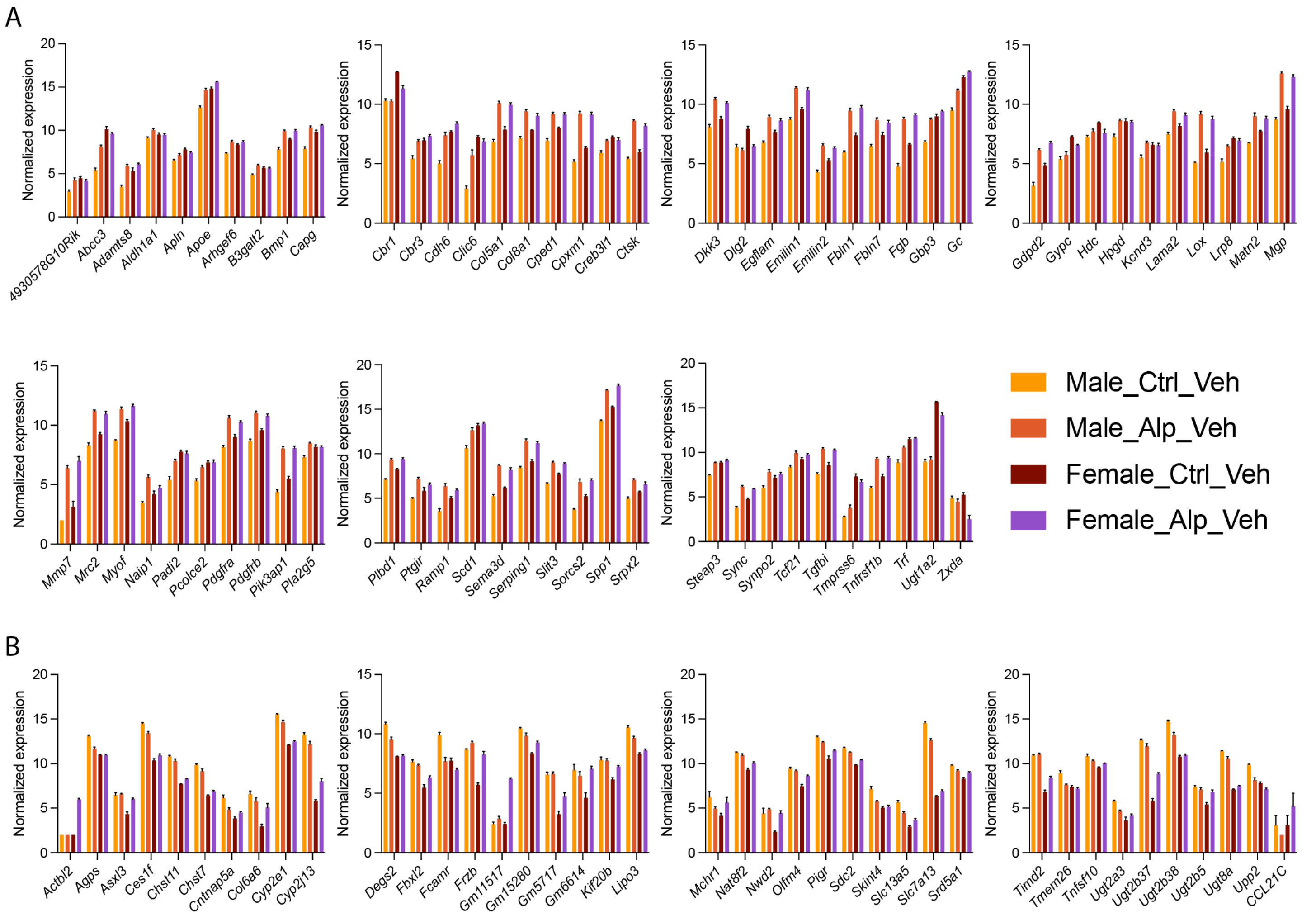

**Figure S25: Genes with a sex:genotype interaction in the *Col4a3*$^{\text{tm1Dec}}$ mouse model of Alport syndrome.** Bulk kidney cortex RNA-seq data from vehicle-treated control and Alport mice were analyzed (*N* = 4 mice per group). (**A**) A total of 74 genes displayed a negative sex:genotype interaction. This means that, when comparing genotypes of the same sex, these genes are either downregulated (e.g., *Abcc3* and *Apln*) or less upregulated (e.g., *Adamts8* and *Apoe*) in the female genotype comparison versus the male genotype comparison. (**B**) A total of 40 genes displayed a positive sex:genotype interaction. This means that, when comparing genotypes of the same sex, these genes are either upregulated (e.g., *Cyp2j13* and *Frzb*) or less downregulated (e.g., *Tmem26* and *Upp2*) in the female genotype comparison versus the male genotype comparison. Alp, Alport; Ctrl, control; Veh, vehicle.

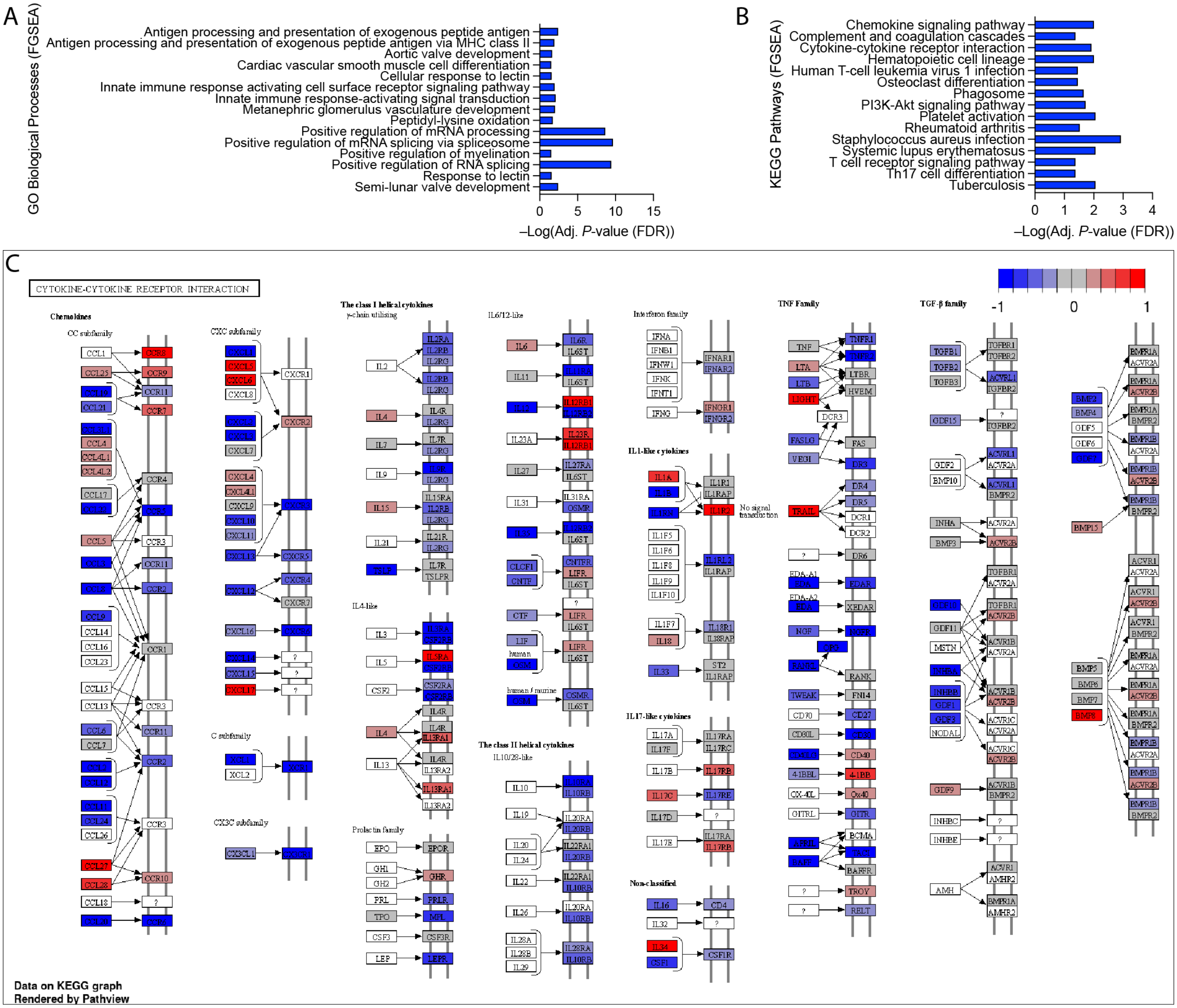

**Figure S26: Females are less susceptible to inflammation- and fibrosis-related transcriptional changes in the *Col4a3*$^{tm1Dec}$ mouse model of Alport syndrome.** Bulk kidney cortex RNA-seq data from vehicle-treated control and Alport mice were analyzed (*N* = 4 mice per group). (**A-B**) Fast gene set enrichment analysis (FGSEA) identified a negative sex:genotype interaction in gene ontology biological process related to inflammation and fibrosis. This means that, when comparing genotypes of the same sex, these biological processes are either downregulated or less upregulated in the female genotype comparison versus the male genotype comparison. (**C**) A KEGG graph was plotted for the cytokine–cytokine receptor interaction pathway (KEGG Entry No. 04060). Red and blue indicate positive and negative sex:genotype interactions, respectively. Most cytokine and cytokine receptors have a negative sex:genotype interaction, indicating less activation in the female genotype comparison versus the male genotype comparison.

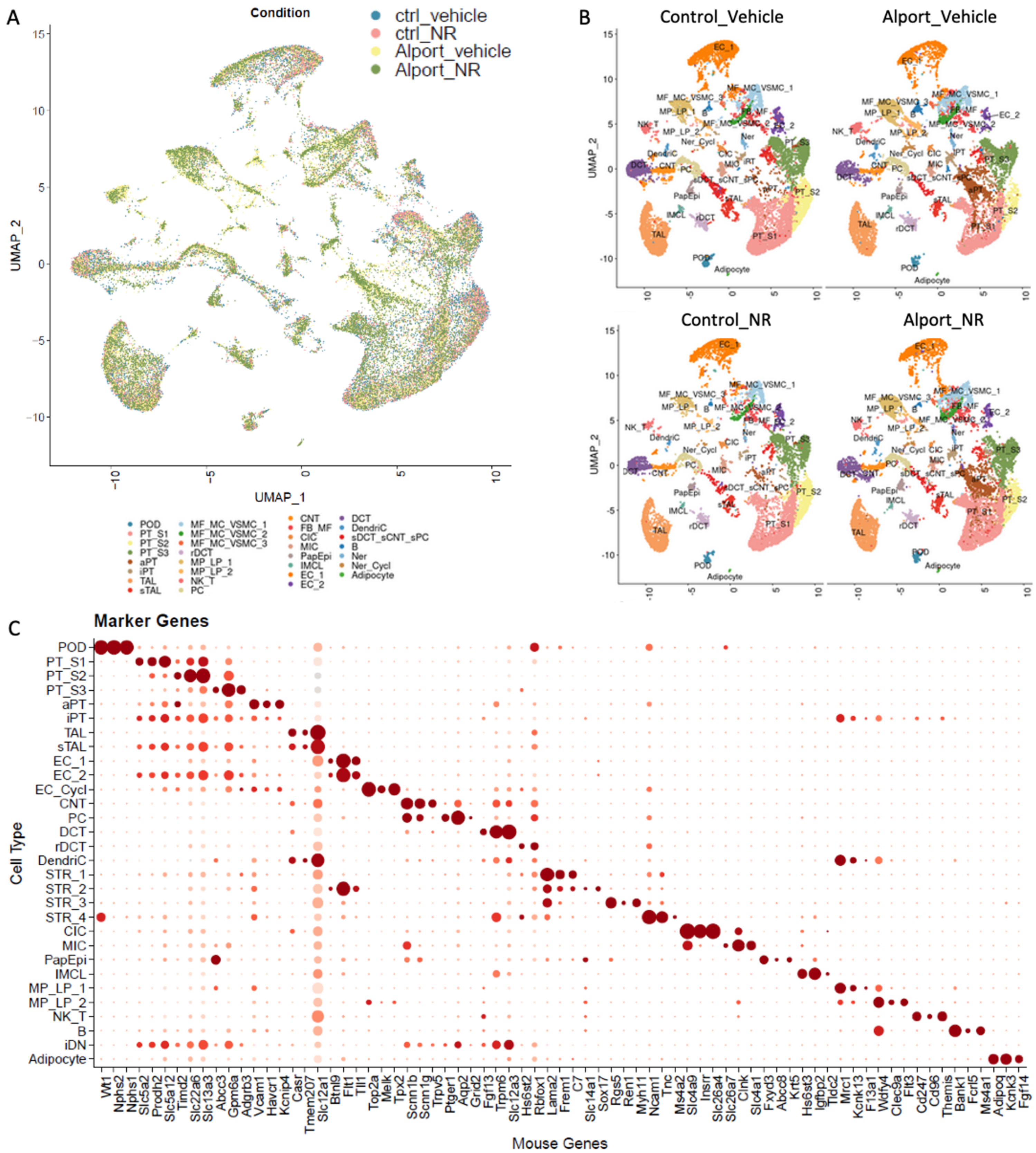

**Figure S27: Single nuclei RNA-sequencing identifies cell types across conditions and cluster markers.** (**A-B**) Uniform manifold approximation and projection reveals the cell type distribution for 4 conditions: control-veh, control-NR, Alport-veh, and Alport-NR. (**C**) Marker genes defining cell type cluster identities.

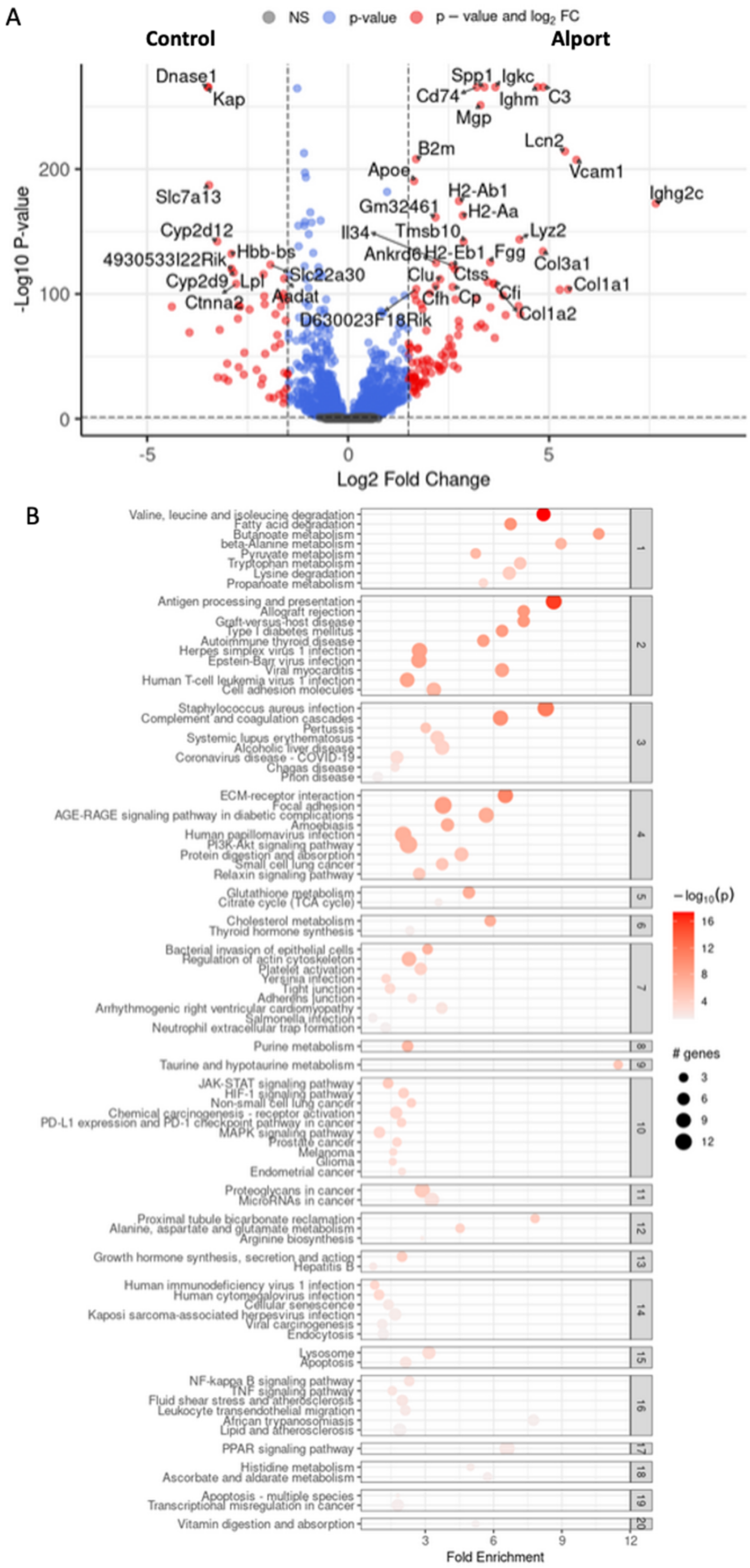

**Figure S28: Single nuclei RNA-sequencing identifies proximal tubule transcriptional differences between vehicle-treated Alport and vehicle-treated control mice.** (**A-B**) Differentially expressed genes (*A*) and pathways (*B*) of the proximal tubule in Alport mice as compared to control mice.

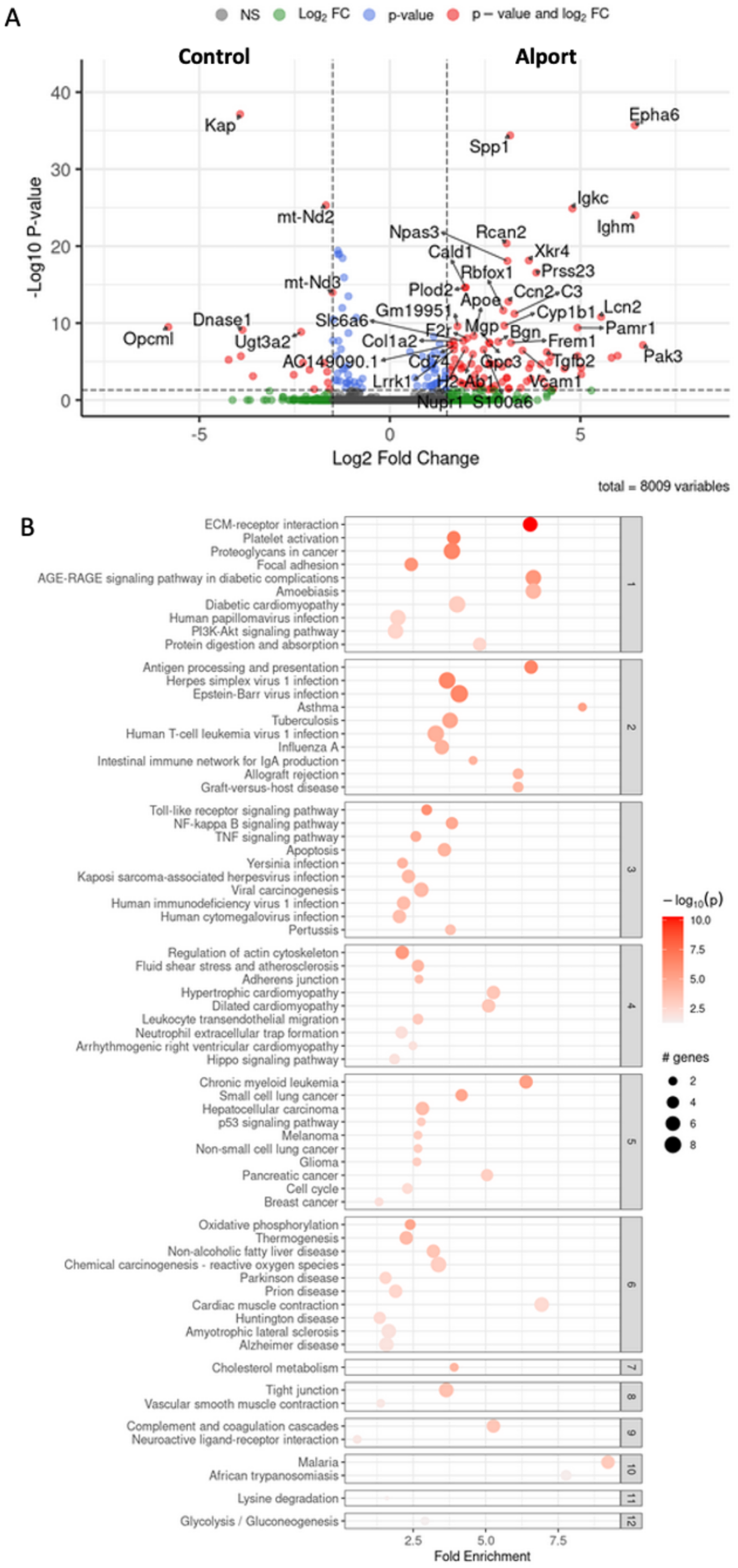

**Figure S29: Single nuclei RNA-sequencing identifies podocyte transcriptional differences between vehicle-treated Alport and vehicle-treated control mice.** (**A-B**) Differentially expressed genes (*A*) and pathways (*B*) of the podocytes in Alport mice as compared to control mice.

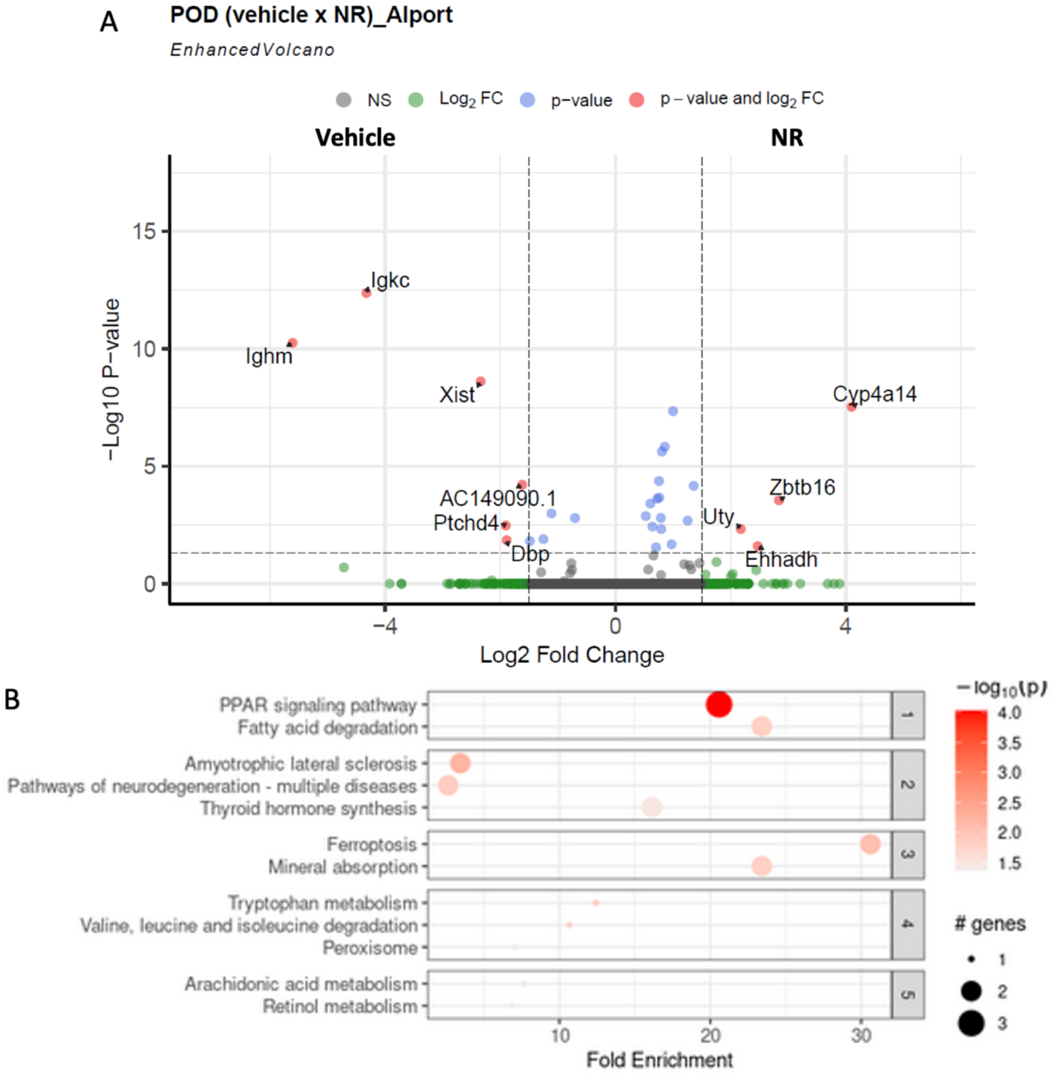

**Figure S30: Single nuclei RNA-sequencing identifies podocyte transcriptional differences between NR-treated Alport mice and vehicle-treated Alport mice.** (**A-B**) Differentially expressed genes (A) and pathways (B) of the podocyte in NR-treated Alport mice as compared to vehicle-treated Alport mice.

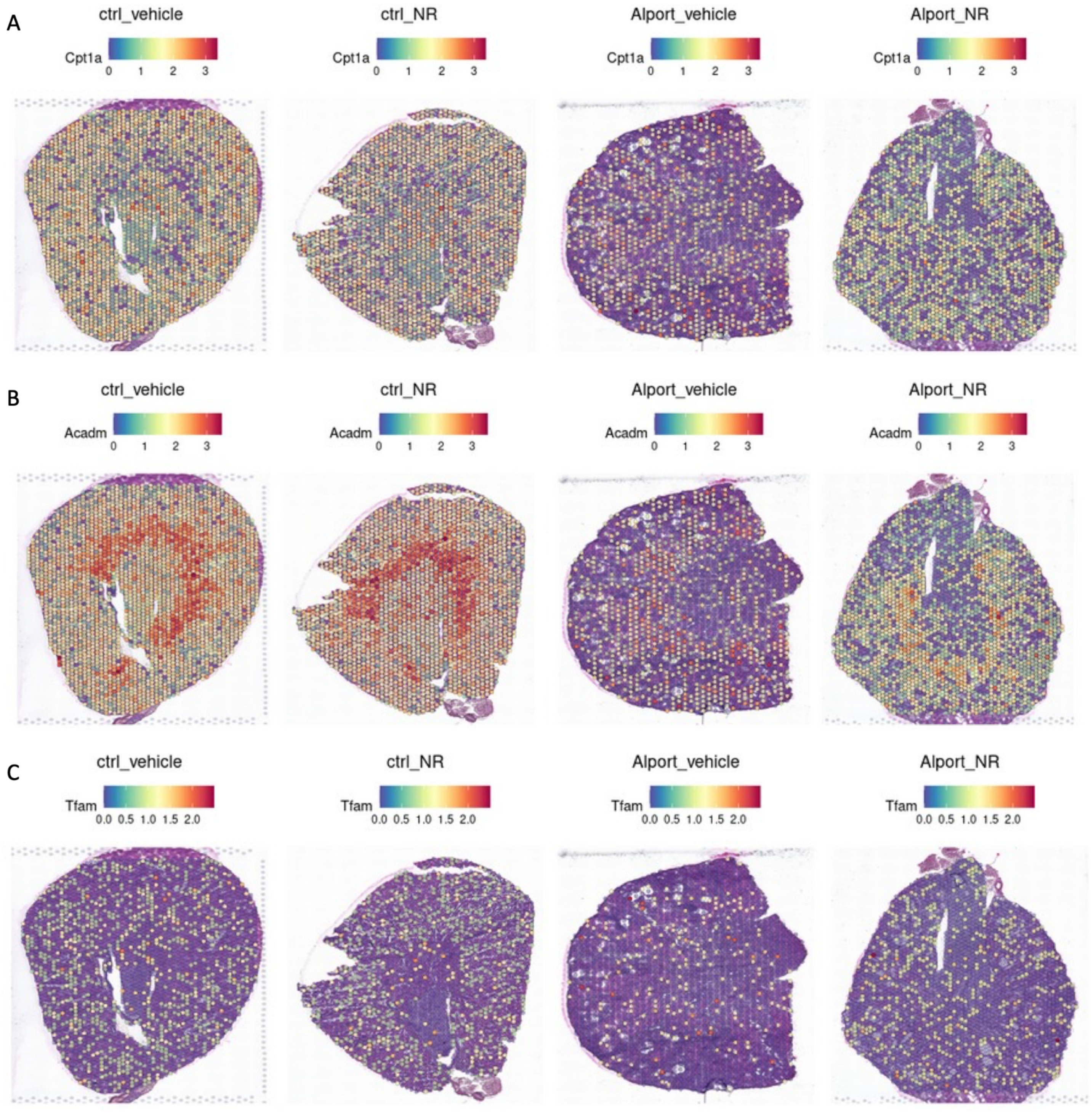

**Figure S31: Spatial localization of gene expression across 8,748 spots.** Visium spatial transcriptomics was performed on vehicle-treated male control mice (*far left*), NR-treated male control mice (*center left*), vehicle-treated male Alport mice (*center right*), and NR-treated male Alport mice (*far right*). (**A-C**) *Cpt1a* (*A*), *Acadm* (*B*), and *Tfam* (*C*) expression are reduced in Alport mice and restored with NR treatment. The ctrl_vehicle, Alport_vehicle, and Alport_NR panels of *Cpt1a* and *Acadm* are identical to those of **Fig. 8** and included here to enable comparison of all 4 conditions. *Acadm*, medium-chain acyl-coenzyme A dehydrogenase (MCAD); *Cpt1a*, carnitine palmitoyltransferase 1-α; Ctrl, control; NR, nicotinamide riboside; *Tfam*, mitochondrial transcription factor A.

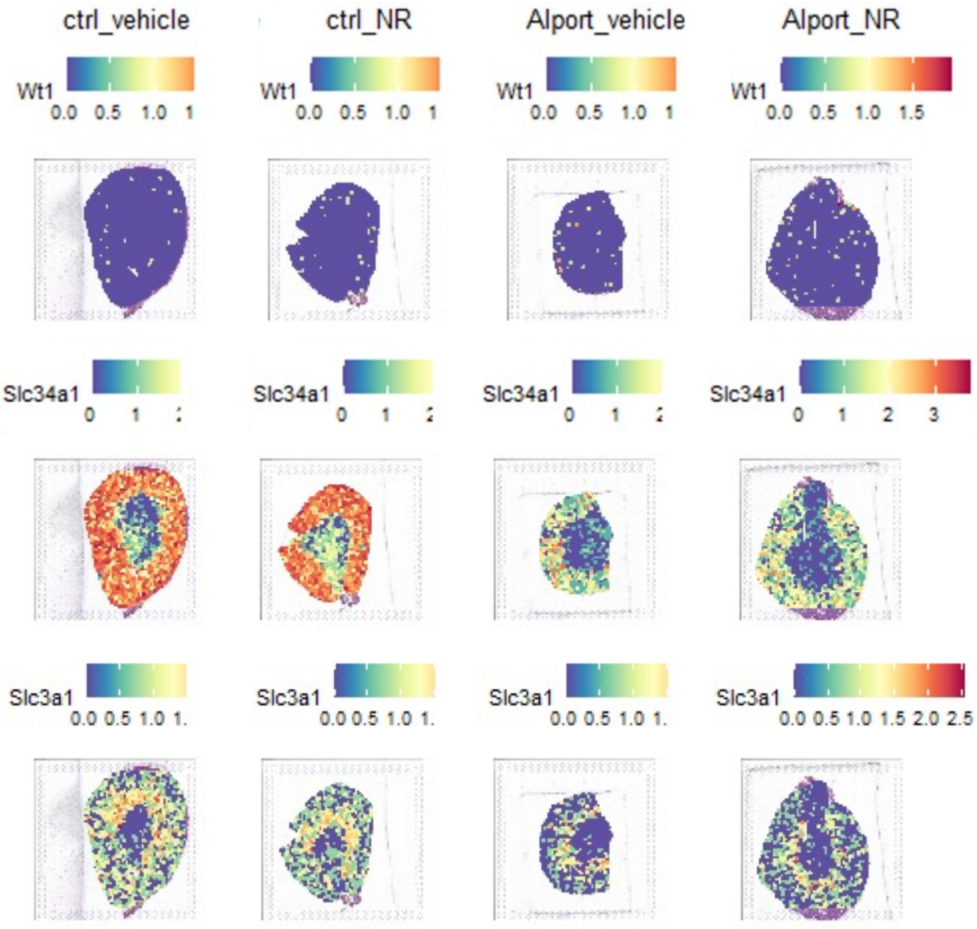

**Figure S32: Localization of the functional tissue units within the kidney.** Both histology and marker gene expression were used to define the functional tissue unit of most spots, including those in glomeruli (*Wt1* and *Nphs2*), the cortical tubulointerstitium (*Slc34a1*), and the outer medullary stripe (*Slc3a1*).

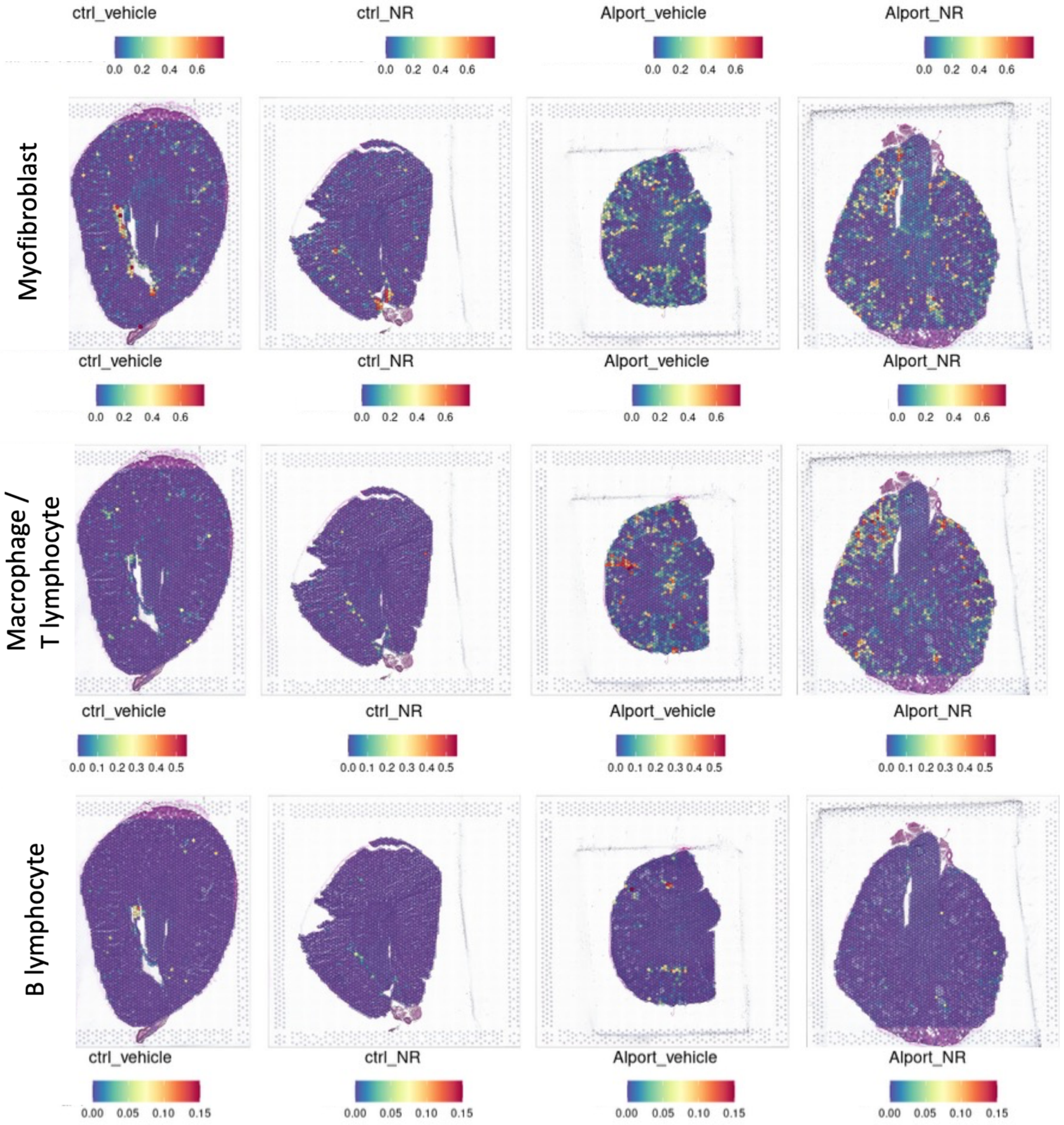

**Figure S33: NAD$^+$ supplementation reduces myofibroblast and immune cells on spatial transcriptomics.** Spatial localization of cell types across 8,748 spots. Labels were transferred from snRNA-seq to the spatial transcriptomics data to spatially localize the cell types based on gene expression profiles for the four male groups: control-vehicle (*far left*), control-NR (*center left*), Alport-vehicle (*center right*), and Alport-NR (*far right*). Myofibroblasts (stromal STR_3 cluster), macrophages, T lymphocytes, and B lymphocytes were increased in vehicle-treated Alport mice (versus vehicle-treated control mice). Consistent with the snRNA-seq, NR modestly reduced myofibroblast occurrence and markedly reduced immune cell infiltration in Alport mice.

**Table S1**: Alport mice do not have reduced cardiac function on echocardiography.

| | Control + Veh | Control + NR | Alport + Veh | Alport + NR |
|---|---|---|---|---|
| **Male (M-mode)** | *N* = 13 | *N* = 11 | *N* = 13 | *N* = 12 |
| Stroke Volume (µL) | 40.1 ± 2.5 | 44.3 ± 2.2 | 39.8 ± 2.3 | 42.1 ± 3.0 |
| Ejection Fraction (%) | 61.8 ± 2.3 | 65.3 ± 2.7 | 60.4 ± 2.8 | 63.3 ± 2.2 |
| Cardiac Output (mL/min) | 15.6 ± 1.5 | 16.4 ± 1.1 | 13.8 ± 1.8 | 15.3 ± 1.8 |
| LV Mass (mg) | 132.9 ± 5.1 | 141.2 ± 6.9 | 152.8 ± 12.7 | 158.0 ± 8.7 |
| **Male (B-mode)** | *N* = 13 | *N* = 11 | *N* = 13 | *N* = 12 |
| Stroke Volume (µL) | 29.5 ± 1.5 | 34.0 ± 2.1 | 30.3 ± 2.1 | 31.2 ± 2.0 |
| Ejection Fraction (%) | 52.8 ± 2.0 | 55.3 ± 2.4 | 51.5 ± 2.3 | 55.5 ± 2.0 |
| Cardiac Output (mL/min) | 12.6 ± 1.0 | 18.7 ± 2.4 | 14.7 ± 1.7 | 13.3 ± 1.9 |
| **Female (M-mode)** | *N* = 12 | *N* = 11 | *N* = 12 | *N* = 11 |
| Stroke Volume (µL) | 34.4 ± 1.9 | 40.7 ± 1.3 | 38.9 ± 2.4 | 40.6 ± 3.3 |
| Ejection Fraction (%) | 59.0 ± 2.8 | 62.3 ± 3.2 | 60.3 ± 2.5 | 63.6 ± 2.0 |
| Cardiac Output (mL/min) | 11.0 ± 1.1 | 17.5 ± 1.7 | 16.2 ± 2.1 | 13.6 ± 1.7 |
| LV Mass (mg) | 111.7 ± 6.6 | 111.4 ± 7.7 | 120.9 ± 5.9 | 119.3 ± 9.2 |
| **Female (B-mode)** | *N* = 12 | *N* = 11 | *N* = 12 | *N* = 11 |
| Stroke Volume (µL) | 25.6 ± 1.1 | 26.1 ± 2.0 | 31.0 ± 1.9 | 25.7 ± 2.7 |
| Ejection Fraction (%) | 53.3 ± 1.7 | 55.8 ± 3.1 | 54.3 ± 1.9 | 51.5 ± 3.4 |
| Cardiac Output (mL/min) | 11.1 ± 0.6 | 14.2 ± 1.9 | 13.9 ± 1.0 | 11.9 ± 2.5 |

Significance was determined by 1-way ANOVA with the Holm-Šídák correction for multiple comparisons. No comparisons were statistically significant. Data are expressed as the mean ± SEM. LV, left ventricle; *N*, number of mice; NR, nicotinamide riboside; Veh, vehicle.

**Table S2**: Alport mice do not have elevated blood pressure on photoplethysmography.

| | **Control + Veh** | **Control + NR** | **Alport + Veh** | **Alport + NR** |
|---|---|---|---|---|
| **Male** | *N* = 11 | *N* = 9 | *N* = 11 | *N* = 9 |
| Systolic BP (mmHg) | 106.3 ± 3.8 | 108.6 ± 4.4 | 107.4 ± 2.7 | 106.5 ± 4.4 |
| Diastolic BP (mmHg) | 56.3 ± 3.0 | 63.0 ± 4.3 | 53.1 ± 2.1 | 54.8 ± 4.2 |
| Mean BP (mmHg) | 73.1 ± 3.2 | 78.4 ± 4.2 | 71.3 ± 2.0 | 72.6 ± 4.1 |
| **Female** | *N* = 12 | *N* = 8 | *N* = 10 | *N* = 8 |
| Systolic BP (mmHg) | 105.0 ± 1.9 | 108.0 ± 3.3 | 105.1 ± 3.9 | 109.4 ± 4.5 |
| Diastolic BP (mmHg) | 56.7 ± 4.3 | 54.5 ± 2.6 | 55.3 ± 2.4 | 56.6 ± 3.9 |
| Mean BP (mmHg) | 73.2 ± 3.4 | 72.0 ± 2.9 | 72.4 ± 2.4 | 74.8 ± 3.4 |

Significance was determined by 1-way ANOVA with the Holm-Šídák correction for multiple comparisons. No comparisons were statistically significant. Data are expressed as the mean ± SEM. BP, blood pressure; *N*, number of mice; NR, nicotinamide riboside; Veh, vehicle.

**Table S3**: Plasma creatinine is unchanged at the 25-week timepoint.

| | **Control + Veh** | **Control + NR** | **Alport + Veh** | **Alport + NR** |
|---|---|---|---|---|
| **Male** | *N* = 13 | *N* = 12 | *N* = 10 | *N* = 11 |
| Plasma Creatinine (mg/dL) | 0.27 ± 0.02 | 0.27 ± 0.02 | 0.44 ± 0.10 | 0.34 ± 0.04 |
| **Female** | *N* = 12 | *N* = 13 | *N* = 12 | *N* = 11 |
| Plasma Creatinine (mg/dL) | 0.35 ± 0.03 | 0.40 ± 0.03 | 0.40 ± 0.02 | 0.41 ± 0.02 |

Significance was determined by 1-way ANOVA with the Holm-Šídák correction for multiple comparisons. No comparisons were statistically significant. Data are expressed as the mean ± SEM. *N*, number of mice; NR, nicotinamide riboside; Veh, vehicle.
